# Expanding the *TaNAS* gene family in bread wheat and exploring potential for intragenic biofortification

**DOI:** 10.64898/2025.12.07.692865

**Authors:** Oscar Carey-Fung, Jesse T. Beasley, Edbert Josua Felix, Lily Tarry-Smith, Nathan Leckie, Damien L. Callahan, Rudi Appels, Alexander A. T. Johnson

## Abstract

Higher plants utilise nicotianamine synthase (NAS) enzymes to produce nicotianamine (NA), a non-protein amino acid that chelates metals such as iron (Fe) and zinc (Zn) for long-distance transport. We identified 34 *TaNAS* genes in bread wheat (*Triticum aestivum* L., cv. Chinese Spring), and four additional cultivar-specific *TaNAS* genes, representing the largest *NAS* gene family identified to date. The expression of all *TaNAS* genes was highest in roots and upregulated (apart from the *TaNAS9* homoeologs) in response to hydroponic Fe deficiency. The TaNAS proteins ranged between 180 and 384 amino acids in length and showed variable N- and C-termini. To understand *TaNAS* function, we transformed bread wheat cv. Fielder to overexpress coding sequences from either *TaNAS1*, *TaNAS3*, *TaNAS4*, *TaNAS6*, or *TaNAS7*, and isolated three single locus homozygous events and null segregant (NS) controls for each *TaNAS* overexpression line for glasshouse and field evaluation. Under field conditions, grain Fe, Zn and NA concentrations were up to 1.6-fold, 1.8-fold and 3.7-fold higher, respectively, in *TaNAS6* events relative to NS without any negative impacts on agronomic traits. These results demonstrate the usefulness of available online genomic resources for analysing the *TaNAS* gene family in bread wheat and highlight intragenic strategies to enhance grain nutritional quality.

**Significance statement:** *Nicotianamine synthase* (*NAS*) genes play key roles in metal transport and homeostasis in plants. This study describes the identification of 34 *TaNAS* genes in bread wheat, representing a significant expansion on previous *TaNAS* studies, and demonstrates that constitutive overexpression of specific *TaNAS* genes could provide a novel intragenic approach for iron and zinc biofortification of bread wheat.

## Introduction

Nicotianamine (NA) is a non-protein amino acid, first purified from tobacco (*Nicotiana tabacum* L.), where it was shown to be critical for maintaining iron (Fe) homeostasis and in turn fundamental biological processes such as chlorophyll biosynthesis (Noma *et al*., 1971; Ripperger, 1986). Nicotianamine synthase (NAS), the enzyme responsible for catalysing the trimerisation of S-adenosylmethionine (SAM) into NA, was first identified in the *chloronerva* tomato mutant, which exhibits poor growth and leaf chlorosis due to a complete loss in the ability to synthesise NA (Higuchi *et al*., 1996; Ling *et al*., 1999). It is now widely understood that NA is ubiquitous to all higher plants and chelates a range of divalent metal cations including Fe^2+^, zinc (Zn^2+^), manganese (Mn^2+^) and nickel (Ni^2+^) for their long-distance transport between plant tissues (Von Wirén *et al*., 1999; Vacchina *et al*., 2003; Koike *et al*., 2004).

In graminaceous species, the *NAS* genes are classified into two clades. Clade I *NAS* genes are predominantly expressed in root tissues where NA facilitates long-distance transport of metals from source (i.e., roots and leaves) to sink (i.e., grain) tissues through the phloem, and uptake of Fe from the rhizosphere by serving as biosynthetic precursor to 2’-deoxymugineic acid (DMA) *via* nicotianamine aminotransferase (NAAT) and 2’-deoxymugineic acid synthase (DMAS) enzymes. In bread wheat, DMA is secreted into the rhizosphere by the transporter of mugineic acid 1 (TOM1), and Fe-DMA complexes are taken up into the roots by members of the yellow stripe-like (YSL) transporter family (Inoue *et al*., 2009; Nozoye *et al*., 2011). By contrast, Zn-DMA complexes are crucial for Zn translocation within the plant rather than Zn uptake from the soil (Suzuki *et al*., 2008). Soil Zn uptake is mediated mainly by Zn-regulated, iron-regulated transporter-like protein (ZIP) transporters such as the rice OsZIP9 transporter (Huang *et al*., 2020; Yang *et al*., 2020). Other OsZIP proteins play crucial roles in transporting Zn between rice tissues, and it is expected that the ZIP proteins in bread wheat play a similar role (Lee *et al*., 2010*a*,*b*; Sasaki *et al*., 2015; Li *et al*., 2021). In rice and *Arabidopsis halleri*, NA is proposed to form stable Zn-NA complexes in root cells where symplastic movement to the xylem allows for Zn-NA accumulation in the shoots and seeds (Lee *et al*., 2011; Deinlein *et al*., 2012). Clade II *NAS* genes are primarily expressed in shoot tissues and under conditions of Fe excess, where they maintain cellular Fe homeostasis in part by regulating short-distance cytoplasmic distribution (Aung *et al*., 2019; Seregin and Kozhevnikova, 2023). In the metal hyperaccumulator *Arabidopsis halleri*, NA accumulation and secretion confers tolerance to conditions of Zn excess (Tsednee *et al*., 2014). Within the cell, NA plays a critical role in the sequestration of excess metal ions into vacuoles and by preventing the formation of reactive oxygen species (ROS) (Pich *et al*., 2001). During the digestion of plant-based foods, NA plays a role in enhancing dietary Fe bioavailability and Caco-2 (human intestinal epithelial) cells show greater absorption of Fe bound to NA compared to other Fe-bioavailability enhancers (e.g., ascorbic acid, epicatechin) (Zheng *et al*., 2010; Lee *et al*., 2012; Eagling *et al*., 2014; Beasley *et al*., 2019*b*). Furthermore, rice (*Oryza sativa* L.) that contained high concentrations of NA reduced the symptoms of Fe-deficiency anaemia in mice more than conventional rice (Lee *et al*., 2012).

To date, overexpressing *NAS* genes has been a successful strategy for Fe and Zn biofortification of crops. Rice grain Fe, Zn, and NA concentrations were increased up to 3-, 2- , and 10.6-fold, respectively, following overexpression of *HvNAS1* from barley (*Hordeum vulgare* L.), 4-, 2-, and 9.3-fold, respectively, following overexpression of *OsNAS2* from rice, 2.9-, 2.2-, and 9.6-fold, respectively, following overexpression of *OsNAS3*, and 4-, 3.5-, and 50.6-fold, respectively following overexpression of both *OsNAS2* and *OsNAS3* (Masuda et al., 2009; Lee et al., 2009, Johnson et al., 2011, Lee et al., 2023). In wheat (*Triticum aestivum* L.), overexpressing *OsNAS2* increased grain Fe, Zn, and NA concentrations up to 1.5-, 1.6-, 15-fold, respectively, under glasshouse conditions and 1.3-, 1.4-, 2.9-fold, respectively, under field conditions (Beasley *et al*., 2019*a*). In tobacco, overexpressing *HvNAS1* increased seed Fe and Zn concentrations by 2.3-fold and 1.8-fold, respectively, and overexpressing the apple (*Malus xiaojinensis*) *MxNAS1* and *MxNAS2* genes increased leaf NA content ∼3.5-fold and improved plant tolerance to Fe deficiency (Takahashi et al., 2003; Han et al., 2013; Yang et al., 2015). Overexpressing the *Arabidopsis* (*Arabidopsis thaliana*) *AtNAS1* gene in potato (*Solanum tuberosum* L.) increased tuber Fe concentration by 2.4-fold (Zha *et al*., 2022).

The number of *NAS* genes present in plant genomes varies widely between rice (three *OsNAS* genes), maize (*Zea mays* L., nine *ZmNAS* genes), barley (seven *HvNAS* genes), and bread wheat (21 *TaNAS* genes) (Higuchi *et al*., 1999; Inoue *et al*., 2003; Zhou *et al*., 2013; Bonneau *et al*., 2016). Here we describe the identification of 34 *TaNAS* genes in the recently annotated cv. Chinese Spring bread wheat genome (IWGSC Refseq 1.1 and 2.1) and the 10+ Wheat Genome Project, representing a significant expansion of the *TaNAS* gene family (Walkowiak *et al*., 2020; Zhu *et al*., 2021). Furthermore, we demonstrate that constitutively overexpressing six unique *TaNAS* genes provides a route for the intragenic biofortification of bread wheat.

## Materials & methods

### Identification and annotation of TaNAS genes

Previously annotated *NAS* coding sequences (CDSs) from *Arabidopsis thaliana*, *Hordeum vulgare* L., *Oryza sativa* L., *Triticum aestivum* L., and *Zea mays* L. were used as queries to blast the IWGSC Refseq1.1 and IWGSC Refseq 2.1 in EnsemblPlants (https://plants.ensembl.org/index.html), leading to the identification of 43 potential *TaNAS* genes in cv. Chinese Spring (Table S1, S5, S6) (Bolser *et al*., 2015; Howe *et al*., 2020). Additional *TaNAS* genes in the 10+ Wheat Genome Project were identified in cultivars ArinaLrFor, Julius, Lancer, Norin61, SY Mattis, Mace, Jagger, Landmark, and Stanley and in *Triticum spelta* L. by blasting the *TaNAS* CDSs from cv. Chinese Spring against each of these genomes on EnsemblPlants (Appels *et al*., 2018; Zhu *et al*., 2021). All *TaNAS* genes in cv. Fielder were identified using Earlham Institute’s Grassroots Infrastructure BLAST tool (https://grassroots.tools/service/blast-blastn) (Bian *et al*., 2017). The *TaNAS4-D1* and *TaNAS4-D2* homoeologs in cultivars ArinaLrFor, Julius, Lancer, Norin61, SY Mattis, Mace, Stanley, and Fielder and *TaNAS7-A1* homoeolog in cv. Jagger could not be identified on a chromosome. The cv. Chinese Spring *TaNAS* CDSs were annotated following visualisation of the *TaNAS* genes in the Apollo database (https://apollo-portal.genome.edu.au/), which integrates the Refseq2.1 reference genome sequence, models from NCBI and RNA-seq data (Dunn *et al*., 2019; Walkowiak *et al*., 2020; Zhu *et al*., 2021). Only the CDS of each *TaNAS* gene was annotated, as the 5’ and 3’ UTR annotations were often inconsistent between the Refseq1.1, Refseq2.1, and NCBI models, and did not necessarily match the RNA-seq read expression profiles. Any discrepancies between Refseq1.1, Refseq2.1, and NCBI annotations in the CDS of *TaNAS* genes were resolved using the RNA-seq datasets integrated into Apollo and the RNA-seq dataset generated in this study (Table S2, Fig. S2B). Additional Ribo-seq data to resolve the presence of a putative N-terminus extension in TaNAS6-D1 was sourced from Guo et al., (2023) (Fig. S2A). The Phyre2 (v2.0) protein fold recognition software was used to predict the alignment of *TaNAS* proteins with the c3fpjA_ template, corresponding to alignment with the 3D structure of an archaeal NAS protein from *Methanothermobacter thermautotrophicus* in the Protein Data Bank (PDB) (Dreyfus *et al*., 2009; Kelley *et al*., 2015). Nine potential *TaNAS* genes in cv. Chinese Spring were disqualified as they either did not contain the c3pjA_ protein structure and/or did not respond to Fe deficiency (Table S2, Fig. 3). Bread wheat, einkorn wheat, barley, maize and rice *NAS* CDSs were aligned using the Geneious alignment tool (v2024.0.7), and phylogenetic construction was performed using the PhyML Geneious plugin with the K80: Kimura substitution model and 1000 bootstrap replications (Guindon *et al*., 2010). Protein sequences in Fig. 2B were aligned using the Geneious alignment tool (v2024.0.7).

**Figure 1.**
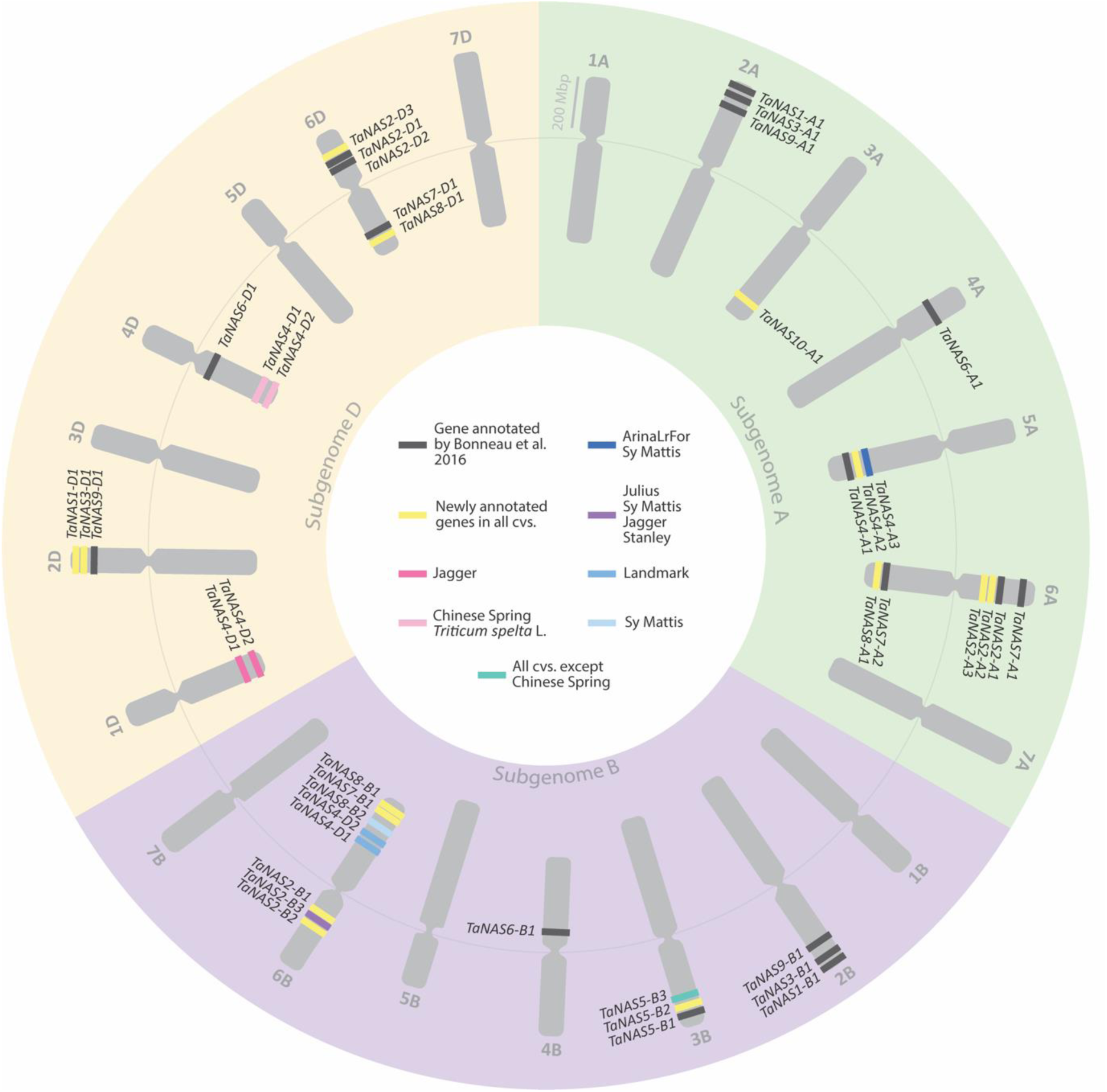
The location of *TaNAS* genes across 11 bread wheat (*Triticum aestivum* L.) cultivars and *Triticum spelta* L. Chromosomes within the A (green), B (purple), and D (orange) subgenomes of bread wheat are depicted clockwise. Bread wheat *TaNAS* genes that were previously annotated in Bonneau et al., (2016) (black), newly annotated in all cultivars (yellow); or exclusively present in cv. Jagger (dark pink); cv. Chinese Spring and cv. *Triticum spelta* L. (light pink); cv. ArinaLrFor and cv. SY Mattis (dark blue); cv. Julius, cv. SY Mattis, cv. Jagger, and cv. Stanley (purple); cv. Landmark (blue); cv. SY Mattis (light blue); or all cultivars except for cv. Chinese Spring (teal) are indicated in approximate genomic regions of each chromosome.

**Figure 2.**
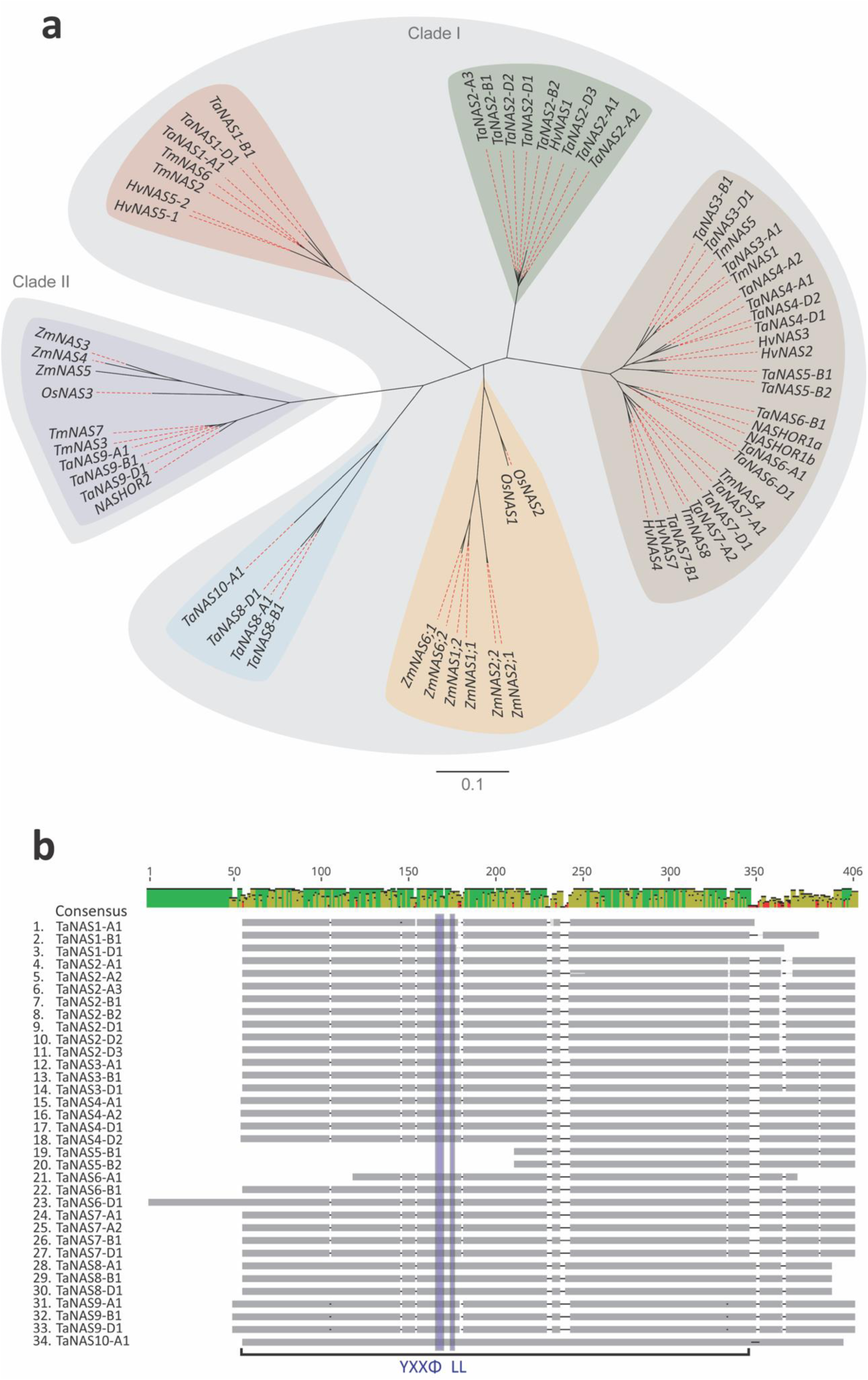
Phylogenetic analysis of *TaNAS* genes and TaNAS protein alignment. (A) Unrooted phylogenetic tree of *NAS* coding sequences in bread wheat (*Triticum aestivum* L., *Ta*), einkorn wheat (*Triticum monococcum* L., *Tm*), barley (*Hordeum vulgare* L., *Hv*), maize (*Zea mays* L., *Zm*), and rice (*Oryza sativa* L., *Os*). The scale bar represents evolutionary distance in substitutions per site. (B) Protein alignment of the 34 TaNAS proteins in cv. Chinese Spring. The top identity bar indicates 100% consensus amongst aligned sequences (green bars), medium consensus (yellow bars), and low consensus (red bars). Deletions in the majority (>50%) or minority (<50%) of sequence variations are indicated (black lines and grey lines, respectively). The blue transparent rectangles highlight the location of the conserved YXXФ and LL motifs and the black bracket indicates the conserved ‘core region’ within the TaNAS proteins.

**Figure 3.**
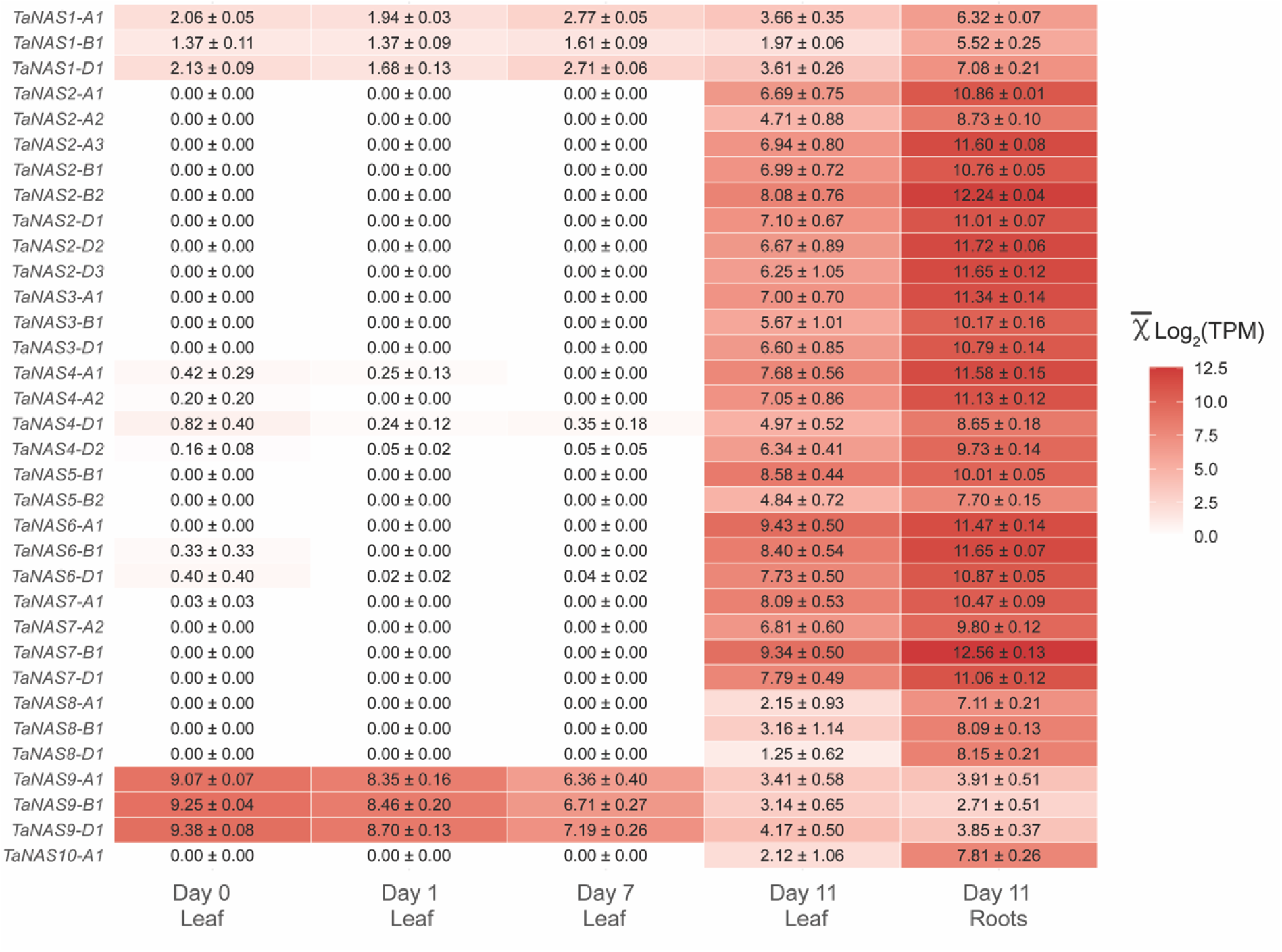
Expression of 34 bread wheat *TaNAS* genes in cv. Fielder under hydroponic Fe sufficient and deficient conditions. The expression of 34 *TaNAS* genes in hydroponically grown plants at day 0 (50 µM Fe), 1, 7, and 11 of Fe deficiency (0 µM Fe) in leaf tissues, and day 11 of Fe deficiency in root tissues. Values represent the average of three log transformed transcripts per million (TPM) values and the standard error of the mean for each gene (n=3). Genes with no detected expression and high expression are indicated (white and red, respectively).

### Bread wheat transformation

Each *TaNAS* CDS was amplified from bread wheat genomic DNA (cv. Chinese Spring) using Phusion™ high-fidelity Taq polymerase (Thermo Fisher Scientific, Waltham, Massachusetts, USA), and cloned into the pENTR vector (Karimi *et al*., 2007). The pENTR vector containing a *TaNAS* CDS was digested with AseI (New England Biolabs, Ipswich, Massachusetts, USA) and recombined (LR reaction) into a modified pMDC32 vector containing a maize ubiquitin 1 promoter to drive expression of the *TaNAS* CDS and hygromycin resistance (*hptII*) gene for bread wheat transformation selection (Curtis and Grossniklaus, 2003). Bread wheat transformation was performed at the University of Adelaide using the *Agrobacterium tumefaciens* strain AGL1 as described (Okada *et al*., 2019). Briefly, immature bread wheat embryos were harvested from cv. Fielder plants 14 days post anthesis and co-cultured with Agrobacterium containing the pMDC32 vector. Embryonic axes from the embryos were removed following co-culture. Resting, callus induction, selection and regeneration were performed over 10 weeks prior to soil transfer in a glasshouse. Each plantlet derived from callus tissue was considered an individual event. Harvested T_1_ seeds from mature plants of each event were sent to the University of Melbourne.

### Isolation of single-locus homozygous lines

A total of 16 T_1_ grain from 10-15 events from each *TaNAS*-overexpression (OE) construct were stratified (4°C for two days) and germinated in 56-well trays containing sifted potting mix (pine bark, peat, and sand mixture) with Osmocote® fertiliser (4.5 g/L). Approximately one cm of leaf was sampled and placed in a 96-well plate. Genomic DNA was extracted by adding 50 µL of Extract-N-Amp extraction solution (Merck, Darmstadt, Germany), incubating the samples at 95°C for ten minutes and adding 50 µL of Extract-N-Amp dilution solution. Melt curve analysis was performed using 1 µL of DNA template, 5 µL of KAPA SYBR® FAST (Merck, Darmstadt, Germany), 0.2 µL of primer (10 µM), and 10 µL of nuclease free H_2_O. Forward (5’CTCGGAGGGCGAAGAATCTC) and reverse (5’GCGGGAGATGCAATAGGTCA) primers were used to amplify *hptII* in the T-DNA and forward (5’CAAGCCGCTGCACTACAAGG) and reverse (5’AGGGGACGGTGCAGATGAA) primers were used to amplify *TaCyclophilin* (housekeeping gene) as a reaction control. PCR conditions included initial denaturation at 95°C for 3 minutes, and 35 cycles of: 1. denaturation at 95°C for 20 seconds, 2. primer annealing at 63°C for 20 seconds, and 3. fluorescence measurement by plate reading. A melt curve was generated by incubating the samples from 78°C to 90°C in 0.1°C increments for five seconds at each increment. For T_1_ populations with a transgene negative:positive ratio of 1:3, the event was determined to be a single locus event and two null segregants (NSs) and six transgenics were grown to maturity. Sixteen T_2_ progeny grain were placed on moist filter paper for 2 days and transferred into wells of a 96 well plate where the grain were genotyped by the method described above. A T_1_ parent was determined to be homozygous if all T_2_ progeny were transgenic positive. A T_1_ NS sibling from each single locus event was also advanced for glasshouse and field experiments to control for the effects of tissue culture (Miyao *et al*., 2012).

### Glasshouse, speed breeding, and field growth conditions

Glasshouse growth of T_1_, T_3_, and T_4_ plants and speed breeding of T_2_ plants was performed at Parkville Campus at The University of Melbourne, Victoria, Australia. Either 1 L (glasshouse) or 8.1 L (speed breeding) pots containing general potting mix with Osmocote® fertiliser (4.5 g/L) were used for plant growth. Plants were watered using drip irrigation (∼50 mL) once per day. Approximately 10 mL of ammonium nitrate (NH₄NO₃ solution, 28.57 g/L) fertiliser was added to the glasshouse plants 6 weeks and 9 weeks after sowing. In the glasshouse plants were sown in May/June and harvested in December/January. Speed bred plants were grown under a 22-hour day (22°C) and 2-hour night (17°C) cycle with 65% humidity and Heliospectra LED Grow Lights programmed to 400-600 photosynthetic photon flux densities at pot height (Heliospextra, Gothenburg, Sweden) (Watson *et al*., 2018). Field experiments were performed at Dookie Campus at The University of Melbourne, Victoria, Australia. Each plot comprised of six furrows (75 cm long) spaced 25 cm apart and grain was sown at a density of 6 grams per plot (2022) or 10 grams per plot (2023). At sowing, plots were fertilised with Di-Ammonium Phosphate (DAP) at a rate of 100 kg/Ha, and urea was applied at a rate of 150 kg/Ha during tillering and stem elongation stages. Plants relied on rainfall throughout the growing seasons. Agronomic traits of field grown plants were determined by harvesting and phenotyping one representative row and multiplying by 6 to calculate biomass, grain weight or tiller number per m^2^.

### RNA-sequencing of hydroponically grown plants

Grain harvested from cv. Fielder plants were stratified at 4°C for two days and then surface sterilised with VitaVax 200FF (Hewitt & Whitty, Geelong, Victoria, Australia) and placed on sterile moist filter paper at 20°C for two days. Germinated seedlings were transferred to hydroponic boxes containing a nutrient replete solution: 2 mM NH_4_NO_3_, 5 mM KNO_3_, 2 mM Ca(NO_3_)_2_·4H_2_O, 2mM MgSO_4_·7H_2_O, 0.1 mM KH_2_PO_4_, 10 µM H_3_BO_3_, 5 µM MnCl_2_·4H_2_O, 5 µM ZnSO_4_·7H_2_O, 0.5 µM CuSO_4_·5H_2_O, 0.1 µM Na_2_MoO_4_·2H_2_O, and 50 µM NaFe^3+^EDTA.

Following 12 days of nutrient replete growth, the solution was replaced with an Fe deficient solution (0 µM NaFe^3+^EDTA) and grown for a further 11 days. The nutrient solution was replaced every four days. Leaf tissues were harvested at days 0, 1, 7 and 11 of Fe deficiency and root tissues were harvested at day 11 of Fe deficiency. All tissues were immediately snap frozen in liquid nitrogen following sampling. Approximately 50 mg of tissue in TRI Reagent® (Merck, Darmstadt, Germany) was homogenised with stainless steel beads and a TissueLyser II (Qiagen, Hilden, Germany). The Direct-zol RNA Miniprep Kit (Zymo Research, Irvine, California, USA) was used to extract RNA according to the manufacturer’s instructions. Quality control and RNA-sequencing was performed by the Australian Genome Research Facility (AGRF, Melbourne, Victoria, Australia). Briefly, sample quality was assessed using the Agilent 4200 TapeStation and Agilent RNA ScreenTape (Agilent Technologies, Santa Clara, California, USA) and reagents according to the manufacturer’s instructions. Sample concentrations were measured using the QuantiFluor RNA Kit (Promega, Madison, Wisconsin, USA). Library preparation was performed using the Illumina Stranded mRNA library preparation workflow and sequencing was completed with the Illumina NovaSeq X Plus with 150 paired end reads and one lane of a 10B-300 flow cell according to the manufacturer’s instructions (Illumina, San Diego, California, USA).

### Gene expression analysis

Quantitative reverse transcription PCR analysis of each *TaNAS*-OE transgene was performed on five-day old seedlings. Shoot tissue above the crown from three plants per *TaNAS* event was individually harvested and snap frozen in liquid nitrogen and homogenized in TRIzol (Thermo Fisher, Carlsbad, California, USA). Total RNA was extracted using the Direct-zol RNA Miniprep Kit (Zymo Research, Irvine, California, USA), according to the manufacturer’s instructions. All RNA samples were normalised to 83 ng/µL and genomic DNA was removed using the RQ1 RNase-Free DNAse kit (Promega, Madison, Wisconsin, USA), according to the manufacturer’s instructions. Reverse transcription was performed using the Tetro cDNA Synthesis kit (Bioline, Meridian Bioscience, Memphis, Tennessee, USA), according to the manufacturer’s instructions. Gene expression was quantified using KAPA SYBR^®^ FAST qPCR Master Mix (2X) Kit (Sigma Aldrich, St. Louis, Missouri, USA) with three technical replicates per biological replicate. Expression of the *TaNAS*-OE gene was determined using primers within the attB2 cloning site and 5’ end of the nosT (5’ CCGACCCAGCTTTCTTGTAC and 5’ TGTTTGAACGATCGGGGAAA). Expression of the *TaNAS*-OE transgene was normalised against the geometric mean of three housekeeping genes: *TaACT* (5’ GACAATGGAACCGGAATGGTC and 5’ GTGTGATGCCAGATTTTCTCCAT), *TaGAPDH* (5’ TTCAACATCATTCCAAGCACCA and 5’ CGTAACCCAAAATGCCCTTG), and *TaSAR* (5’ GAGTCTGCCCACCCATTCGTAA and 5’ GACATGCCATAGGTTTCAGCGAC). Analysis of Ct values was performed using the 2^-ΔΔCt^ method and normalised against *TaNAS1-1* (Livak and Schmittgen, 2001).

Analysis of RNA-sequencing data was performed in Galaxy Australia (v1.0.0). Paired-end files from each sample and lane were concatenated (Galaxy v1.0.0) to generate single files for each read pair (R1 and R2). Quality checks were performed using FastQC (v0.74+galaxy1) and summarised with MultiQC (v1.11+galaxy1). Adapter trimming and low-quality base removal were done with Trim Galore (v0.6.7+galaxy0), applying a Phred score threshold of 10 and discarding reads under 30 base pairs (bp). Post-trimming quality was reassessed with FastQC. Reads were aligned to the “161010_Chinese_Spring_v1.0_pseudomolecules.fasta” reference genome (https://urgi.versailles.inra.fr/) using HISAT2 (v2.2.1+galaxy1), with paired-end and reverse strand (RF) options. Transcripts were assembled with StringTie (v2.2.3+galaxy0) using the HISAT2 BAM output and the reference high confidence GTF file (IWGSC_v1.1_HC_20170706.gtf) and also the low confidence GTF file (IWGSC_v1.1_LC_20170706.gtf) to capture all low confidence *TaNAS* genes (https://urgi.versailles.inra.fr/). Gene counts were quantified with FeatureCounts (v2.0.3+galaxy2), creating a count and gene length matrix for downstream expression analysis. Count and length matrices were loaded into Rstudio (v4.4.1) to calculate Transcripts Per Million (TPM) for each sample. Calculations followed standard protocols using the dplyr package (v1.1.4), where raw counts were divided by gene lengths in kilobases to generate reads per kilobase (RPK). The total RPK was summed to create a scaling factor for each sample. TPM values were then obtained by dividing RPK values by this scaling factor and multiplying by 10^6^, ensuring comparability across samples. This approach was applied to three biological replicates across five different conditions for both high-confidence (HC) and low-confidence (LC) annotations.

### ICP-OES analysis

Harvested spikes from the glasshouse and field experiments were dried in an oven (30°C) for at least two days. Grain samples (∼3 g) from each plant were ground (60 seconds, 25000 rpm) into wholegrain flour using a tube-mill (Tube Mill Control, IKA, Staufen, Germany). Wholegrain flour mineral concentrations were analysed at The University of Melbourne’s Trace Analysis for Chemical, Earth and Environmental Sciences (TrACEES) platform as previously described (Wheal *et al*., 2011). Flour samples (0.225-0.250 g) were digested in batches (n=40) using either nitric acid (2 mL, 69%)/hydrogen peroxide (0.5 mL, 30%) or nitric acid (7 mL, 69%)/hydrogen peroxide (1 mL, 30%) solutions in a heat block for 2 hrs at 125°C or a microwave digester (ETHOS™ EASY, Metrohm Australia, Mitcham, VIC, Aus) for 30 mins at 200°C, respectively. Samples had a final nitric acid concentration of 9%. Analysis was performed on a Perkin Elmer 8300 DV ICP-OES (Syngystix v3.0) relative to a 9% nitric acid sample matrix and internal standards for each element. Random samples were selected for repeat analysis throughout the run as quality controls.

### NA and DMA analysis

Grain nicotianamine (NA) and 2’-deoxymugineic acid (DMA) quantification was performed using reverse phase liquid-chromatography (Selby-Pham *et al*., 2017; Beasley *et al*., 2019*a*, 2022). Briefly, 25 mg of ground wholegrain flour was combined with 500 µL of methanol (Sigma Aldrich, St. Louis, Missouri, USA) in a 2 mL tube and incubated in a thermoshaker (TS-100 Thermoshaker, Biosan, Riga, Latvia) at 30°C and 1400 RPM for 15 minutes. The tubes were centrifuged (13300 rpm, 10 mins), and the supernatant was transferred into a new 2 mL tube. The original pellet was resuspended in 500 µL of MilliQ H_2_O *via* vortexing and centrifuged (13300 rpm, 10 mins). The resulting supernatant was combined with the methanol supernatant from the previous step. The combined extract was centrifuged prior to storage at -20°C. After thawing, the extract was centrifuged and an aliquot (5 µL) was combined with 10 µL of sodium borate buffer (1 M, (Merck, Darmstadt, Germany), 10 µL of EDTA (50 mM, (Merck, Darmstadt, Germany), and 40 µL of Fluorenylmethyloxycarbonyl-chloride (FMOC-Cl, 50 mM, Merck, Darmstadt, Germany), and incubated in a thermoshaker at 60 °C and 700 RPM for 15 minutes. The derivatised product was centrifuged and combined with 8.9 µL of formic acid (5%, Tokyo Chemical Industry, Chuo City, Tokyo, Japan) and vortexed to quench the reaction. Derivatised samples analysed by LC-MS using a Zorbax Eclipse XDB-C18 Rapid Resolution HS 2.1x100 mm, 1.8 µM particle size column (Agilent Technologies Inc., Santa Clara, CA, USA) and a triple Quadrupole LC-MS system (Shimadzu-8050) with electrospray ionisation in positive ion MRM mode. A binary mobile phase system was used for separation with 0.1% formic acid in MilliQ H_2_O (aqueous) and 0.1% formic acid in 100% acetonitrile (organic) mobile phases (Merck, Darmstadt, Germany). Quantification was carried out using an external calibration curve with standards from 0.075 to 75 µM prepared from 750 µM aqueous NA and DMA stocks (Toronto Research Chemicals, Toronto, ON, Canada). Peak area values were calculated using LabsSolutions Insight software (https://www.shimadzu.com/) and Microsoft Excel 170 (https://www.microsoft.com/en-us/microsoft-365/excel). Pooled biological quality control samples and blanks were run, and the system re-calibrated every 100 samples. Samples were randomised prior to running.

## Results

### Identification of 34 TaNAS genes in bread wheat across different chromosomes and cultivars

Thirty-four *TaNAS* genes were identified in the bread wheat genome (cv. Chinese Spring), including 14 that were previously uncharacterised (Fig. 1 and Table S1). The average CDS identity for the 34 *TaNAS* genes was 78.8% and ranged from 61.8 to 100%. The *TaNAS4-D1* and *TaNAS4-D2* homoeologs have the same CDS, possibly due to a recent intrachromosomal gene duplication event. All 34 *TaNAS* genes were present in the genomes of cultivars ArinaLrFor, Julius, Lancer, Norin61, SY Mattis, Mace, Jagger, Landmark, Stanley, and Fielder and 33 *TaNAS* genes (all apart from *TaNAS7-A1*) were present in the genome of *Triticum spelta* L. Four additional *TaNAS* genes (*TaNAS2-B3*, *TaNAS4-A3*, *TaNAS5-B3*, and *TaNAS8-B2*) were found in the cultivars analysed but not in cv. Chinese Spring (Table S3). The genomic location and presence of the *TaNAS* genes varied across cultivars (Fig. 1). Cultivars ArinaLrFor and SY Mattis contained an additional *TaNAS* gene on chromosome 5A (*TaNAS4-A3*), which shared over 95% CDS identity with the *TaNAS4-A1, TaNAS4-A2*, *TaNAS4-D1,* and *TaNAS4-D2* homoeologs (Fig. 1, Table S3). Cultivars Julius, SY Mattis, Jagger and Stanley contained an additional *TaNAS* gene on chromosome 6B (*TaNAS2-B3*), which shared over 95% CDS identity with the *TaNAS2-A1*, *TaNAS2-A2*, *TaNAS2-A3*, *TaNAS2-B1*, *TaNAS2-B2*, *TaNAS2-D1*, *TaNAS2-D2*, and *TaNAS2-D3* homoeologs. Cultivar SY Mattis contained an additional *TaNAS* gene on chromosome 6B (*TaNAS8-B2*), which shared over 95.4% CDS identity to the *TaNAS8-A*, *TaNAS8-B*, and *TaNAS8-D* homoeologs. All cultivars apart from cv. Chinese Spring contained an additional *TaNAS* gene on chromosome 3B (*TaNAS5-B3*), which shared over 90.6% CDS identity to the *TaNAS5-B1* and *TaNAS5-B2* homoeologs. All 425 *TaNAS* CDSs from cvs. Chinese Spring, ArinaLrFor, Julius, Lancer, Norin61, SY Mattis, Mace, Jagger, Landmark, Stanley, Fielder, and *Triticum spelta* L. were aligned (Fig. S1), to assess gene and protein variation between cultivars (Table S3). Identity analysis using PhyML in Geneious clustered each *TaNAS1*, *TaNAS2*, *TaNAS3*, *TaNAS4*, *TaNAS5*, *TaNAS6*, *TaNAS7*, *TaNAS8*, *TaNAS9*, and *TaNAS10* group together (Fig. S1). Slight sequence variation between cultivars was observed, with up to seven haplotypes present in some homoeologs (Table S3). The *TaNAS2-A1*, *TaNAS2-A3*, *TaNAS2-D1*, *TaNAS2-D2*, *TaNAS2-D3*, *TaNAS3-B1*, and *TaNAS6-B1* genes shared 100% CDS identity between all cultivars.

### All TaNAS genes encode proteins that are conserved but vary in length

Phylogenetic analysis showed that the cv. Chinese Spring *TaNAS* genes are highly conserved amongst monocots and cluster into two clades (Fig. 2A). Clade II is comprised of *OsNAS3*, *ZmNAS3*, *ZmNAS4*, *ZmNAS5*, *TmNAS7*, *TmNAS3*, *NASHOR2*, *TaNAS9-A1*, *TaNAS9-B1*, and *TaNAS9-D1*. Clade I is comprised of the remaining *NAS* genes and clusters into five sub-groups. All *TaNAS* proteins were predicted to align with the c3fpjA_ template of an archaeal NAS protein from *Methanothermobacter thermautotrophicus* with 100% confidence (Table S2). The TaNAS proteins shared a conserved region that is between 126 to 284 amino acids (aa) long (the ‘core region’), and for 19 TaNAS proteins this core region was 278 aa long. The average identity of the core region was 77.3% (52.5 - 100%). Outside the core region, the C-terminus region ranged between 3 to 57 aa long and had an average identity of 31.4% (0 - 100%) (Fig. 2B). The TaNAS1 proteins have the shortest C-termini amongst all bread wheat TaNAS proteins (18 aa in length on average) (Fig. 2B). Some cultivars are predicted to have an extended N-terminus in TaNAS6-A1 (cvs. SY Mattis, Mace, Norin61, Jagger, Stanley, ArinaLrFor, Landmark, Julius, and Lancer) and TaNAS6-D1 (cvs. Chinese Spring and *Triticum spelta* L.). To determine whether a similar extended N-terminus was present in TaNAS6-D1 of cv. Fielder, we conducted RNA-seq under Fe sufficient and deficient conditions (Fig. S2B). Coverage of RNA-seq data was only observed downstream of the predicted extended N-terminus translation start site and is only expressed under Fe deficiency, suggesting that the extended TaNAS6-D1 N-terminus is not translated in Fielder. To verify IWGCS Refseq1.0 and Refseq2.1 annotations that an extended N-terminus is present in TaNAS6-D1 of cv. Chinese Spring, we sourced Ribo-seq data from a study that performed ribosomal and RNA sequencing of developing cv. Chinese Spring wheat grain (Guo *et al*., 2023). Using this data, we observed that RNA-Seq and ribosomal coverage was only present downstream of the predicted extended N-terminus, suggesting the extended TaNAS6-D1 N-terminus is also not translated in cv. Chinese Spring developing grain (Fig. S2A). All TaNAS proteins contained the conserved LL motif and tyrosine-based sorting motif (YXXΦ), where X is any amino acid and Φ is an amino acid with a bulky hydrophobic side chains, apart from *TaNAS5-B2* and *TaNAS5-B3* (Fig. 2B). Additionally, in cv. Chinese Spring, a single nucleotide polymorphism causes truncation of the *TaNAS5-B1* protein, resulting in the loss of both the YXXΦ and LL motifs. The full-length *TaNAS5-B1* is present in all other cultivars (Table S3).

### The 34 TaNAS genes are mostly expressed in roots and are regulated by environmental Fe conditions

RNA-seq analysis on hydroponically grown cv. Fielder plants revealed that the expression (measured in log transformed transcripts per million) of most *TaNAS2*, *TaNAS3*, *TaNAS4*, *TaNAS5*, *TaNAS6*, *TaNAS7*, *TaNAS8* and *TaNAS10* genes was undetectable in leaf tissue under Fe sufficient conditions, and strongly upregulated by day 11 of an Fe deficiency treatment (Fig. 3). By contrast, the *TaNAS1* genes were moderately expressed in leaf tissue under Fe sufficient conditions and were upregulated between 1.4- to 1.8-fold at day 11 of Fe deficiency. Of all the *TaNAS* genes, the *TaNAS9* homoeologs showed the highest expression in leaf tissue under Fe sufficient conditions and the lowest expression in root tissue under Fe deficient conditions, and the expression of these genes was downregulated between 2.3- to 2.9-fold in the leaves at day 11 of Fe deficiency. The expression of all *TaNAS* genes was between 1.2- to 6.5-fold higher in roots compared to leaves at day 11 of Fe deficiency, excluding the *TaNAS9* genes. From RNA-seq datasets integrated into the Apollo database, most *TaNAS* genes showed high expression in wheat root tissues under Fe sufficient conditions, apart from *TaNAS7-A2* and *TaNAS10-A1* which showed low to moderate expression in all tissues and the *TaNAS9* genes which showed low expression in root tissues and high expression in leaf/stem tissues (Table S2).

### Generation of six single-locus transgenic bread wheat lines overexpressing different versions of TaNAS genes

Bread wheat cultivar (cv.) Fielder transformants constitutively overexpressing (OE) a *TaNAS* gene (*TaNAS*-OE) under regulatory control of the maize ubiquitin (UBI-1) promoter were generated by using the CDS of the following *TaNAS* genes: *TaNAS1-A1* (with transformed plants hereafter referred to as *TaNAS1*); *TaNAS3-B1* (hereafter referred to as *TaNAS3*); a chimera of the *TaNAS4-A1* N-terminus and the *TaNAS4-A2* C-terminus (hereafter referred to as *TaNAS4*); *TaNAS6-D1* (hereafter referred to as *TaNAS6*); a truncated version of *TaNAS6-D1* without the extended N-terminus (hereafter referred to as *TaNAS6S*); and *TaNAS7-A1* (hereafter referred to as *TaNAS7*) (Fig. 4A). Three T_1_ homozygous single-locus events (termed 1, 2, and 3) and their respective null segregant (NS) controls were isolated for each *TaNAS* construct, using a high-throughput melt-curve approach to determine segregation ratios and zygosity (Fig. 4B, C). Three events per construct were analysed to account for variation in transgene expression due to differences in site-specific integration. Transgene expression analysis revealed some variation between *TaNAS*-OE events; however, the differences were not statistically significant (Fig. 4D). Transgene expression was detected in all events apart from *TaNAS7-1*-OE and this event was therefore excluded from all subsequent analyses.

**Figure 4.**
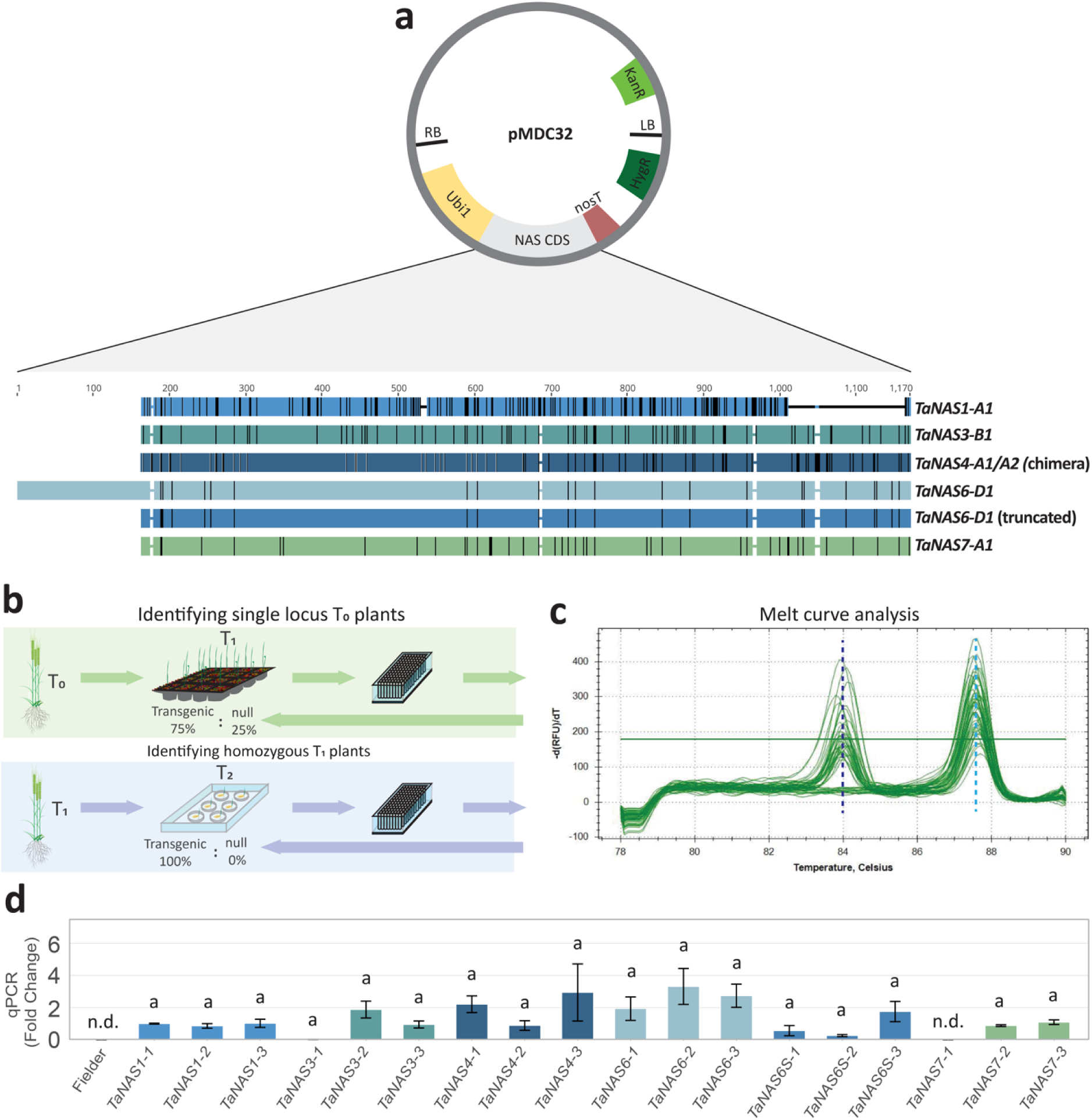
Development of six *TaNAS* overexpression (*TaNAS*-OE) single locus homozygous lines. (A) Schematic of the pMDC32 construct containing the right border (RB), Ubiquitin promoter 1 (Ubi1) driving either the *TaNAS1-A1*, *TaNAS3-B1*, *TaNAS4-A1*/*A2* chimera, *TaNAS6-D1*, truncated *TaNAS6-D1*, or *TaNAS7-A1* coding sequence, nopaline synthase terminator (nosT), *hygromycin* resistance (HygR), left border (LB), and *kanamycin* resistance (KanR). An alignment of the six *TaNAS* coding sequences with sequence variations (vertical black lines) or deletions (horizontal lines) is indicated. Nucleotide positions are indicated above the alignment. (B) Single locus T_0_ plants were identified by screening T_1_ populations for a transgenic:null ratio of 3:1 (top panel). Homozygous T_1_ plants were identified by screening T_2_ populations for a transgenic proportion of 100% (bottom panel). (C) Both screening methods used a melt curve analysis to rapidly distinguish a null and transgenic samples. A peak at the 84°C (dark blue dashed line) indicates presence of *HygR* (transgenic) and a peak at 87.6°C (light blue dashed line) indicates presence of a bread wheat housekeeping gene (*TaCyclophilin*). (D) Relative gene expression analysis of the *TaNAS* transgene within 18 homozygous transgenic events. The y-axis represents fold-change relative to *TaNAS1-1*. Transgene expression was normalised against the geometric mean of three housekeeping genes (*TaACT*, *TaGAPDH*, and *TaSAR*). The error bars represent the standard error of the mean of three biological replicates (seedlings at the one-leaf stage) (n = 3), where each biological replicate is comprised of three technical replicates. The letters indicate significant differences as determined by a one-way ANOVA (post hoc Tukey’s HSD test). Lines where expression was not detected (n.d.) is indicated. The donor (cv. Fielder) was used as a negative control.

**Figure 5.**
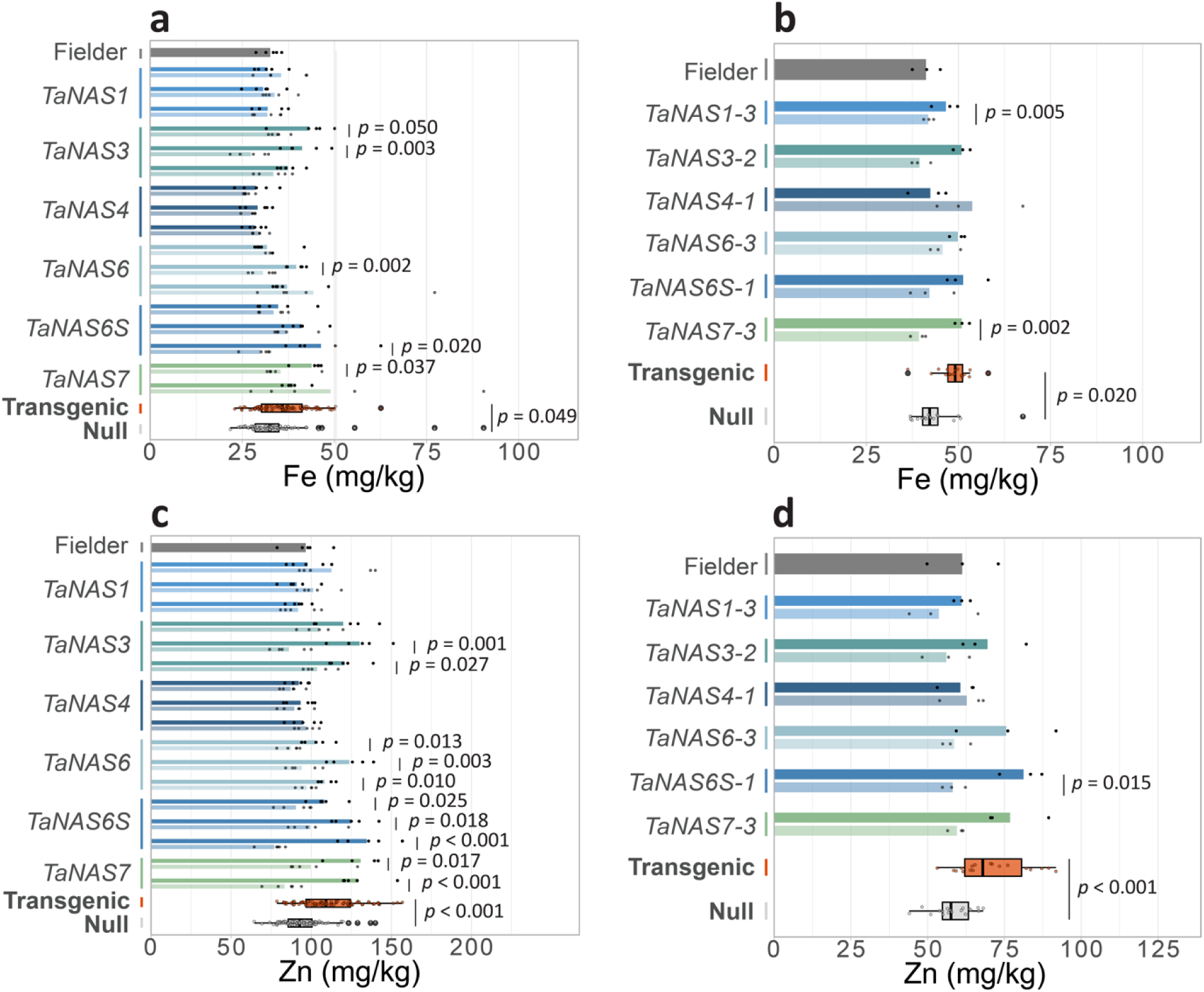
Grain Fe and Zn concentrations of T_3_ *TaNAS*-OE and null plants in glasshouse and field experiments. Grain (A, B) Fe and (C, D) Zn concentrations of *TaNAS1* (blue), *TaNAS3* (teal), *TaNAS4* (navy blue), *TaNAS6* (light blue), *TaNAS6S* (dark blue) and *TaNAS7* (green) lines, respective nulls (faded colour of each), and cv. Fielder (grey) in the glasshouse and field in 2022. For the glasshouse experiment (A, C) events 1, 2 and 3 for each *TaNAS*-OE line are shown at the top, middle and bottom, respectively (n=5). For the field experiment (B, D), events *TaNAS1-3*, *TaNAS3-2*, *TaNAS4-1*, *TaNAS6-3*, *TaNAS6S-1*, and *TaNAS7-3* are shown (n=3). The box plots represent all transgenic (orange) values compared with all null (grey) values. Statistical differences between transgenic events and their respective NS are shown as p values determined by a two-sample Student’s t-test assuming unequal variance with a 95% confidence interval.

### The TaNAS-OE constructs did not impact normal plant growth under glasshouse or field conditions

At the T_1_ generation, no phenotypic differences were detected between *TaNAS*-OE plants and their respective NS (Fig. S3A-F). To expedite grain bulking, the T_2_ generation was speed bred, and the T_3_ *TaNAS*-OE and NS plants were grown under glasshouse and field conditions in 2022. At the T_3_ in the glasshouse, above ground biomass (AGB), tiller number and total grain weight (TGW) were all similar between *TaNAS*-OE plants and their respective NS (Fig. S4). Exceptions include *TaNAS1-2* which had 1.2-fold (*p* = 0.041) higher AGB and 1.3-fold (*p* = 0.011) higher TGW relative to the NS, and *TaNAS3-2* which had 1.2-fold (*p* = 0.028) lower AGB and 1.3-fold (*p* = 0.010) lower TGW relative to the NS (Fig. S4A, C). At the T_3_ in the field, no differences in subsampled grain weight were detected between *TaNAS*-OE plants and their respective NS (Fig. S4C). At the T_4_ in the field, agronomic traits were similar between all *TaNAS*-OE plants and their respective NS (Fig. 6). Exceptions include the *TaNAS1-2* event which had a 1.4-fold higher AGB relative to the NS (*p* = 0.044) and *TaNAS6S-1* event which had 1.4-fold lower tiller number relative to the NS (*p* = 0.039) (Fig. 6A, C).

**Figure 6.**
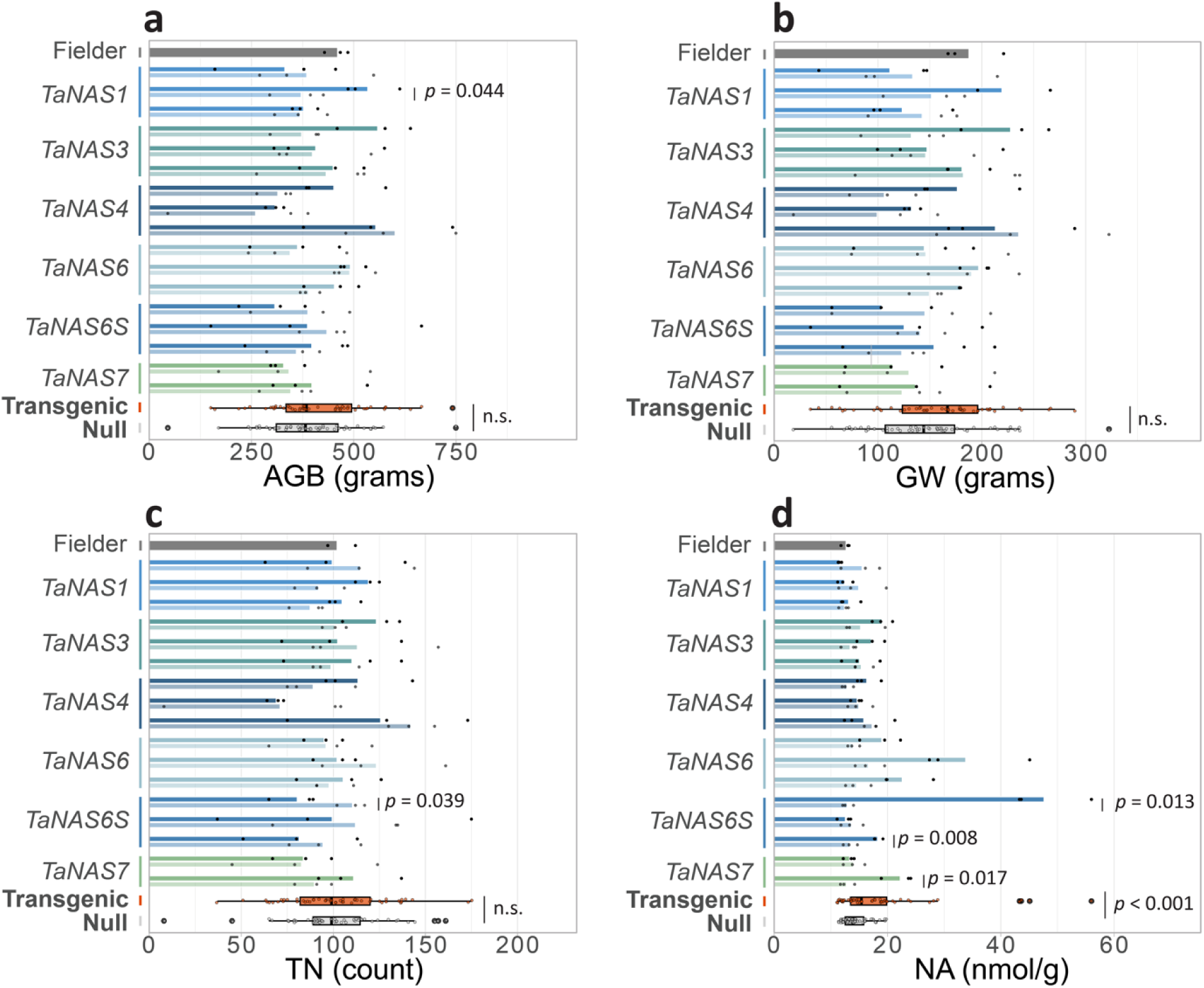
Agronomic traits and grain nicotianamine concentrations of T_4_ *TaNAS*-OE and NS plants in the field. (A) Above ground biomass (AGB, g/m^2^), (B) grain weight (GW, g/m^2^), (C) tiller number (TN/m^2^), and (D) nicotianamine (NA) concentrations (nmol/g) of *TaNAS1* (blue), *TaNAS3* (teal), *TaNAS4* (navy blue), *TaNAS6* (light blue), *TaNAS6S* (dark blue) and *TaNAS7* (green) lines, respective nulls (faded colour of each), and cv. Fielder (grey) in the field in 2023. The donor (cv. Fielder) is at the top (grey bar). Events 1, 2 and 3 for each *TaNAS*-OE line are shown at the top, middle and bottom, respectively (n=3). Statistical differences between transgenic events and their respective NS are shown as p values determined by a two-sample Student’s t-test assuming unequal variance with a 95% confidence interval. The box plots represent all transgenic (orange) values compared with all null (grey) values. Non-significant (n.s.) values between all transgenic and all null segregant values as determined by a two-sample Student’s t-test assuming unequal variance are indicated.

### Overexpressing the TaNAS6-D1 homoeolog led to the highest grain NA in bread wheat

At the T_1_ generation in the glasshouse, grain nicotianamine (NA) concentrations were more than 2-fold higher in *TaNAS3-1* (2.7-fold), *TaNAS3-2* (2.5-fold), *TaNAS6-2* (2.5-fold), *TaNAS6-3* (2.4-fold), *TaNAS6S-1* (13.6-fold), *TaNAS6S-3* (2.0-fold), *TaNAS7-2* (4.2-fold), and *TaNAS7-3* (2.6-fold) grain relative to their respective NS (Fig. S3G-L). Notably, grain 2’-deoxymugineic acid (DMA) concentrations were also 2.6-fold higher in *TaNAS6S-1* relative to the NS (Fig. S3G-H). At the T_4_ generation in the field, grain NA concentrations were 3.7- (*p* = 0.013) and 1.4-fold (*p* = 0.008) higher in *TaNAS6S-1* and *TaNAS6S-3*, respectively relative to the NS (Fig. 6D, Table S4). At the T_3_ in the glasshouse, grain Fe concentrations were 1.2- (*p* = 0.050), 1.5- (*p* = 0.003), 1.3- (*p* = 0.002), 1.6 - (*p* = 0.020), and 1.2-fold (*p* = 0.037) higher in *TaNAS3-1*, *TaNAS3-2*, *TaNAS6-2*, *TaNAS6S-3*, and *TaNAS7-2*, respectively relative to their NS (Fig. 5A, Table S4). Under the same conditions, grain Zn concentrations were between 1.2- to 1.5-fold higher in *TaNAS3,* between 1.1- to 1.3-fold higher in *TaNAS6*, between 1.2- to 1.8-fold higher in *TaNAS6S*, and between 1.3- to 1.6-fold higher in *TaNAS7* plants relative to their respective NS (Fig. 5C, Table S4). At the T_3_ in the field, grain Fe concentrations were 1.3-fold (*p* = 0.005) and 1.3-fold (p = 0.002) higher in *TaNAS3-2* and *TaNAS7-3*, respectively, and grain Zn concentration was 1.40-fold (*p* = 0.015) higher in *TaNAS6S-1*, relative to their respective NS (Fig. 5B, Fig. 5D, Table S4). Grain phosphorus, manganese and copper did not differ significantly between the *TaNAS*-OE lines and their respective NS at the T_3_ generation (Fig. S5, Table S5).

## Discussion

Our analysis of the *TaNAS* gene family in bread wheat included the updated IWGSC reference sequence (Refseq2.1) and outputs of the 10+ Wheat Genome Project (Borrill *et al*., 2016; Appels *et al*., 2018; Ramírez-González *et al*., 2018; Walkowiak *et al*., 2020; Zhu *et al*., 2021). The Apollo platform, which integrates high- and low-confidence gene annotations from IWGSC RefSeq v2.1 and NCBI, alongside RNA-seq and proteome data, provided the resource for manually annotating genes within the bread wheat genome. In Apollo, RNA-seq read alignment validated the expression of several *TaNAS* genes, resolving discrepancies in coding sequence (CDS) annotations between IWGSC RefSeq v2.1 and NCBI for *TaNAS1-B1*, *TaNAS1-D1*, *TaNAS5-B1*, and *TaNAS6-A1* (Table S1). In some cases (for example, the *TaNAS4-D1* and *TaNAS4-D2* genes) the IWGSC RefSeq v2.1 lacked a sufficient annotation, and the RNA-seq data present in Apollo corroborated the annotations from NCBI or IWGSC RefSeq v1.1. The criteria for *TaNAS* gene classification included 100% alignment with the c3fpjA_ template and a transcriptional response to Fe deficiency. Although the YXXΦ and LL motifs are generally considered important for NAS function, the *TaNAS5-B1* and *TaNAS5-B2* genes in cv. Chinese Spring and the *TaNAS5-B2* and *TaNAS5-B3* genes in all other cultivars do not contain these motifs (Fig. 1, Fig. 2, Table S2), as do some eudicot NAS proteins (Nozoye et al., 2014; Seebach et al., 2023). We therefore included these *TaNAS5* genes in our analysis due to their 100% alignment with the c3fpjA_ template, high sequence conservation with other *TaNAS* genes, and transcriptional upregulation under Fe deficiency (Fig. 2B, Fig. 3). However, future *in planta* studies investigating the enzymatic activity of these truncated TaNAS5 proteins that lack these YXXΦ and LL motifs are required to confirm their function.

All TaNAS proteins contained a highly conserved ‘core’ amino acid (aa) sequence (which ranged between 126 to 284 aa in length), flanked by variable N- and C-terminal regions (Fig. 2B). Our *TaNAS* annotations did not identify the N-terminal extensions previously described for *TaNAS5-B1* and *TaNAS6-A1*, due to discrepancies between these original annotations (prior to publication of the first wheat genome reference), Refseq2.1, and NCBI (Bonneau *et al*., 2016). Interestingly, the *TaNAS6-A1* annotation demonstrated inconsistent sequence length in base pairs (bp) between Refseq2.1 (which predicted a truncated protein) and NCBI (which predicted a slightly extended N terminal without transcriptional evidence), so the longest open reading frame (ORF) that contained the YXXΦ and LL motifs, and had RNA-seq evidence of transcription, was annotated. Both Refseq2.1 and NCBI predicted the 162 bp extended N-terminus in *TaNAS6-D1*. However, we were unable to identify any evidence that the extended N-terminus is transcribed or translated in cv. Fielder plants under Fe deficiency or in cv. Chinese Spring developing grain under control conditions (Fig. S2) (Guo *et al*., 2023).

We overexpressed the *TaNAS6-D1* gene with and without this extended N-terminal region and observed that the *TaNAS6S-1* event (without the extended N-terminal region) showed the greatest fold-difference in grain nicotianamine (NA) (3.7-fold), Zn (1.4-fold), and Fe (1.2-fold) concentrations relative to the null segregant (NS) observed under field conditions (Fig. 5B, 5D, 6D, Table S4). Further studies are warranted to determine whether these differences in the *TaNAS6S-1* event translate into changes in grain mineral localisation and/or Fe and Zn bioavailability. Interestingly, overexpressing the *TaNAS6-D1* either with (the *TaNAS6* events) or without (the *TaNAS6S* events) the extended N terminal region were both successful strategies for enhancing grain nutrition without impacting plant growth, indicating that the extended N-terminus does not affect Fe and Zn biofortification of bread wheat (Fig. 5, 6A, S1, S2). While the extended N-termini identified in the original characterisation of TaNAS proteins were predicted to determine subcellular localisation, a separate study in maize concluded that extended N-termini do not influence ZmNAS protein localisation (Emanuelsson *et al*., 2000; Zhou *et al*., 2013; Bonneau *et al*., 2016). Ultimately, additional high quality bread wheat ribosomal sequencing or proteomic datasets across multiple environments and developmental stages will be necessary to confirm whether any of the TaNAS proteins contain a functional extended N-terminus (Guo *et al*., 2023). The TaNAS1 proteins contain truncated C-terminal regions relative to other TaNAS proteins (Fig. 2B), and truncation of the C-terminus in the *Arabidopsis* AtNAS1 protein increases enzymatic activity due to loss of an autoinhibitory motif (TRGCMFMPCNCS) (Seebach *et al*., 2023). Although the same motif is poorly conserved among the TaNAS proteins, other monocot-specific C-terminal autoinhibitory motifs may be present and, given their truncated C-terminal regions, the TaNAS1 proteins may be missing these putative autoinhibitory motifs (Fig. 2B, 4A). Overexpressing the *TaNAS1-A1* gene (the shortest of the *TaNAS1* homoeologs) did not influence grain Fe, Zn, or NA concentrations in all three *TaNAS1* events, suggesting that the TaNAS1-A1 protein may not have increased enzymatic activity due to the absence of a C-terminal autoinhibitory motif (Fig. 5A, 5B). Interestingly, the *TaNAS1-2* event had higher above ground biomass (AGB) and grain weight in the 2022 glasshouse season (2.60-fold, *p* = 0.041 for AGB and 1.33-fold, *p* = 0.011 for grain weight) and 2023 field season (1.43-fold, *p* = 0.044 for AGB and 1.44-fold, *p* = 0.112 for grain weight) relative to the NS (Fig. 6A, S2A, S2C). Whether these changes in *TaNAS1-2* agronomic traits are due to an unexamined effect of *TaNAS1-A1* overexpression on plant growth (e.g., elevated NA concentrations in *TaNAS1-2* vegetative tissues) or the integration of the *TaNAS1* transgene into a genomic location that regulates plant growth is unclear. Further investigation into the determinants of *TaNAS1* agromorphology will occur alongside functional enzymatic assays to determine the presence of autoinhibitory motifs in TaNAS C-termini. The variation in grain Fe, Zn and NA concentrations among independent events of the same construct (e.g. *TaNAS6S-1*-OE, *TaNAS6S-2*-OE, *TaNAS6S-3-*OE) was not explained by transgene expression levels (Fig. 3.4d, 3.5, 3.6d, S3.3g). This finding could be explained by the fact that transgene expression in seedling shoots does not accurately reflect transgene expression at other developmental stages as grain filling. Further characterisation of both transgene expression and site-specific transgene integration may be required to explain the high grain NA concentrations observed in the *TaNAS6S-1*-OE event relative to *TaNAS6S-2*-OE and *TaNAS6S-3*-OE (Fig. 3.6d, S3.3g).

Conventional breeding strategies to increase grain Fe concentrations in wheat continue to be limited by a lack of natural genetic variation in modern breeding populations (Borrill *et al*., 2014; Ludwig and Slamet-Loedin, 2019; Saini *et al*., 2020). Landraces and synthetic varieties display greater variation in grain Fe concentrations but often have lower yield relative to elite cultivars (Heidari *et al*., 2024). Enhancing grain NA concentration *via NAS*-overexpression has long been the most promising single-gene overexpression strategy for Fe and Zn biofortification of rice and wheat (Inoue *et al*., 2003; Wirth *et al*., 2009; Zheng *et al*., 2010; Johnson *et al*., 2011; Lee *et al*., 2012; Eagling *et al*., 2014; Singh *et al*., 2017; Beasley *et al*., 2019*b*,*a*, 2022). While current evidence indicates that *NAS* overexpression in rice does not increase grain cadmium (Cd) concentration under normal and contaminated growth conditions (Trijatmiko *et al*., 2016), the recent observation of elevated seed Cd concentrations in *NAS*-overexpressing *Arabidopsis thaliana* grown on Cd contaminated soil highlights a need for thorough assessment of heavy-metal accumulation in *NAS*-engineered rice and wheat (Hollmann *et al*., 2025). Overexpression of the rice *OsNAS2* gene in bread wheat results in consistently higher NA concentrations and Fe bioavailability in white wheat flour relative to NS controls, with no associated agronomic penalty (Beasley *et al*., 2019*a*, 2022). Here we conducted 3 years of glasshouse and field trials and demonstrated that overexpressing *TaNAS* genes has no negative effects on agronomic performance – a key factor in grower acceptance and adoption (Fig. 5, S1, S2). Furthermore, intragenic strategies (which utilise promoters, genes, terminators and selectable markers that are entirely from sexually compatible species) are expected to face fewer regulatory challenges than conventional transgenic methods (Espinoza *et al*., 2013; Holme *et al*., 2013). The results described in this study, along with the updated repository of *TaNAS* genes present in multiple wheat genomes, provide breeders with a wealth of conventional, precision and intragenic resources to enhance bread wheat nutrition.

## Supplementary Data

The following supplementary data are available online.

Fig. S1. Alignment of 425 *TaNAS* coding sequences from cvs. Chinese Spring, ArinaLrFor, Julius, Lancer, Norin61, SY Mattis, Mace, Jagger, Landmark, Stanley, Fielder, and *Triticum spelta* L.

Fig. S2. Ribosomal profile and RNA-seq coverage of *TaNAS6-D1*.

Fig. S3. Phenotypic and nutritional analysis of T_1_ *TaNAS*-OE and NS plants in the glasshouse.

Fig. S4. Agronomic traits of T_3_ *TaNAS*-OE and NS plants in the glasshouse and field.

Fig. S5. Grain Mn, Cu, and P concentrations of T_3_ *TaNAS*-OE and null plants in glasshouse and field experiments.

Table S1. Gene models and chromosomal location of 34 *TaNAS* genes in bread wheat cv. Chinese Spring and the additional 9 potential *TaNAS* genes that were excluded from this study.

Table S2: Expression patterns and protein features of 34 *TaNAS* genes in bread wheat cv. Chinese Spring and the additional 9 potential *TaNAS* genes that were excluded from this study.

Table S3. Haplotypes of *TaNAS* genes in 11 different bread wheat cultivars and *Triticum spelta* 1. L.

Table S4. Grain Fe, Zn, and NA fold-differences between *TaNAS*-OE and null plants at the T_3_ and T_4_ generations.

Table S5. Grain P, Mn, and Cu fold-differences between *TaNAS*-OE and null plants at the T_3_ generation.

Table S6. Gene IDs and coding sequences used as queries to blast for novel bread wheat

*NAS* genes.

## Acknowledgements

This research was supported by a grant from the Australian Research Council (LP190100631). We acknowledge The University of Adelaide for bread wheat transformation services, The University of Melbourne TrACEES platform for elemental analysis, and the Australian Apollo Service for genome assembly and curation software. We are grateful to Nathan Anderson, Tim Reeks, and numerous volunteers – Rucha Patil, Amy Liu, Chris Beasley, Stuart Roy, Abdeljalil El Habti, Hannah Ebert, Bhawana Bhatta, Troy Bentley, Bill Robinson, Rajiv Thapa, and Shubham Vishwakarma – for assistance in setting up and maintaining wheat field trials. We also thank Julien Bonneau and Martin O’Brien for construct assembly.

## Author contributions

AATJ, JTB, OCF, EJF, and RA conceptualised the project. AATJ acquired the project funding and provided resources. AATJ, JTB, and RA provided supervision. OCF, JTB, EJF, LTS, DLC, and NL performed the experiments. OCF and EJF curated the data and performed the analyses. OCF and JTB prepared the original draft. All authors edited and reviewed the final draft of the manuscript.

## Conflict of interest

The researchers declare no conflict of interest.

## Funding

This work was supported by the Australian Research Council Linkage Project (LP190100631).

## Data availability

Raw data is included in the supplementary information within this manuscript and RNA-seq data is available from NCBI-GEO (GEO accession no. GSE283676).

## Notes

### Competing Interest Statement

The authors have declared no competing interest.

