## Supplementary tables and figures for "Expanding the *TaNAS* gene family in bread wheat and exploring potential for intragenic biofortification"

**Table S1. Gene models and chromosomal location of 34 *TaNAS* genes in bread wheat cv. Chinese Spring and the additional 9 potential *TaNAS* genes that were excluded from this study.** Gene models (IWGSC RefSeq1.1, 2.1, and NCBI) of the included *TaNAS* genes (rows 1-34) and excluded *TaNAS* genes (rows 35-43), and their chromosomal location (IWGSC RefSeq2.1) are described.

| No | New annotation | IWGSC RefSeqv1.1 | IWGSC RefSeqv2.1 | NCBI annotation | Chromosomal Location | CDS annotation used |
|---|---|---|---|---|---|---|
| 1 | ***TaNAS1-A1*** | TraesCS2A02G033500 | TraesCS2A03G066300 | LOC123187039 | Chr2A:18046576-18048604 - | Refseq2.1 |
| 2 | ***TaNAS1-B1*** | TraesCS2B02G047100 | TraesCS2B03G098000 | LOC123043227 | Chr2B:26588743-26592485 - | Refseq2.1 |
| 3 | ***TaNAS1-D1*** | TraesCS2D02G033000 | TraesCS2D03G061000 | LOC123051093 | Chr2D:13092629-13096137 - | Refseq2.1 |
| 4 | ***TaNAS2-A1*** | TraesCS6A02G163100 | TraesCS6A03G395600 | LOC123130501 | Chr6A:160986866-160988156 - | Refseq2.1 |
| 5 | ***TaNAS2-A2*** | TraesCS6A02G163200 | TraesCS6A03G396300 | LOC123130503 | Chr6A:161119099-161125376 - | Refseq2.1 |
| 6 | ***TaNAS2-A3*** | TraesCS6A02G165100 | TraesCS6A03G404500 | LOC123130517 | Chr6A:165538805-165539794 - | Refseq2.1 |
| 7 | ***TaNAS2-B1*** | TraesCS6B02G187400 | TraesCS6B03G477500 | LOC123133995 | Chr6B:219612150-219613139 - | Refseq2.1 |
| 8 | ***TaNAS2-B2*** | TraesCS6B02G186100 | TraesCS6B03G474000 | LOC123133989 | Chr6B:218289819-218290808 - | Refseq2.1 |
| 9 | ***TaNAS2-D1*** | TraesCS6D02G148200 | TraesCS6D03G330700 | LOC123141358 | Chr6D:142768539-142771342 + | Refseq2.1 |
| 10 | ***TaNAS2-D2*** | TraesCS6D02G148600 | TraesCS6D03G331700 | LOC123144159 | Chr6D:143120609-143121939 - | Refseq2.1 |
| 11 | ***TaNAS2-D3*** | TraesCS6D02G148000 | TraesCS6D03G330000 | LOC123141357 | Chr6D:142582221-142583640 + | Refseq2.1 |
| 12 | ***TaNAS3-A1*** | TraesCS2A02G049900 | TraesCS2A03G093200 | LOC123184397 | Chr2A:22269439-22270719 - | Refseq2.1 |
| 13 | ***TaNAS3-B1*** | TraesCS2B02G060800 | TraesCS2B03G130700 | LOC123040191 | Chr2B:32997616-32998896 - | Refseq2.1 |
| 14 | ***TaNAS3-D1*** | TraesCS2D02G049200 | TraesCS2D03G092200 | LOC123048357 | Chr2D:18625470-18626900 - | Refseq2.1 |
| 15 | ***TaNAS4-A1*** | TraesCS5A02G552400 | TraesCS5A03G1289600 | LOC123106177 | Chr5A:708105606-708106934 - | Refseq2.1 |
| 16 | ***TaNAS4-A2*** | TraesCS5A02G552000 | TraesCS5A03G1289100 | LOC123106176 | Chr5A:708052165-708053166 - | Refseq2.1 |

| | | | | | | |
|---|---|---|---|---|---|---|
| 17 | ***TaNAS4-D1*** | TraesCSU02G125200 | n/a | LOC12309921<br>6 | Chr4D:513233447-<br>513234914 - | Refseq1.1 |
| 18 | ***TaNAS4-D2*** | TraesCSU02G125500 | n/a | LOC12309921<br>4 | Chr4D:513215952-<br>513217368 + | Refseq1.1 |
| 19 | ***TaNAS5-B1*** | TraesCS3B02G068500 | TraesCS3B03G154800 | LOC12306466<br>1 | Chr3B:50315898-<br>50316948 + | Refseq2.1 |
| 20 | ***TaNAS5-B2*** | TraesCS3B02G068400 | TraesCS3B03G154700 | n/a | Chr3B:50077736-<br>50078278 + | Refseq2.1 |
| 21 | ***TaNAS6-A1*** | TraesCS4A02G127900LC | TraesCS4A03G249100LC | LOC12308376<br>6 | Chr4A:148780629-<br>148781781 + | Manual<br>annotation |
| 22 | ***TaNAS6-B1*** | TraesCS4B02G183900 | TraesCS4B03G520200 | LOC12309488<br>5 | Chr4B:402432887-<br>402433879 - | Refseq2.1 |
| 23 | ***TaNAS6-D1*** | TraesCS4D02G184900 | TraesCS4D03G458900 | LOC12309767<br>8 | Chr4D:323251547-<br>323253910 - | Refseq2.1 |
| 24 | ***TaNAS7-A1*** | TraesCS6A02G093000 | TraesCS6A03G214000 | LOC12313029<br>2 | Chr6A:63817836-<br>63819203 - | Refseq2.1 |
| 25 | ***TaNAS7-A2*** | TraesCS6A02G386200 | TraesCS6A03G973000 | LOC12313132<br>5 | Chr6A:606021899-<br>606022891 + | Refseq2.1 |
| 26 | ***TaNAS7-B1*** | TraesCS6B02G425200 | TraesCS6B03G1190400 | LOC12313501<br>2 | Chr6B:703619814-<br>703620806 + | Refseq2.1 |
| 27 | ***TaNAS7-D1*** | TraesCS6D02G370800 | TraesCS6D03G854100 | LOC12314611<br>7 | Chr6D:477989584-<br>477990867 + | Refseq2.1 |
| 28 | ***TaNAS8-A1*** | TraesCS6A02G398700 | TraesCS6A03G1000600 | LOC12313137<br>1 | Chr6A:612027263-<br>612028597 - | Refseq2.1 |
| 29 | ***TaNAS8-B1*** | TraesCS6B02G438900 | TraesCS6B03G1227100 | LOC12313509<br>9 | Chr6B:714051809-<br>714053050 - | Refseq2.1 |
| 30 | ***TaNAS8-D1*** | TraesCS6D02G382900 | TraesCS6D03G881800 | LOC12314218<br>2 | Chr6D:483833945-<br>483835250 + | Refseq2.1 |
| 31 | ***TaNAS9-A1*** | TraesCS2A02G095700 | TraesCS2A03G196300 | LOC12318740<br>3 | Chr2A:53939313-<br>53940335 + | Refseq2.1 |
| 32 | ***TaNAS9-B1*** | TraesCS2B02G111100 | TraesCS2B03G264600 | LOC12304359<br>8 | Chr2B:80456493-<br>80458103 + | Refseq2.1 |
| 33 | ***TaNAS9-D1*** | TraesCS2D02G094200 | TraesCS2D03G196500 | LOC12305146<br>5 | Chr2D:48272558-<br>48273580 + | Refseq2.1 |
| 34 | ***TaNAS10-A1*** | TraesCS3A02G468300 | TraesCS3A03G1088500 | LOC12305924<br>4 | Chr3A:700464391-<br>700465811 + | Refseq2.1 |
| 35 | | TraesCS2A02G031500LC | TraesCS2A03G065900LC | | Chr2A:18030402-<br>18030782 - | Refseq2.1 |

| | | | | | |
|---|---|---|---|---|---|
| 36 | n/a | n/a | n/a | Chr3B:50111254-50113113 + | n/a |
| 37 | TraesCS3D02G055800 | TraesCS3D03G103600 | n/a | Chr3D:23087305-23088112 + | Refseq2.1 |
| 38 | TraesCS4A02G128000LC | TraesCS4A03G249200LC | n/a | Chr4A:148782303-148782891 + | Refseq2.1 |
| 39 | TraesCS4B02G599900LC | TraesCS4B03G989100LC | n/a | Chr4B:665502039-665503016 + | Nontranslating CDS |
| 40 | TraesCS4B02G600600LC | TraesCS4B03G989900LC | n/a | Chr4B:665533725-665543791 - | Nontranslating CDS |
| 41 | TraesCSU02G125100 | n/a | n/a | Chr4D:513253441-513254120 - | Refseq1.1 |
| 42 | TraesCS5A02G733600LC | TraesCS5A03G1288900LC | n/a | Chr5A:708031745-708032418 + | Refseq2.1 |
| 43 | TraesCS7B02G813200LC | TraesCS7B03G1308900LC | n/a | Chr7B:755163999-755164979 + | n/a |

**Table S2. Expression patterns and protein features of 34 *TaNAS* genes in bread wheat cv. Chinese Spring and the additional 9 potential *TaNAS* genes that were excluded from this study.** Gene expression of root, leaf, stem, spike or grains at all time points (INRA RNA-seq viewed in Apollo) are categorised with 0-5 (n/a), 5-100 (Low), 100-500 (Med), and greater than 500 (High) maximum RNA-seq reads. Upregulation or down regulation of a *TaNAS* gene in response to Fe deficiency was indicated with a 'Yes', and no change in expression was indicated with a 'No', as determined by RNA-seq data generated in this study. Protein features including the alignment of a TaNAS protein to the c3fpjA_ template with 100% confidence is indicated with a 'Yes', and lower confidence or no alignment to the c3fpjA_ template is indicated. Protein features where genes could not be annotated with confidence are indicated with 'n/a'.

| No | New annotation | Root expression (Apollo) | Leaf expression (Apollo) | Stem expression (Apollo) | Spike expression (Apollo) | Grain expression (Apollo) | Response to Fe deficiency | c3fpjA_ protein | YXXΦ | LL | Length |
|---|---|---|---|---|---|---|---|---|---|---|---|
| 1 | ***TaNAS1-A1*** | High | Low | Med | Med | Low | Yes | Yes | Yes | Yes | 843 |
| 2 | ***TaNAS1-B1*** | High | Low | Low | Low | Low | Yes | Yes | Yes | Yes | 933 |
| 3 | ***TaNAS1-D1*** | High | Med | Med | Med | Med | Yes | Yes | Yes | Yes | 894 |
| 4 | ***TaNAS2-A1*** | High | n/a | n/a | Low | Low | Yes | Yes | Yes | Yes | 981 |
| 5 | ***TaNAS2-A2*** | High | n/a | n/a | n/a | n/a | Yes | Yes | Yes | Yes | 951 |
| 6 | ***TaNAS2-A3*** | High | n/a | n/a | n/a | n/a | Yes | Yes | Yes | Yes | 990 |
| 7 | ***TaNAS2-B1*** | High | n/a | n/a | n/a | n/a | Yes | Yes | Yes | Yes | 990 |
| 8 | ***TaNAS2-B2*** | High | n/a | n/a | low | n/a | Yes | Yes | Yes | Yes | 990 |
| 9 | ***TaNAS2-D1*** | High | n/a | n/a | n/a | n/a | Yes | Yes | Yes | Yes | 990 |
| 10 | ***TaNAS2-D2*** | High | n/a | n/a | n/a | n/a | Yes | Yes | Yes | Yes | 990 |
| 11 | ***TaNAS2-D3*** | High | n/a | n/a | n/a | n/a | Yes | Yes | Yes | Yes | 990 |
| 12 | ***TaNAS3-A1*** | High | n/a | n/a | n/a | n/a | Yes | Yes | Yes | Yes | 993 |
| 13 | ***TaNAS3-B1*** | High | n/a | n/a | Low | n/a | Yes | Yes | Yes | Yes | 993 |
| 14 | ***TaNAS3-D1*** | High | n/a | n/a | Low | n/a | Yes | Yes | Yes | Yes | 993 |
| 15 | ***TaNAS4-A1*** | High | n/a | n/a | Low | n/a | Yes | Yes | Yes | Yes | 1002 |
| 16 | ***TaNAS4-A2*** | High | n/a | n/a | n/a | n/a | Yes | Yes | Yes | Yes | 1002 |
| 17 | ***TaNAS4-D1*** | High | Low | Low | Low | n/a | Yes | Yes | Yes | Yes | 1002 |

| | | | | | | | | | | | |
|---|---|---|---|---|---|---|---|---|---|---|---|
| 18 | ***TaNAS4-D2*** | High | Low | Low | Low | n/a | Yes | Yes | Yes | Yes | 1002 |
| 19 | ***TaNAS5-B1*** | High | n/a | Low | Low | n/a | Yes | Yes | No | No | 543 |
| 20 | ***TaNAS5-B2*** | High | n/a | n/a | n/a | n/a | Yes | Yes | No | No | 543 |
| 21 | ***TaNAS6-A1*** | High | n/a | n/a | n/a | Low | Yes | Yes | Yes | Yes | 711 |
| 22 | ***TaNAS6-B1*** | High | n/a | n/a | n/a | Med | Yes | Yes | Yes | Yes | 993 |
| 23 | ***TaNAS6-D1*** | High | Low | Low | Low | Med | Yes | Yes | Yes | Yes | 1155 |
| 24 | ***TaNAS7-A1*** | High | n/a | n/a | n/a | n/a | Yes | Yes | Yes | Yes | 993 |
| 25 | ***TaNAS7-A2*** | Low | n/a | n/a | Low | n/a | Yes | Yes | Yes | Yes | 993 |
| 26 | ***TaNAS7-B1*** | High | n/a | n/a | n/a | n/a | Yes | Yes | Yes | Yes | 993 |
| 27 | ***TaNAS7-D1*** | High | n/a | n/a | n/a | n/a | Yes | Yes | Yes | Yes | 993 |
| 28 | ***TaNAS8-A1*** | High | n/a | n/a | n/a | n/a | Yes | Yes | Yes | Yes | 987 |
| 29 | ***TaNAS8-B1*** | High | n/a | n/a | n/a | n/a | Yes | Yes | Yes | Yes | 987 |
| 30 | ***TaNAS8-D1*** | High | n/a | n/a | n/a | n/a | Yes | Yes | Yes | Yes | 987 |
| 31 | ***TaNAS9-A1*** | Med | High | High | Low | Med | Yes | Yes | Yes | Yes | 1023 |
| 32 | ***TaNAS9-B1*** | Med | High | High | Med | Med | Yes | Yes | Yes | Yes | 1023 |
| 33 | ***TaNAS9-D1*** | Low | High | High | Med | Med | Yes | Yes | Yes | Yes | 1023 |
| 34 | ***TaNAS10-A1*** | Med | n/a | n/a | n/a | n/a | Yes | Yes | Yes | Yes | 1023 |
| 35 | | n/a | n/a | n/a | n/a | n/a | No | No | No | No | 381 |
| 36 | | n/a | n/a | n/a | n/a | n/a | No | n/a | n/a | n/a | n/a |
| 37 | | Low | n/a | n/a | n/a | n/a | Low | 99.9 | No | No | 648 |
| 38 | | Low | n/a | n/a | n/a | n/a | Yes | No | No | No | 177 |
| 39 | | n/a | n/a | n/a | n/a | n/a | No | n/a | n/a | n/a | n/a |
| 40 | | n/a | n/a | n/a | n/a | n/a | No | No | n/a | n/a | n/a |
| 41 | | Med | n/a | n/a | n/a | n/a | Yes | No | No | No | 315 |
| 42 | | n/a | n/a | n/a | n/a | n/a | No | 99.9 | Yes | Yes | 498 |
| 43 | | n/a | n/a | n/a | n/a | n/a | No | n/a | n/a | n/a | n/a |

**Table S3. Haplotypes of *TaNAS* genes in 11 different bread wheat cultivars and *Triticum spelta* L.** *TaNAS* genes present in cvs. Chinese Spring, ArinaLrFor, Julius, Lance, Norin61, SY Mattis, Mace, *Triticum spelta* L., Jagger, Landmark, and Stanley are in first section of the table (rows 1-137). *TaNAS* genes not present in cv. Chinese Spring, but present in other cultivars are in the second section of the table (rows 138-154). Gaps in cells indicate that the *TaNAS* gene was not identified. Chromosomal locations were supplied where gene models were not available. The gene models for cv. Chinese Spring used IWGSC RefSeq2.1. Gene and protein variations between the 11 different bread wheat cultivars and *Triticum spelta* L. with Chinese Spring are indicated except for *TaNAS3-B1* and *TaNAS6-A1* where the cv. Chinese Spring *TaNAS* gene variation is unique relative to all other cultivars. The *TaNAS2-B3*, *TaNAS4-A3*, *TaNAS5-B3*, and *TaNAS8-B2* genes are not present in cv. Chinese Spring and variations are compared to the most common haplotype (H). Nucleotide (nt), Single Nucleotide Polymorphisms (SNPs), conserved (cv) and non-conserved (ncs) amino acid (aa) substitutions (sub), replacements (rep), extensions (ext), truncations (trun), deletions (del), insertions (ins) at the N-terminus (N-ter), and C-terminus (C-ter) are indicated.

| Gene | Chinese Spring RefSeq2.1 | ArinaLrFor | Julius | Lancer | Norin61 | SY Mattis | Mace | Triticum spelta | Jagger | Landmark | Stanley | Fielder |
|---|---|---|---|---|---|---|---|---|---|---|---|---|
| **TaNAS1-A1** | **TraesCS2A03G066300** | **TraesARI2A03G00595190** | **TraesJUL2A03G00591680** | **TraesLAC2A03G00592550** | **TraesNOR2A03G00596190** | **TraesSYM2A03G00595200** | **TraesMAC2A03G00587670** | **TraesTSP2A01G037400** | **TraesJAG2A03G00587010** | **TraesLDM2A03G00590820** | **TraesSTA2A03G00587780** | **2A:14977723-14976881 Reverse** |
| Gene | - | 100% identical | 50→83 nt rep (3') | 100% identical | 100% identical | 57 SNPs | 57 SNPs | 46 SNPs | 57 SNPs | 100% identical | 57 SNPs | 100% identical |
| Protein | - | - | 16→27 aa rep (C-ter) | - | - | 9 cs & 6 ncs aa subs, 6 aa ext (C-ter) | 9 cs & 6 ncs aa subs, 6 aa ext (C-ter) | 4 cs & 8 ncs aa subs | 9 cs & 6 ncs aa subs, 6 aa ext (C-ter) | - | 9 cs & 6 ncs aa subs, 6 aa ext (C-ter) | - |
| Haplotype | H1 | H1 | H4 | H1 | H1 | H2 | H2 | H3 | H2 | H1 | H2 | H1 |
| **TaNAS1-B1** | **TraesCS2B03G098000** | **TraesARI2B03G00842250** | **TraesJUL2B03G00837550** | **TraesLAC2B03G00829000** | **TraesNOR2B03G00843360** | **TraesSYM2B03G00842670** | **TraesMAC2B03G00830370** | **TraesTSP2B01G050200** | **TraesJAG2B03G00832430** | **TraesLDM2B03G00834640** | **TraesSTA2B03G00833000** | **2B:23550688-23549857 Reverse** |
| Gene | - | 100% identical | 100% identical | 100% identical | 100% identical | 4 SNPs | 100% identical | 3 nt del | 100% identical | 100% identical | 100% identical | 100% identical |
| Protein | - | - | - | - | - | 1 cs aa sub | - | 1 aa del | - | - | - | - |
| Haplotype | H1 | H1 | H1 | H1 | H1 | H2 | H1 | H3 | H1 | H1 | H1 | H1 |
| **TaNAS1-D1** | **TraesCS2D03G061000** | **TraesARI2D03G01105860** | **TraesJUL2D03G01095670** | **TraesLAC2D03G01042340** | **TraesNOR2D03G01105730** | **TraesSYM2D03G01104740** | **TraesMAC2D03G01089010** | **TraesTSP2D01G038800** | **TraesJAG2D03G01093810** | **TraesLDM2D03G01091450** | **TraesSTA2D03G01078740** | **2D:12872919-12872085 Reverse** |

| | | | | | | | | | | | | |
|---|---|---|---|---|---|---|---|---|---|---|---|---|
| Gene | - | 59→26 nt rep (3') | 23 SNPs | 59→26 nt rep (3') | 59→26 nt rep (3') | 59→26 nt rep (3') | 59→26 nt rep (3') | 100% identical | 59→26 nt rep (3') | 59→26 nt rep (3') | 59→26 nt rep (3') | 59→26 nt rep (3') |
| Protein | - | 19→8 aa rep (C-ter) | 2 cs & 4 ncs aa subs | 19→8 aa rep (C-ter) | 19→8 aa rep (C-ter) | 19→8 aa rep (C-ter) | 19→8 aa rep (C-ter) | - | 19→8 aa rep (C-ter) | 19→8 aa rep (C-ter) | 19→8 aa rep (C-ter) | 19→8 aa rep (C-ter) |
| Haplotype | H2 | H1 | H3 | H1 | H1 | H1 | H1 | H2 | H1 | H1 | H1 | H1 |
| **TaNAS2-A1** | **TraesCS6A03G395600** | **TraesARI6A03G03251690** | **TraesJUL6A03G03322480** | **TraesLAC6A03G03251150** | **TraesNOR6A03G03327840** | **TraesSYM6A03G03237010** | **TraesMAC6A03G03295720** | **TraesTSP6A01G136800** | **TraesJAG6A03G03290620** | **TraesLDM6A03G03299590** | **TraesSTA6A03G03287000** | **6A:158317844-158316864 Reverse** |
| Gene | - | 100% identical | 100% identical | 100% identical | 100% identical | 100% identical | 100% identical | 100% identical | 100% identical | 100% identical | 100% identical | 100% identical |
| Protein | - | - | - | - | - | - | - | - | - | - | - | - |
| Haplotype | H1 | H1 | H1 | H1 | H1 | H1 | H1 | H1 | H1 | H1 | H1 | H1 |
| **TaNAS2-A2** | **TraesCS6A03G396300** | **TraesARI6A03G03251820** | **TraesJUL6A03G03322590** | **TraesLAC6A03G03251260** | **TraesNOR6A03G03327970** | **TraesSYM6A03G03237130** | **TraesMAC6A03G03295840** | **TraesTSP6A01G137000** | **TraesJAG6A03G03290740** | **TraesLDM6A03G03299720** | **TraesSTA6A03G03287100** | **6A:158454932-158448655 Reverse** |
| Gene | - | 30 nt ins, 2 SNPs | 33 nt ins, 1 SNP | 30 nt ins, 1 SNP | 33 nt ins, 1 SNP | 30 nt ins, 1 SNP | 30 nt ins, 2 SNPs | 1 SNP | 33 nt ins, 1 SNP | 2 SNPs | 2 SNPs | 1 SNP |
| Protein | - | 10 aa insert, 1 ncs aa sub | 11 aa ins, 1 ncs aa sub | 10 aa ins, 1 ncs aa sub | 11 aa ins, 1 ncs aa sub | 10 aa ins, 1 ncs aa sub | 10 aa ins, 1 ncs aa sub | 1 ncs aa sub | 10 aa ins, 1 ncs aa sub | 1 ncs sub | 1 ncs sub | 1 ncs aa sub |
| Haplotype | H7 | H4 | H6 | H3 | H5 | H3 | H4 | H2 | H5 | H1 | H1 | H2 |
| **TaNAS2-A3** | **TraesCS6A03G404500** | **TraesARI6A03G03253210** | **TraesJUL6A03G03323910** | **TraesLAC6A03G03252550** | **TraesNOR6A03G03329190** | **TraesSYM6A03G03238460** | **TraesMAC6A03G03297120** | **TraesTSP6A01G139100** | **TraesJAG6A03G03291990** | **TraesLDM6A03G03301050** | **TraesSTA6A03G03288370** | **6A:162891463-162890474 Reverse** |
| Gene | - | 100% identical | 100% identical | 100% identical | 100% identical | 100% identical | 100% identical | 100% identical | 100% identical | 100% identical | 100% identical | 100% identical |
| Protein | - | - | - | - | - | - | - | - | - | - | - | - |
| Haplotype | H1 | H1 | H1 | H1 | H1 | H1 | H1 | H1 | H1 | H1 | H1 | H1 |
| **TaNAS2-B1** | **TraesCS6B03G477500** | **TraesARI6B03G03457020** | **TraesJUL6B03G03520670** | **TraesLAC6B03G03452760** | **TraesNOR6B03G03532210** | **TraesSYM6B03G03436680** | **TraesMAC6B03G03496990** | **TraesTSP6B01G203600** | **TraesJAG6B03G03488970** | **TraesLDM6B03G03502950** | **TraesSTA6B03G03489330** | **6B:213488328-213487339 Reverse** |

| | | | | | | | | | | | | |
|---|---|---|---|---|---|---|---|---|---|---|---|---|
| Gene | - | 100% identical | 100% identical | 1 SNP | 100% identical | 100% identical | 1 SNP | 100% identical | 23 SNPs | 100% identical | 23 SNPs | 100% identical |
| Protein | - | - | - | Identical | - | - | Identical | - | 3 cs & 3 ncs subs | - | 3 cs & 3 ncs subs | - |
| Haplotype | H1 | H1 | H1 | H2 | H1 | H1 | H2 | H1 | H3 | H1 | H3 | H1 |
| **TaNAS2-B2** | **TraesCS6B03G474000** | **TraesARI6B03G03456420** | **TraesJUL6B03G03521300** | **TraesLAC6B03G03452140** | **TraesNOR6B03G03531560** | **TraesSYM6B03G03437300** | **TraesMAC6B03G03496390** | **TraesTSP6B01G202200** | **TraesJAG6B03G03488340** | **TraesLDM6B03G03502410** | **TraesSTA6B03G03488680** | **6B:212159685-212158696 Reverse** |
| Gene | - | 100% identical | 6 SNPs | 100% identical | 100% identical | 6 SNPs | 100% identical | 100% identical | 12 SNPs | 100% identical | 12 SNPs | 2 SNPs |
| Protein | - | - | 1 cs & 1 ncs aa sub | - | - | 1 cs & 1 ncs aa sub | - | - | 2 cs & 3 ncs aa subs | - | 2 cs & 3 ncs aa subs | 1 ncs aa sub |
| Haplotype | H1 | H1 | H3 | H1 | H1 | H3 | H1 | H1 | H4 | H1 | H4 | H2 |
| **TaNAS2-D1** | **TraesCS6D03G330700** | **TraesARI6D03G03658000** | **TraesJUL6D03G03726950** | **TraesLAC6D03G03644720** | **TraesNOR6D03G03734690** | **TraesSYM6D03G03641330** | **TraesMAC6D03G03692730** | **TraesTSP6D01G156200** | **TraesJAG6D03G03677370** | **TraesLDM6D03G03697920** | **TraesSTA6D03G03688240** | **6D:121225558-121226547 Forward** |
| Gene | - | 100% identical | 100% identical | 100% identical | 100% identical | 100% identical | 100% identical | 100% identical | 100% identical | 100% identical | 100% identical | 100% identical |
| Protein | - | - | - | - | - | - | - | - | - | - | - | - |
| Haplotype | H1 | H1 | H1 | H1 | H1 | H1 | H1 | H1 | H1 | H1 | H1 | H1 |
| **TaNAS2-D2** | **TraesCS6D03G331700** | **TraesARI6D03G03658210** | **TraesJUL6D03G03727150** | **TraesLAC6D03G03644920** | **TraesNOR6D03G03734910** | **TraesSYM6D03G03641550** | **TraesMAC6D03G03692940** | **TraesTSP6D01G156600** | **TraesJAG6D03G03677580** | **TraesLDM6D03G03698140** | **TraesSTA6D03G03688470** | **6D:121580460-121579471 Reverse** |
| Gene | - | 100% identical | 100% identical | 100% identical | 100% identical | 100% identical | 100% identical | 100% identical | 100% identical | 100% identical | 100% identical | 100% identical |
| Protein | - | - | - | - | - | - | - | - | - | - | - | - |
| Haplotype | H1 | H1 | H1 | H1 | H1 | H1 | H1 | H1 | H1 | H1 | H1 | H1 |
| **TaNAS2-D3** | **TraesCS6D03G330000** | **TraesARI6D03G03657880** | **TraesJUL6D03G03726840** | **TraesLAC6D03G03644600** | **TraesNOR6D03G03734570** | **TraesSYM6D03G03641210** | **TraesMAC6D03G03692600** | **TraesTSP6D01G156000** | **TraesJAG6D03G03677250** | **TraesLDM6D03G03697800** | **TraesSTA6D03G03688120** | **6D:121064488-121065477 Forward** |

| | | | | | | | | | | | | |
|---|---|---|---|---|---|---|---|---|---|---|---|---|
| Gene | - | 100% identical | 100% identical | 100% identical | 100% identical | 100% identical | 100% identical | 100% identical | 100% identical | 100% identical | 100% identical | 100% identical |
| Protein | - | - | - | - | - | - | - | - | - | - | - | - |
| Haplotype | H1 | H1 | H1 | H1 | H1 | H1 | H1 | H1 | H1 | H1 | H1 | H1 |
| **TaNAS3-A1** | **TraesCS2A03G093200** | **TraesARI2A03G00598990** | **TraesJUL2A03G00595510** | **TraesLAC2A03G00596440** | **TraesNOR2A03G00600070** | **TraesSYM2A03G00599320** | **TraesMAC2A03G00591870** | **TraesTSP2A01G052900** | **TraesJAG2A03G00591190** | **TraesLDM2A03G00594610** | **TraesSTA2A03G00591890** | **2A:19164152-19163160 Reverse** |
| Gene | - | 100% identical | 100% identical | 100% identical | 100% identical | 42 SNPs | 42 SNPs | 22 SNPs | 42 SNPs | 100% identical | 42 SNPs | 100% identical |
| Protein | - | - | - | - | - | 7 cs & 7 ncs aa subs | 7 cs & 7 ncs aa subs | 1 cs & 2 ncs aa subs | 7 cs & 7 ncs aa subs | - | 7 cs & 7 ncs aa subs | - |
| Haplotype | H1 | H1 | H1 | H1 | H1 | H2 | H2 | H3 | H2 | H1 | H2 | H1 |
| **TaNAS3-B1** | **TraesCS2B03G130700** | **TraesARI2B03G00849120** | **TraesJUL2B03G00841820** | **TraesLAC2B03G00833920** | **TraesNOR2B03G00848160** | **TraesSYM2B03G00847200** | **TraesMAC2B03G00835170** | **TraesTSP2B01G068000** | **TraesJAG2B03G00837080** | **TraesLDM2B03G00839040** | **TraesSTA2B03G00837720** | **2B:29119026-29120018 Forward** |
| Gene | 2 SNPs | 100% identical | 100% identical | 100% identical | 100% identical | 100% identical | 100% identical | 100% identical | 100% identical | 100% identical | 100% identical | 100% identical |
| Protein | Identical | - | - | - | - | - | - | - | - | - | - | - |
| Haplotype | H2 | H1 | H1 | H1 | H1 | H1 | H1 | H1 | H1 | H1 | H1 | H1 |
| **TaNAS3-D1** | **TraesCS2D03G092200** | **TraesARI2D03G01109930** | **TraesJUL2D03G01099970** | **TraesLAC2D03G01045510** | **TraesNOR2D03G01109800** | **TraesSYM2D03G01108480** | **TraesMAC2D03G01092650** | **TraesTSP2D01G057400** | **TraesJAG2D03G01097390** | **TraesLDM2D03G01095590** | **TraesSTA2D03G01082660** | **2D:18169945-18168953 Reverse** |
| Gene | - | 12 SNPs | 100% identical | 12 SNPs | 100% identical | 12 SNPs | 12 SNPs | 10 SNPs | 12 SNPs | 100% identical | 100% identical | 3 SNPs |
| Protein | - | - | - | 3 cs & 3 ncs aa subs | - | 3 cs & 3 ncs aa subs | 3 cs & 3 ncs aa subs | 3 cs & 3 cns aa subs | 3 cs & 3 ncs aa subs | - | - | 1 cs aa sub |
| Haplotype | H1 | H4 | H1 | H4 | H1 | H4 | H4 | H3 | H4 | H1 | H1 | H2 |
| **TaNAS4-A1** | **TraesCS5A03G1289600** | **TraesARI5A03G02826930** | **TraesJUL5A03G02803850** | **TraesLAC5A03G02740020** | **TraesNOR5A03G02809500** | **TraesSYM5A03G02817800** | **TraesMAC5A03G02783860** | **TraesTSP5A01G589500** | **TraesJAG5A03G02786450** | **TraesLDM5A03G02788100** | **TraesSTA5A03G02776250** | **5A:705403281-705402280 Reverse** |

| | | | | | | | | | | | | |
|---|---|---|---|---|---|---|---|---|---|---|---|---|
| Gene | - | 53 SNPs | 100% identical | 100% identical | 100% identical | 53 SNPs | 100% identical | 100% identical | 1 SNP→24 nt ext (3') | 1 SNP→24 nt ext (3') | 1 SNP→24 nt ext (3') | 1 SNP |
| Protein | - | 7 ncs subs | - | - | - | 7 ncs subs | - | - | 8 aa ext (C-ter) | 8 aa ext (C-ter) | 8 aa ext (C-ter) | 1 ncs aa sub |
| Haplotype | H1 | H4 | H1 | H1 | H1 | H4 | H1 | H1 | H3 | H3 | H3 | H2 |
| **TaNAS4-A2** | **TraesCS5A03G1289100** | **TraesARI5A03G02826980** | **TraesJUL5A03G02803790** | **TraesLAC5A03G02740080** | **TraesNOR5A03G02809440** | **TraesSYM5A03G02817780** | **TraesMAC5A03G02783780** | **TraesTSP5A01G589900** | **TraesJAG5A03G02786390** | **TraesLDM5A03G02788040** | **TraesSTA5A03G02776180** | **5A:705349537-705348536 Reverse** |
| Gene | - | 27 SNPs | 100% identical | 100% identical | 100% identical | 27 SNPs | 100% identical | 100% identical | 100% identical | 100% identical | 100% identical | 2 SNPs |
| Protein | - | 3 cs & 4 ncs aa subs | - | - | - | 3 cs & 4 ncs aa subs | - | - | - | - | - | Identical |
| Haplotype | H1 | H3 | H1 | H1 | H1 | H3 | H1 | H1 | H1 | H1 | H1 | H2 |
| **TaNAS4-D1** | **4D:513216136-513217137** | **TraesARIUn03G04594360** | **TraesJULUn03G04549510** | **TraesLACUn03G04460380** | **TraesNORUn03G04558740** | **scaffold_corrected_130-1:4924198-4925199** | **TraesMACUn03G04511200** | **TraesTSP4D01G404100** | **TraesJAG1D03G00574960** | **TraesLDM6B03G03624440** | **TraesSTAUn03G04532820** | **Un:108614518-108613517 Reverse** |
| Gene | - | 27 SNPs | 27 SNPs | 27 SNPs | 100% identical | 27 SNPs | 27 SNPs | 28 SNPs | 27 SNPs | 27 SNPs | 27 SNPs | 100% identical |
| Protein | - | 3 cs & 6 ncs aa subs | 3 cs & 6 ncs aa subs | 3 cs & 6 ncs aa subs | - | 3 cs & 6 ncs aa subs | 3 cs & 6 ncs aa subs | 3 cs & 7 ncs aa subs | 3 cs & 6 ncs aa subs | 3 cs & 6 ncs aa subs | 3 cs & 6 ncs aa subs | - |
| Haplotype | H2 | H1 | H1 | H1 | H2 | H1 | H1 | H3 | H1 | H1 | H1 | H2 |
| **TaNAS4-D2** | **4D:513233756-513234757** | **TraesARIUn03G04594330** | **TraesJULUn03G04549540** | **TraesLACUn03G04460410** | **TraesNORUn03G04558770** | **scaffold_corrected_130-1:4906770-4907771** | **TraesMACUn03G04511170** | **TraesTSP4D01G404400** | **TraesJAG1D03G00574990** | **TraesLDM6B03G03624480** | **TraesSTAUn03G04532850** | **Un:108595897-108596898 Forward** |
| Gene | | 31 SNPs | 31 SNPs | 31 SNPs | 100% identical | 31 SNPs | 31 SNPs | 31 SNPs | 31 SNPs | 31 SNPs | 31 SNPs | 8 SNPs |
| Protein | | 1 cs & 3 ncs aa subs | 1 cs & 3 ncs aa subs | 1 cs & 3 ncs aa subs | | 1 cs & 3 ncs aa subs | 1 cs & 3 ncs aa subs | 1 cs & 3 ncs aa subs | 1 cs & 3 ncs aa subs | 1 cs & 3 ncs aa subs | 1 cs & 3 ncs aa subs | 1 cs & 2 ncs aa subs |

| | | | | | | | | | | | | |
|---|---|---|---|---|---|---|---|---|---|---|---|---|
| Haplotype | H2 | H1 | H1 | H1 | H2 | H1 | H1 | H1 | H1 | H1 | H1 | H3 |
| **TaNAS5-B1** | **TraesCS3B03G154800** | **TraesARI3B03G01580550** | **TraesJUL3B03G01569870** | **TraesLAC3B03G01499150** | **TraesNOR3B03G01579440** | **TraesSYM3B03G01580660** | **TraesMAC3B03G01556940** | **TraesTSP3B01G085300** | **TraesJAG3B03G01566210** | **TraesLDM3B03G01558270** | **TraesSTA3B03G01550190** | **3B:40773334-40774332 Forward** |
| Gene | - | 456 nt ext (5') | 456 nt ext (5') | 456 nt ext (5') | 456 nt ext (5') | 456 nt ext (5') | 456 nt ext (5') | 100% identical | 456 nt ext (5') | 456 nt ext (5'), 1 SNP | 456 nt ext (5'), 1 SNP | 456 nt ext (5') |
| Protein | - | 152 aa ext (N-ter) | 152 aa ext (N-ter) | 152 aa ext (N-ter) | 152 aa ext (N-ter) | 152 aa ext (N-ter) | 152 aa ext (N-ter) | - | 152 aa ext (N-ter) | 152 aa ext (N-ter) | 152 aa ext (N-ter) | 152 aa ext (N-ter) |
| Haplotype | H2 | H1 | H1 | H1 | H1 | H1 | H1 | H2 | H1 | H3 | H3 | H1 |
| **TaNAS5-B2** | **TraesCS3B03G0154700** | **TraesARI3B03G01580480** | **TraesJUL3B03G01569780** | **TraesLAC3B03G01499070** | **TraesNOR3B03G01579350** | **TraesSYM3B03G01580590** | **TraesMAC3B03G01556850** | **TraesTSP3B01G085100** | **TraesJAG3B03G01566110** | **TraesLDM3B03G01558180** | **TraesSTA3B03G01550090** | **3B:40535199-40535741 Forward** |
| Gene | - | 1 SNP | 1 SNP | 1 SNP | 1 SNP | 1 SNP | 1 SNP | 1 SNP | 1 SNP | 16 SNPs & 57 nt ext (3') | 16 SNPs & 57 nt ext (3') | 1 SNP |
| Protein | - | 1 ncs aa sub | 1 ncs aa sub | 1 ncs aa sub | 1 ncs aa sub | 1 ncs aa sub | 1 ncs aa sub | 1 ncs aa sub | 1 ncs aa sub | 8 ncs subs, 19 aa ext (C-ter) | 8 ncs subs, 19 aa ext (C-ter) | 1 ncs aa sub |
| Haplotype | H2 | H1 | H1 | H1 | H1 | H1 | H1 | H1 | H1 | H3 | H3 | H1 |
| **TaNAS6-A1** | **TraesCS4A03G249100LC** | **TraesARI4A03G02082110** | **TraesJUL4A03G02065240** | **TraesLAC4A03G02000050** | **TraesNOR4A03G02073110** | **TraesSYM4A03G02072240** | **TraesMAC4A03G02046010** | **4A:148417599-148419883** | **TraesJAG4A03G02053440** | **TraesLDM4A03G02045130** | **TraesSTA4A03G02042380** | **4A:149046701-149047678 Forward** |
| Gene | 348 nt trun (5') | 100% identical | 100% identical | 1 SNP | 140→95 nt sub (3') | 100% identical | 140→95 nt sub (3') | 100% identical | 140→95 nt sub (3') | 100% identical | 140→95 nt sub (3') | 100% identical |
| Protein | 116 aa trun (N-ter) | - | - | 1 ncs aa sub | 46→31 aa (C-ter) | - | 1 ncs aa sub, 46→31 aa (C-ter) | - | 46→31 aa (C-ter) | - | 46→31 aa (C-ter) | - |
| Haplotype | H5 | H1 | H1 | H2 | H3 | H1 | H4 | H1 | H3 | H1 | H3 | H1 |
| **TaNAS6-B1** | **TraesCS4B03G520200** | **TraesARI4B03G02364430** | **TraesJUL4B03G02345670** | **TraesLAC4B03G02280440** | **TraesNOR4B03G02343820** | **TraesSYM4B03G02353360** | **TraesMAC4B03G02326520** | **TraesTSP4B01G198800** | **TraesJAG4B03G02324730** | **TraesLDM4B03G02327870** | **TraesSTA4B03G02322000** | **4B:403254329-403253337 Reverse** |

| | | | | | | | | | | | | |
|---|---|---|---|---|---|---|---|---|---|---|---|---|
| Gene | - | 100% identical | 100% identical | 100% identical | 100% identical | 100% identical | 100% identical | 100% identical | 100% identical | 100% identical | 100% identical | 100% identical |
| Protein | - | - | - | - | - | - | - | - | - | - | - | - |
| Haplotype | H1 | H1 | H1 | H1 | H1 | H1 | H1 | H1 | H1 | H1 | H1 | H1 |
| **TaNAS6-D1** | **TraesCS4D03G458900** | **TraesARI4D03G02539330** | **TraesJUL4D03G02519460** | **TraesLAC4D03G02453710** | **TraesNOR4D03G02517730** | **TraesSYM4D03G02528210** | **TraesMAC4D03G02498320** | **TraesTSP4D01G198500** | **TraesJAG4D03G02497690** | **TraesLDM4D03G02502750** | **TraesSTA4D03G02495500** | **4D:323098111-323096957 Reverse** |
| Gene | - | 162 nt trun (5') | 162 nt trun (5') | 162 nt trun (5') | 162 nt trun (5') | 162 nt trun (5') | 162 nt trun (5') | 100% identical | 162 nt trun (5') | 162 nt trun (5') | 162 nt trun (5') | 162 nt trun (5') |
| Protein | - | 54 aa trun (N-ter) | 54 aa trun (N-ter) | 54 aa trun (N-ter) | 54 aa trun (N-ter) | 54 aa trun (N-ter) | 54 aa trun (N-ter) | - | 54 aa trun (N-ter) | 54 aa trun (N-ter) | 54 aa trun (N-ter) | 54 aa trun (N-ter) |
| Haplotype | H2 | H1 | H1 | H1 | H1 | H1 | H1 | H2 | H1 | H1 | H1 | H1 |
| **TaNAS7-A1** | **TraesCS6A03G214000** | **TraesARI6A03G03221180** | **TraesJUL6A03G03292100** | **TraesLAC6A03G03220000** | **TraesNOR6A03G03297080** | **TraesSYM6A03G03206290** | **TraesMAC6A03G03265460** | **-** | **TraesJAGUn03G04499290** | **TraesLDM6A03G03269390** | **TraesSTA6A03G03256200** | **6A:60973173-60972181 Reverse** |
| Gene | - | 2 SNPs | 2 SNPs | 100% identical | 100% identical | 2 SNPs | 100% identical | - | 100% identical | 100% identical | 100% identical | 100% identical |
| Protein | - | 1 cs aa sub | 1 cs aa sub | - | - | 1 cs aa sub | - | - | - | - | - | - |
| Haplotype | H1 | H2 | H2 | H1 | H1 | H2 | H1 | - | H1 | H1 | H1 | H1 |
| **TaNAS7-A2** | **TraesCS6A03G973000** | **TraesARI6A03G03362910** | **TraesJUL6A03G03432260** | **TraesLAC6A03G03362370** | **TraesNOR6A03G03439190** | **TraesSYM6A03G03348250** | **TraesMAC6A03G03404280** | **TraesTSP6A01G374500** | **TraesJAG6A03G03398040** | **TraesLDM6A03G03408800** | **TraesSTA6A03G03395830** | **6A:603249197-603250187 Forward** |
| Gene | - | 100% identical | 100% identical | 17 SNPs | 100% identical | 100% identical | 100% identical | 2 SNPs | 100% identical | 100% identical | 100% identical | 16 SNPs |
| Protein | - | - | - | 5 cs & 1 ncs aa sub | - | - | - | 1 ncs aa sub | - | - | - | 5 cs & 1 ncs sub |
| Haplotype | H1 | H1 | H1 | H4 | H1 | H1 | H1 | H2 | H1 | H1 | H1 | H3 |
| **TaNAS7-B1** | **TraesCS6B03G1190400** | **TraesARI6B03G03586430** | **TraesJUL6B03G03656530** | **TraesLAC6B03G03580460** | **TraesNOR6B03G03662980** | **TraesSYM6B03G03568520** | **TraesMAC6B03G03626210** | **TraesTSP6B01G462100** | **TraesJAG6B03G03616290** | **TraesLDM6B03G03626760** | **TraesSTA6B03G03614750** | **6B:694258986-694259978 Forward** |

| | | | | | | | | | | | | |
|---|---|---|---|---|---|---|---|---|---|---|---|---|
| Gene | - | 7 SNPs | 5 SNPs | 43 SNPs | 5 SNPs | 5 SNPs | 43 SNPs | 43 SNPs | 5 SNPs | 5 SNPs | 100% identical | 5 SNPs |
| Protein | - | 1 cs & 1 ncs aa sub | 1 ncs aa sub | 7 cs & 10 ncs aa subs | 1 ncs aa sub | 1 ncs aa sub | 7 cs & 10 ncs aa subs | 7 cs & 10 ncs aa subs | 1 ncs aa sub | 1 ncs aa sub | - | 1 ncs aa sub |
| Haplotype | H3 | H4 | H1 | H2 | H1 | H1 | H2 | H2 | H1 | H1 | H3 | H1 |
| **TaNAS7-D1** | **TraesCS6D03G854100** | **TraesARI6D03G03755720** | **TraesJUL6D03G03824340** | **TraesLAC6D03G03741940** | **TraesNOR6D03G03831960** | **TraesSYM6D03G03739130** | **TraesMAC6D03G03789530** | **TraesTSP6D01G398500** | **TraesJAG6D03G03774220** | **TraesLDM6D03G03795250** | **TraesSTA6D03G03784240** | **6D:456540534-456541526 Forward** |
| Gene | - | 100% identical | 100% identical | 4 SNPs | 100% identical | 4 SNPs | 100% identical | 1 SNP | 100% identical | 100% identical | 100% identical | 4 SNPs |
| Protein | - | - | - | 1 cs & 2 ncs aa subs | - | 1 cs & 2 ncs aa subs | - | Identical | - | - | - | 1 cs & 2 ncs aa subs |
| Haplotype | H1 | H1 | H1 | H2 | H1 | H2 | H1 | H3 | H1 | H1 | H1 | H2 |
| **TaNAS8-A1** | **TraesCS6A03G1000600** | **TraesARI6A03G03366660** | **TraesJUL6A03G03435950** | **TraesLAC6A03G03365270** | **TraesNOR6A03G03443150** | **TraesSYM6A03G03352220** | **TraesMAC6A03G03408270** | **TraesTSP6A01G387500** | **TraesJAG6A03G03401930** | **TraesLDM6A03G03412690** | **TraesSTA6A03G03399820** | **6A:609149647-609148661 Reverse** |
| Gene | - | 16 SNPs | 16 SNPs | 16 SNPs | 100% identical | 16 SNPs | 16 SNPs | 100% identical | 16 SNPs | 16 SNPs | 16 SNPs | 3 SNPs |
| Protein | - | 2 cs & 2 ncs aa subs | 2 cs & 2 ncs aa subs | 2 cs & 2 ncs aa subs | - | 2 cs & 2 ncs aa subs | 2 cs & 2 ncs aa subs | - | 2 cs & 2 ncs aa subs | 2 cs & 2 ncs aa subs | 2 cs & 2 ncs aa subs | 2 cs & 1 ncs aa subs |
| Haplotype | H2 | H1 | H1 | H1 | H2 | H1 | H1 | H2 | H1 | H1 | H1 | H3 |
| **TaNAS8-B1** | **TraesCS6B03G1227100** | **TraesARI6B03G03592370** | **TraesJUL6B03G03662750** | **TraesLAC6B03G03583300** | **TraesNOR6B03G03669280** | **TraesSYM6B03G03574130** | **TraesMAC6B03G03629250** | **TraesTSP6B01G478100** | **TraesJAG6B03G03622080** | **TraesLDM6B03G03631140** | **TraesSTA6B03G03620490** | **6B:704833079-704832093 Reverse** |
| Gene | - | 100% identical | 100% identical | 33 SNPs, 3 nt ins | 100% identical | 34 SNPs, 3 nt ins | 100% identical | 31 SNPs, 3 nt ins | 31 SNPs, 3 nt ins | 34 SNPs, 3 nt ins | 100% identical | 100% identical |
| Protein | - | - | - | 2 cs & 7 ncs aa subs, 1 aa ins | - | 2 cs & 10 ncs aa subs, 1 aa ins, 1 aa del | - | 1 cs & 7 ncs aa subs, 1 aa ins | 1 cs & 7 ncs aa subs, 1 aa ins | 2 cs & 10 ncs aa subs, 1 aa ins, 1 aa del | - | - |
| Haplotype | H1 | H1 | H1 | H4 | H1 | H3 | H1 | H2 | H2 | H3 | H1 | H1 |

| **TaNAS8-D1** | **TraesCS6D03G881800** | **TraesARI6D03G03764430** | **TraesJUL6D03G03828250** | **TraesLAC6D03G03745810** | **TraesNOR6D03G03835840** | **TraesSYM6D03G03742780** | **TraesMAC6D03G03793230** | **TraesTSP6D01G412500** | **TraesJAG6D03G03777380** | **TraesLDM6D03G03799190** | **TraesSTA6D03G03787970** | **6D:462354308-462355294 Forward** |
|---|---|---|---|---|---|---|---|---|---|---|---|---|
| Gene | - | 100% identical | 100% identical | 33 SNPs | 100% identical | 100% identical | 33 SNPs | 33 SNPs | 33 SNPs | 100% identical | 33 SNPs | 26 SNPs |
| Protein | - | - | - | 7 cs & 6 ncs aa subs | - | - | 7 cs & 6 ncs aa subs | 7 cs & 6 ncs aa subs | 7 cs & 6 ncs aa subs | - | 7 cs & 6 ncs aa subs | 6 cs & 4 ncs aa subs |
| Haplotype | H1 | H1 | H1 | H2 | H1 | H1 | H2 | H2 | H2 | H1 | H2 | H3 |
| **TaNAS9-A1** | **TraesCS2A03G196300** | **TraesARI2A03G00615350** | **TraesJUL2A03G00611970** | **TraesLAC2A03G00612520** | **TraesNOR2A03G00616330** | **TraesSYM2A03G00614950** | **TraesMAC2A03G00607040** | **TraesTSP2A01G104200** | **TraesJAG2A03G00607290** | **TraesLDM2A03G00610790** | **TraesSTA2A03G00607220** | **2A:49221108-49222130 Forward** |
| Gene | - | 100% identical | 100% identical | 100% identical | 100% identical | 100% identical | 100% identical | 100% identical | 33 nt 5' trunation | 100% identical | 100% identical | 100% identical |
| Protein | - | - | - | - | - | - | - | - | 11 aa N-term trunation | - | - | - |
| Haplotype | H1 | H1 | H1 | H1 | H1 | H1 | H1 | H1 | H2 | H1 | H1 | H1 |
| **TaNAS9-B1** | **TraesCS2B03G264600** | **TraesARI2B03G00868570** | **TraesJUL2B03G00862760** | **TraesLAC2B03G00854580** | **TraesNOR2B03G00869150** | **TraesSYM2B03G00868510** | **TraesMAC2B03G00856020** | **TraesTSP2B01G123800** | **TraesJAG2B03G00857470** | **TraesLDM2B03G00859510** | **TraesSTA2B03G00858670** | **2B:72895158-72896180 Forward** |
| Gene | - | 19 SNPs | 18 SNPs | 100% identical | 100% identical | 18 SNPs | 100% identical | 100% identical | 100% identical | 100% identical | 100% identical | 100% identical |
| Protein | - | 4 cs & 2 ncs aa subs | 4 cs & 2 ncs aa subs | - | - | 4 cs & 2 ncs aa subs | - | - | - | - | - | - |
| Haplotype | H1 | H3 | H2 | H1 | H1 | H2 | H1 | H1 | H1 | H1 | H1 | H1 |
| **TaNAS9-D1** | **TraesCS2D03G196500** | **TraesARI2D03G01126000** | **TraesJUL2D03G01115740** | **TraesLAC2D03G01061350** | **TraesNOR2D03G01125590** | **TraesSYM2D03G01123730** | **TraesMAC2D03G01108120** | **TraesTSP2D01G110200** | **TraesJAG2D03G01112430** | **TraesLDM2D03G01110830** | **TraesSTA2D03G01098320** | **2D:45799198-45800220 Forward** |
| Gene | - | 100% identical | 100% identical | 100% identical | 100% identical | 33 nt trun (5') | 100% identical | 100% identical | 100% identical | 33 nt trun (5') | 33 nt trun (5') | 100% identical |
| Protein | - | - | - | - | - | 11 aa trun (N-ter) | - | - | - | 11 aa trun (N-ter) | 11 aa trun (N-ter) | - |

| | | | | | | | | | | | | |
|---|---|---|---|---|---|---|---|---|---|---|---|---|
| Haplotype | H1 | H1 | H1 | H1 | H1 | H2 | H1 | H1 | H1 | H2 | H2 | H1 |
| **TaNAS10-A1** | **TraesCS3A03G1088500** | **TraesARI3A03G01521050** | **TraesJUL3A03G01511990** | **TraesLAC3A03G01443470** | **TraesNOR3A03G01520940** | **TraesSYM3A03G01522450** | **TraesMAC3A03G01497740** | **TraesTSP3A01G496900** | **TraesJAG3A03G01508120** | **TraesLDM3A03G01499820** | **TraesSTA3A03G01491250** | **3A:701211258-701210236 Reverse** |
| Gene | - | 100% identical | 2 SNPs | 100% identical | 100% identical | 100% identical | 100% identical | 2 SNPs | 100% identical | 100% identical | 100% identical | 100% identical |
| Protein | - | - | 1 cs aa sub | - | - | - | - | 2 ncs aa subs | - | - | - | - |
| Haplotype | H1 | H1 | H2 | H1 | H1 | H1 | H1 | H3 | H1 | H1 | H1 | H1 |
| | Chinese Spring | ArinaLrFor | Julius | Lancer | Norin61 | SY Mattis | Mace | Triticum spelta | Jagger | Landmark | Stanley | Fielder |
| **TaNAS2-B3** | - | - | **TraesJUL6B03G03521280** | - | - | **TraesSYM6B03G03437270** | - | - | **TraesJAG6B03G03488360** | - | **TraesSTA6B03G03488710** | - |
| Gene | - | - | - | - | - | - | - | - | 1 SNP | - | 1 SNP | - |
| Protein | - | - | - | - | - | - | - | - | 1 cs aa sub | - | 1 cs aa sub | - |
| Haplotype | - | - | H1 | - | - | H1 | - | - | H2 | - | H2 | - |
| **TaNAS4-A3** | - | **TraesARI5A03G02826920** | - | - | - | **TraesSYM5A03G02817720** | - | - | - | - | - | - |
| Gene | - | - | - | - | - | - | - | - | - | - | - | - |
| Protein | - | - | - | - | - | - | - | - | - | - | - | - |
| Haplotype | - | H1 | - | - | - | H1 | - | - | - | - | - | - |
| **TaNAS5-B3** | - | **TraesARI3B03G01580520** | **TraesJUL3B03G01569830** | **TraesLAC3B03G01499120** | **TraesNOR3B03G01579400** | **TraesSYM3B03G01580630** | **TraesMAC3B03G01556900** | **TraesTSP3B01G085200** | **TraesJAG3B03G01566170** | **TraesLDM3B03G01558230** | **TraesSTA3B03G01550140** | **3B:40569853-40570395 Forward** |
| Gene | - | - | 105 nt ext (5') | - | - | - | - | - | - | 7 SNPs | 7 SNPs | - |
| Protein | - | - | 35 aa ext (N-ter) | - | - | - | - | - | - | 2 cs & 2 ncs aa subs | 2 cs & 2 ncs aa subs | - |
| Haplotype | - | H1 | H3 | H1 | H1 | H1 | H1 | H1 | H1 | H2 | H2 | H1 |

| | | | | | | | | | | | | |
|---|---|---|---|---|---|---|---|---|---|---|---|---|
| **TaNAS8-B2** | **-** | **-** | **-** | **-** | **-** | **TraesSYM6B03G03574110** | **-** | **-** | **-** | **-** | **-** | **-** |
| Gene | - | - | - | - | - | - | - | - | - | - | - | - |
| Protein | - | - | - | - | - | - | - | - | - | - | - | - |
| Haplotype | - | - | - | - | - | H1 | - | - | - | - | - | - |

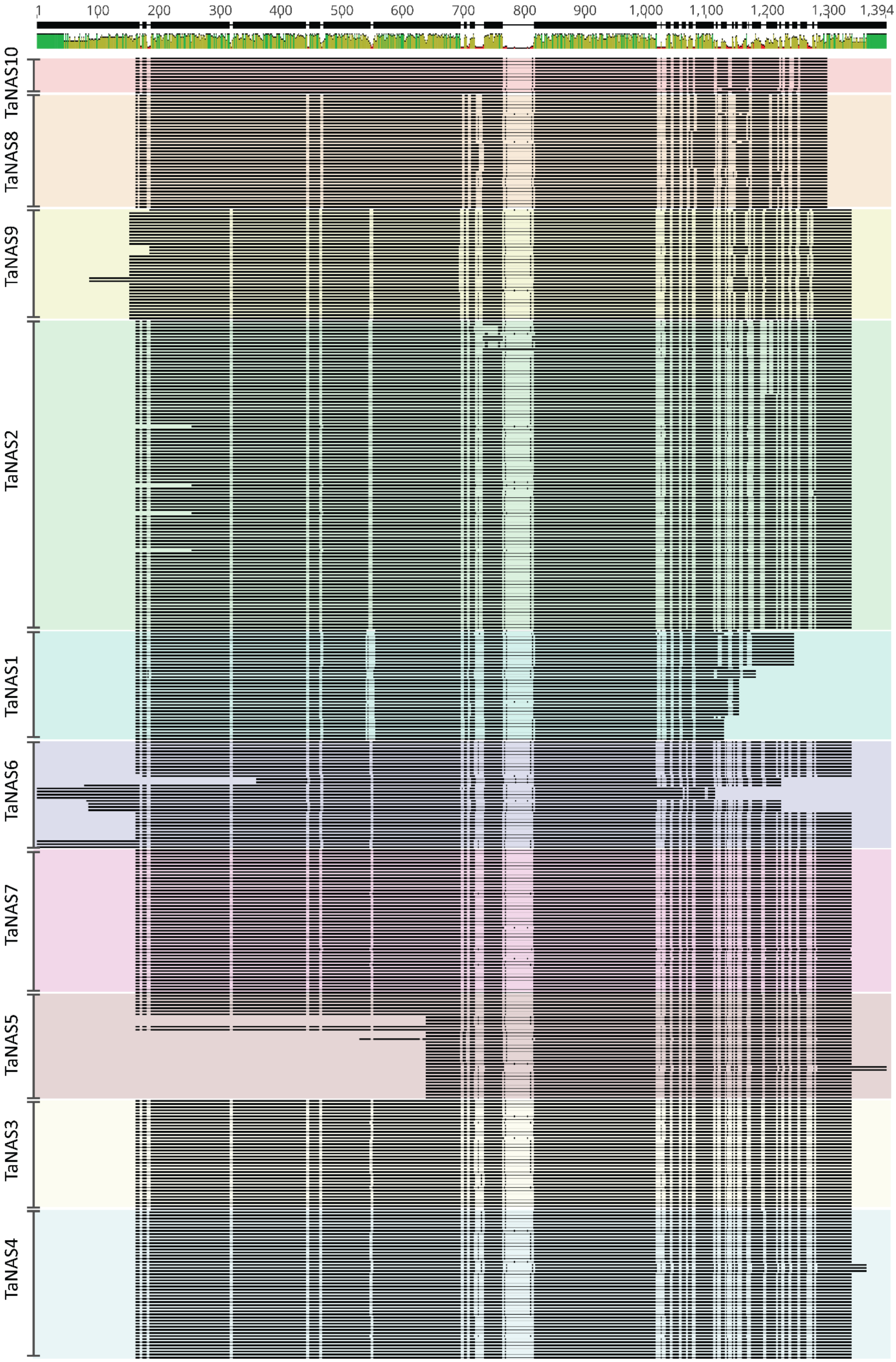
1
100
200
300
400
500
600
700
800
900
1,000
1,100
1,200
1,300
1,394
TaNAS10
TaNAS8
TaNAS9
TaNAS2
TaNAS1
TaNAS6
TaNAS7
TaNAS5
TaNAS3
TaNAS4

**Figure S1. Alignment of 425 *TaNAS* coding sequences from cvs. Chinese Spring, ArinaLrFor, Julius, Lancer, Norin61, SY Mattis, Mace, Jagger, Landmark, Stanley, Fielder, and *Triticum spelta* L.** The alignment was performed using Geneious alignment tool. The top bar indicates consensus (black bars) and high (green), medium (yellow), and low (red) identity indicated as a histogram. The nucleotide positions indicated. Each coloured segment represents a *TaNAS* grouping as determined by the PhyML Geneious plugin with the K80 substitution model and 100 bootstrap replications.

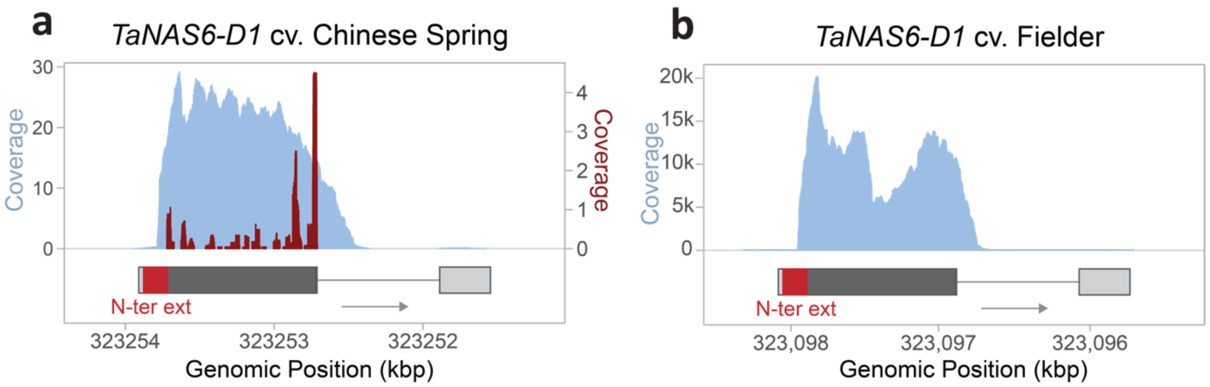


**Figure S2. Ribosomal profile and RNA-seq coverage of *TaNAS6-D1*.** Histograms of overlapping Ribo-seq (red; right y-axis) and RNA-seq (blue; left y-axis) of *TaNAS6-D1* homoeologs in (a) developing grain five days post anthesis from cv. Chinese Spring and (b) roots of cv. Fielder 11 days after Fe deficiency. Coverage in (a) is normalised to reads per million mapped to nuclear coding sequences, with data sourced from (Guo et al., 2023), while (b) shows the raw read count mapped to nuclear coding sequences produced within this study. Arrows indicate gene orientation (5' to 3'). The *TaNAS6-D1* gene model is predicted by IWGSC Refseq2.0 with the 5' and 3' UTRs (light grey rectangles), coding sequences (dark grey rectangles), and introns (horizontal lines) indicated. The 162 bp N-terminus extension (red rectangle) predicted in the cv. Chinese Spring *TaNAS6-D1* gene by the IWGSC Refseq1.0 and 2.0 annotation is highlighted.

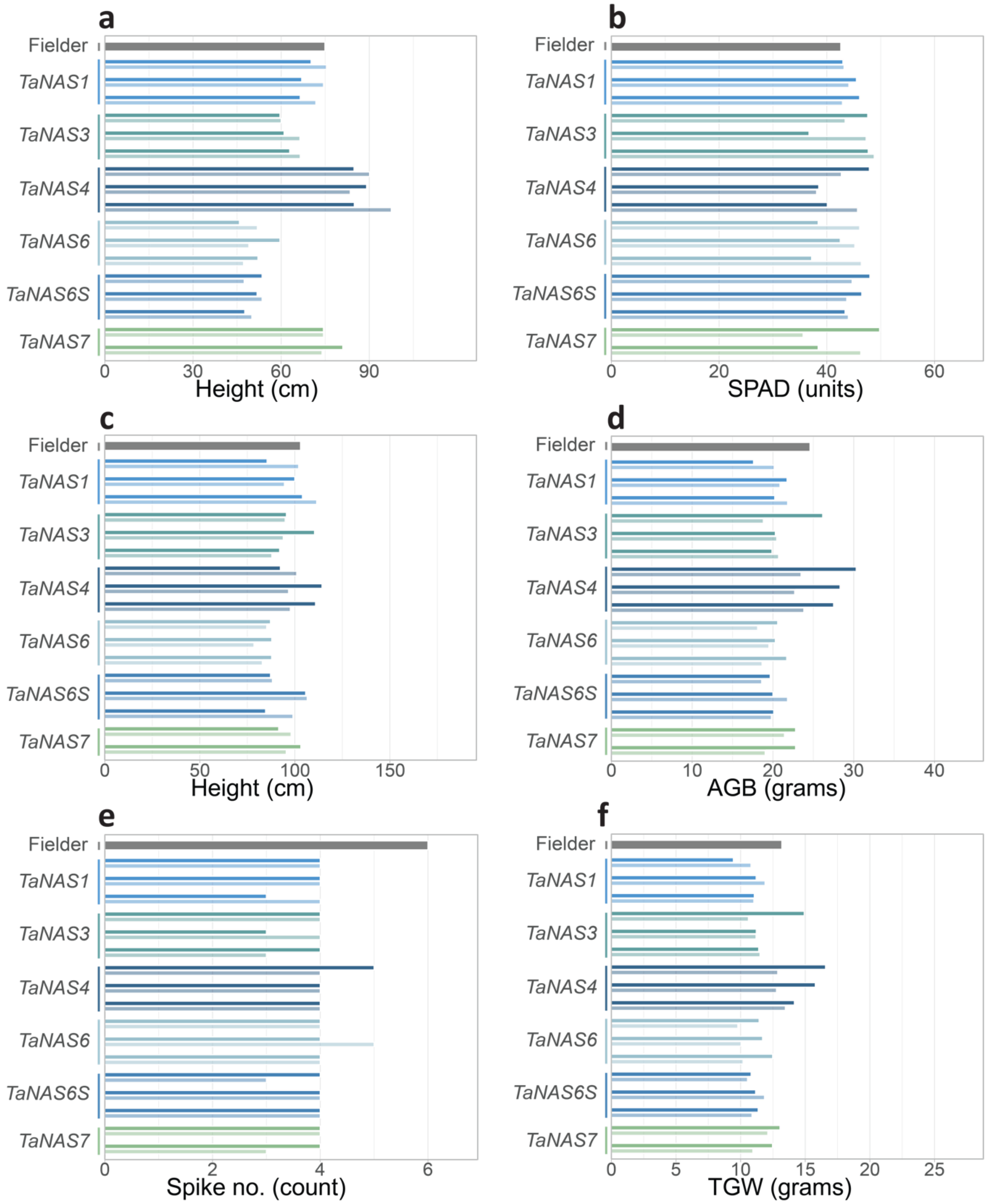

a
Fielder
TaNAS1
TaNAS3
TaNAS4
TaNAS6
TaNAS6S
TaNAS7
0
30
60
90
Height (cm)
b
Fielder
TaNAS1
TaNAS3
TaNAS4
TaNAS6
TaNAS6S
TaNAS7
0
20
40
60
SPAD (units)
c
Fielder
TaNAS1
TaNAS3
TaNAS4
TaNAS6
TaNAS6S
TaNAS7
0
50
100
150
Height (cm)
d
Fielder
TaNAS1
TaNAS3
TaNAS4
TaNAS6
TaNAS6S
TaNAS7
0
10
20
30
40
AGB (grams)
e
Fielder
TaNAS1
TaNAS3
TaNAS4
TaNAS6
TaNAS6S
TaNAS7
0
2
4
6
Spike no. (count)
f
Fielder
TaNAS1
TaNAS3
TaNAS4
TaNAS6
TaNAS6S
TaNAS7
0
5
10
15
20
25
TGW (grams)


Fig S3. cont.

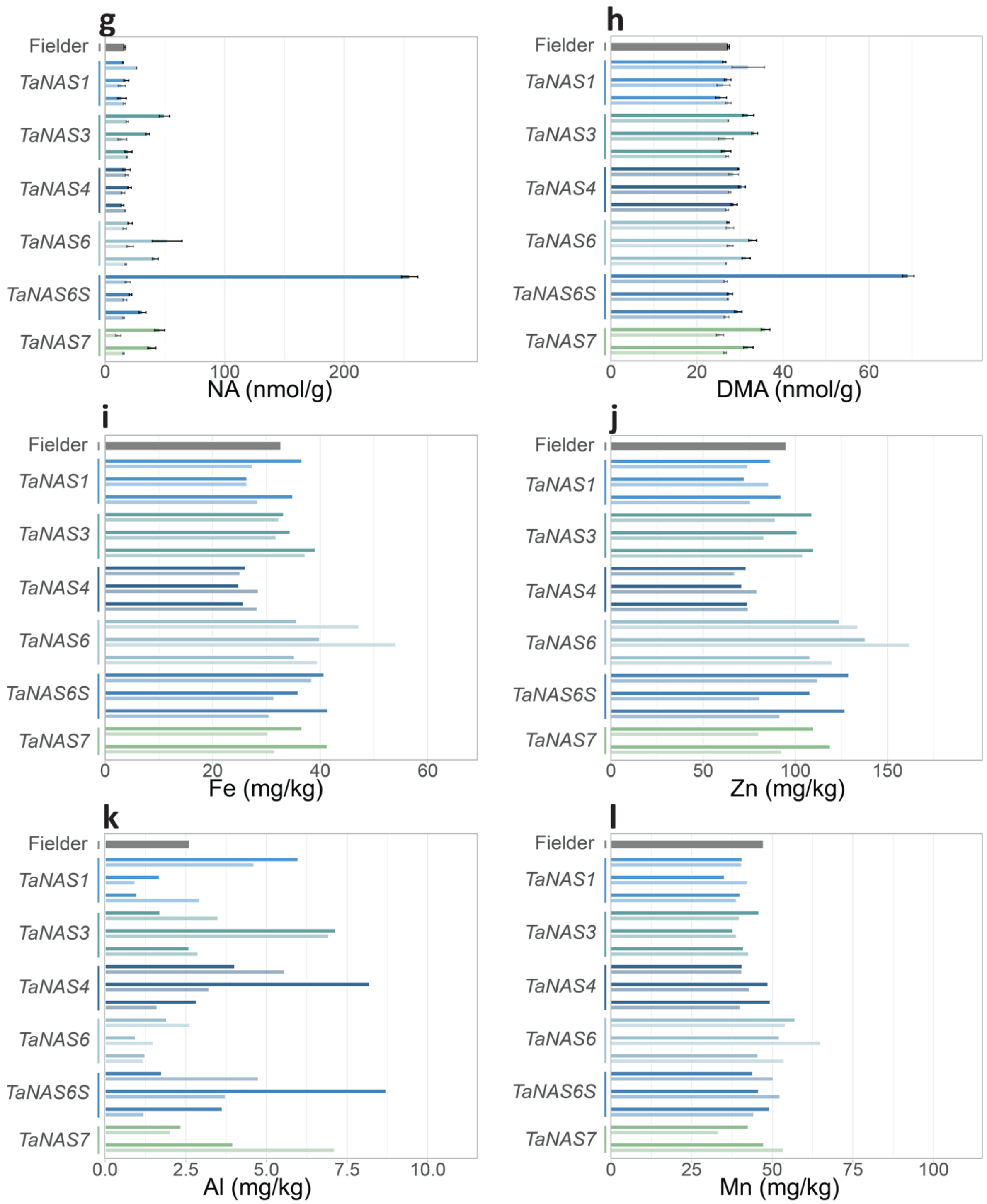


**Figure S3. Phenotypic and nutritional analysis of $T_1$ *TaNAS*-OE and NS plants in the glasshouse.** Plant (a) height (cm) and (b) SPAD of three month-old plants; (c) height (cm), (d) above ground biomass (AGB, g), (e) spike number, and (f) total grain weight (TGW, g) of mature plants; and grain (g) nicotianamine (NA, nmol/g), (h) 2'deoxymugeneic acid (DMA, nmol/g), (i) Fe (mg/kg), (j) Zn (mg/kg), (k) aluminium (Al, mg/kg), and (l) manganese (Mn, mg/kg) of *TaNAS1* (blue), *TaNAS3* (teal), *TaNAS4* (navy blue), *TaNAS6* (light blue), *TaNAS6S* (dark blue) and *TaNAS7* (green) lines, respective nulls (faded colour of each), and cv. Fielder (grey) in the glasshouse in 2021. Events 1, 2 and 3 for each *TaNAS*-OE line are shown at the top, middle and bottom, respectively (n=1). Error bars for NA and DMA concentrations represent the standard error of three technical replicates.

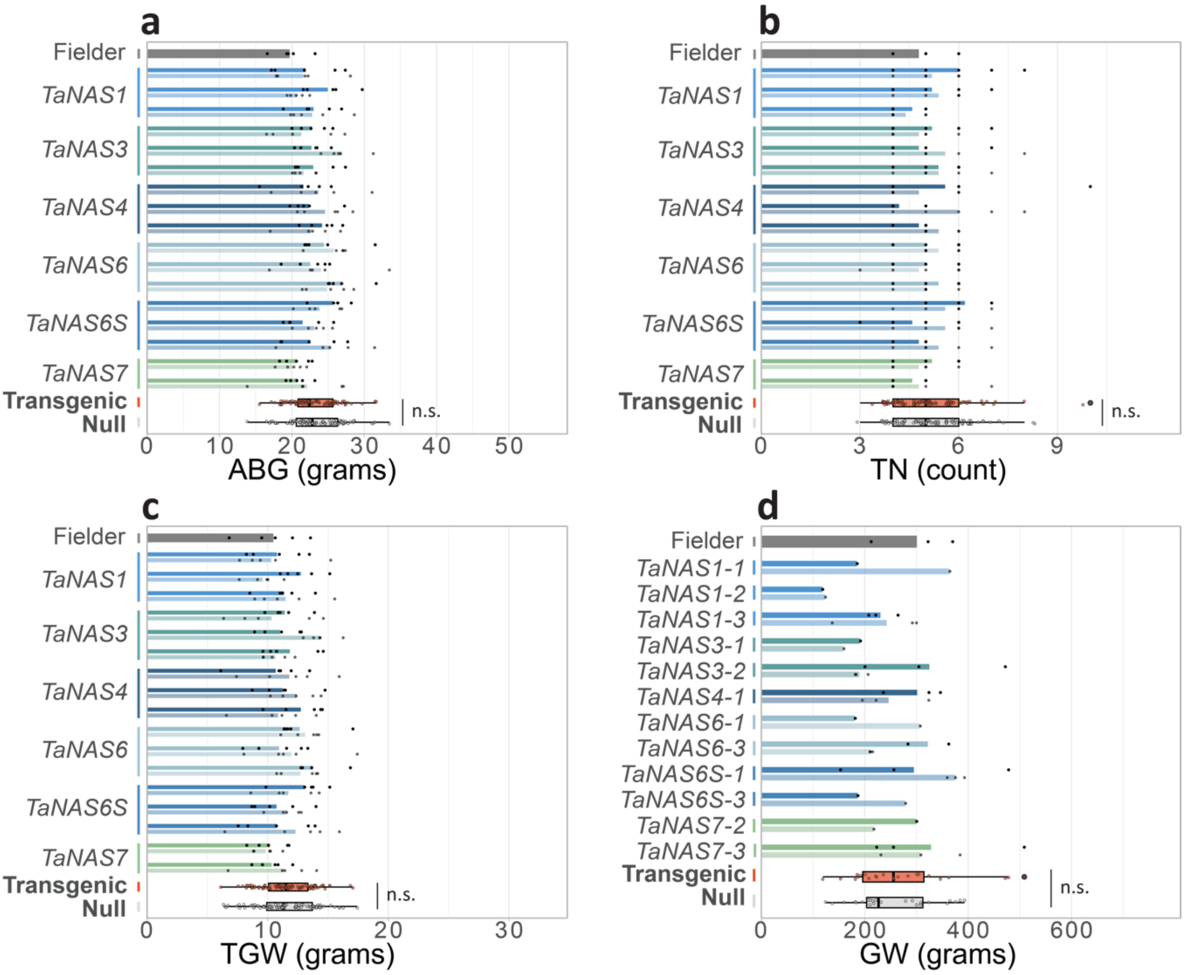


**Figure S4. Agronomic traits of T$_3$ *TaNAS*-OE and NS plants in the glasshouse and field.** Plant (a) above ground biomass (AGB, g), (b) tiller number per plant (TN/plant), (c) total grain weight (TGW, g) from glasshouse (GH) grown plants, and (d) subsampled grain weight (GW, g) of field grown plants of *TaNAS1* (blue), *TaNAS3* (teal), *TaNAS4* (navy blue), *TaNAS6* (light blue), *TaNAS6S* (dark blue) and *TaNAS7* (green) lines, respective nulls (faded colour of each), and cv. Fielder (grey) in the glasshouse in 2022. Events 1, 2 and 3 for each *TaNAS*-OE line are shown at the top, middle and bottom, respectively (n=5). Statistical differences between transgenic events and their respective NS are shown as p values determined by a two-sample Student's t-test assuming unequal variance with a 95% confidence interval. The bars represent the average of one, three, or five biological replicates with the raw datapoints overlaid. The box plots represent all transgenic (orange) values compared with all null (grey) values. Non-significant (n.s.) values between all transgenic and all null segregant values as determined by a two-sample Student's t-test assuming unequal variance are indicated.

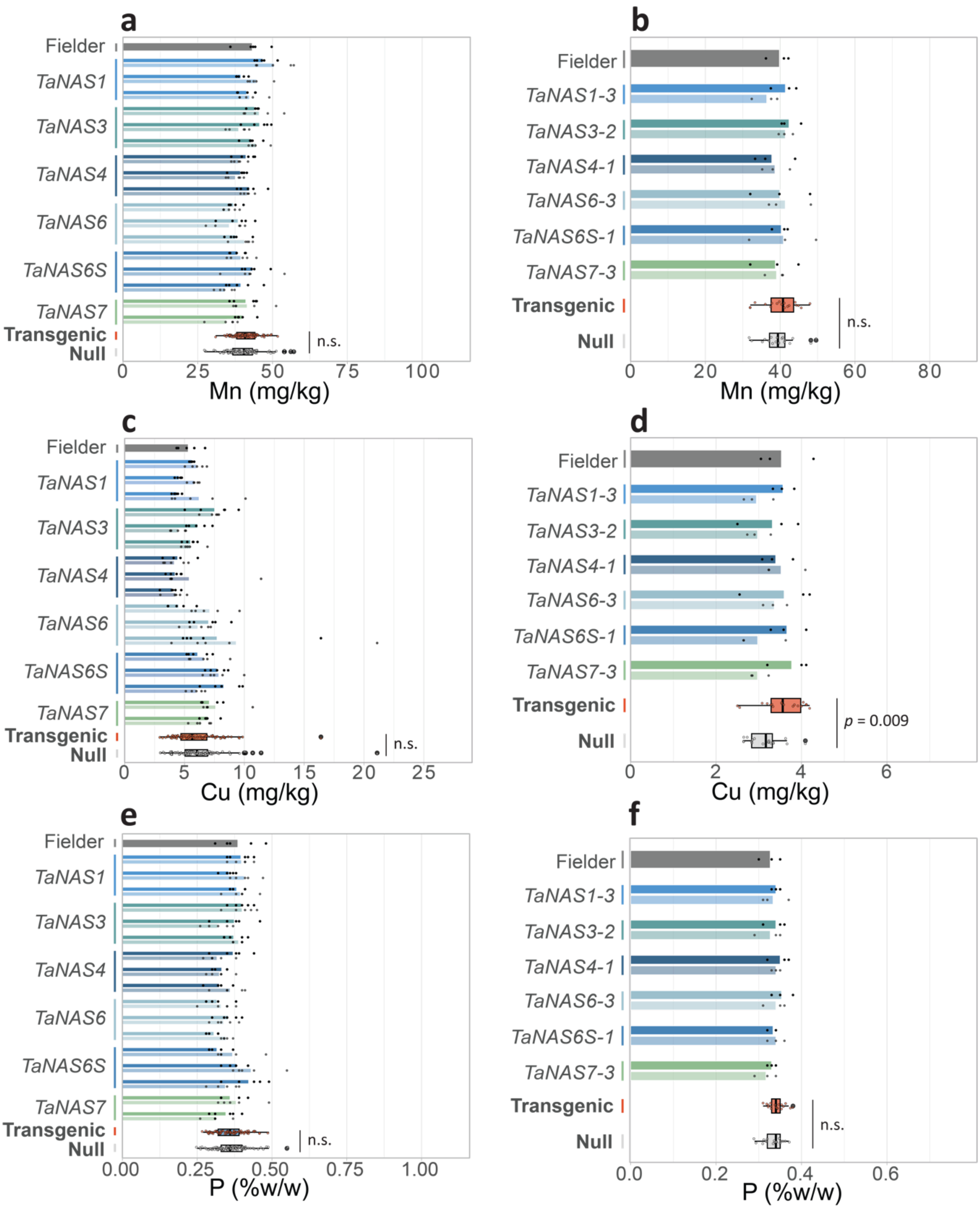


**Figure S5. Grain Mn, Cu, and P concentrations of T$_3$ *TaNAS*-OE and null plants in glasshouse and field experiments.** Grain (a, b) manganese (Mn, mg/kg), (c, d) copper (Cu, mg/kg), (e, f) phosphorus (P, %w/w) concentrations of *TaNAS1* (blue), *TaNAS3* (teal), *TaNAS4* (navy blue), *TaNAS6* (light blue), *TaNAS6S* (dark blue) and *TaNAS7* (green) lines, respective nulls (faded colour of each), and cv. Fielder (grey) in the glasshouse and field in 2022. For the glasshouse experiment (a, c, e) events 1, 2 and 3 for each *TaNAS*-OE line are shown at the top, middle and bottom, respectively (n=5). For the field experiment (b, d, f), events *TaNAS1-3*, *TaNAS3-2*, *TaNAS4-1*, *TaNAS6-3*, *TaNAS6S-1*, and *TaNAS7-3* are shown (n=3). Statistical differences between transgenic events and their respective NS are shown as *p* values determined by a two-sample Student's t-test assuming unequal variance with a 95%

confidence interval. The bars represent the average of three or five biological replicates with the raw datapoints overlaid. The box plots represent all transgenic (orange) values compared with all null (grey) values. Non-significant (n.s.) values between all transgenic and all null segregant values as determined by a two-sample Student's t-test assuming unequal variance are indicated.

**Table S4. Grain Fe, Zn, and NA fold-differences between *TaNAS*-OE and null plants at the $T_3$ and $T_4$ generations.** Fold differences (bold) in grain Fe, Zn and NA concentrations between *TaNAS*-OE events and their respective NS with p-values as determined by a two-sample Student's t-test assuming unequal variance shown in the adjacent column. Underlined values indicated statistically significant differences where $p < 0.05$.

| | **2022 glasshouse** | | | | **2022 field** | | | | **2023 field** | |
|---|---|---|---|---|---|---|---|---|---|---|
| **Event** | **Fe** | **p value** | **Zn** | ***p* value** | **Fe** | ***p* value** | **Zn** | ***p* value** | **NA** | ***p* value** |
| *TaNAS1-1* | **0.898** | 0.315 | **0.868** | 0.253 | | | | | **0.746** | 0.185 |
| *TaNAS1-2* | **0.909** | 0.278 | **0.898** | 0.156 | | | | | **0.832** | 0.430 |
| *TaNAS1-3* | **1.003** | 0.976 | **1.000** | 1.000 | **1.114** | 0.140 | **1.137** | 0.382 | **1.056** | 0.607 |
| *TaNAS3-1* | **1.248** | 0.050 | **1.145** | 0.146 | | | | | **1.247** | 0.222 |
| *TaNAS3-2* | **1.506** | 0.003 | **1.512** | 0.001 | **1.288** | 0.005 | **1.240** | 0.165 | **1.279** | 0.101 |
| *TaNAS3-3* | **1.121** | 0.145 | **1.166** | 0.027 | | | | | **0.980** | 0.902 |
| *TaNAS4-1* | **1.089** | 0.346 | **1.057** | 0.257 | **0.788** | 0.241 | **0.967** | 0.745 | **1.273** | 0.098 |
| *TaNAS4-2* | **1.068** | 0.363 | **1.044** | 0.517 | | | | | **0.980** | 0.847 |
| *TaNAS4-3* | **0.962** | 0.452 | **0.980** | 0.710 | | | | | **0.915** | 0.654 |
| *TaNAS6-1* | **0.973** | 0.746 | **1.177** | 0.013 | | | | | **1.361** | 0.129 |
| *TaNAS6-2* | **1.292** | 0.002 | **1.315** | 0.003 | | | | | **2.032** | 0.085 |
| *TaNAS6-3* | **0.840** | 0.462 | **1.113** | 0.010 | **1.092** | 0.226 | **1.289** | 0.206 | **1.555** | 0.081 |
| *TaNAS6S-1* | **1.038** | 0.728 | **1.196** | 0.025 | **1.217** | 0.132 | **1.396** | 0.015 | **3.702** | 0.013 |
| *TaNAS6S-2* | **1.109** | 0.225 | **1.255** | 0.018 | | | | | **0.924** | 0.495 |
| *TaNAS6S-3* | **1.550** | 0.020 | **1.746** | 0.000 | | | | | **1.364** | 0.008 |
| *TaNAS7-1* | **0.950** | 0.407 | **0.938** | 0.460 | **1.109** | 0.419 | **0.986** | 0.839 | **0.890** | 0.273 |
| *TaNAS7-2* | **1.238** | 0.037 | **1.309** | 0.017 | | | | | **0.950** | 0.612 |
| *TaNAS7-3* | **0.793** | 0.425 | **1.546** | 0.001 | **1.294** | 0.002 | **1.290** | 0.101 | **1.746** | 0.017 |

**Table S5. Grain P, Mn, and Cu fold-differences between *TaNAS*-OE and null plants at the $T_3$ generation.** The table contains fold differences (bolded) of transgenic and their respective NS with p-values as determined by a two-sample Student's t-test assuming unequal variance shown in the adjacent column. Underlined values indicated statistically significant differences where p < 0.05.

| | **2022 glasshouse** | | | | | | **2022 field** | | | | | |
|---|---|---|---|---|---|---|---|---|---|---|---|---|
| **Event** | **P** | ***p* value** | **Mn** | ***p* value** | **Cu** | ***p* value** | **P** | ***p* value** | **Mn** | ***p* value** | **Cu** | ***p* value** |
| *TaNAS1-1* | **0.995** | 0.934 | **0.927** | 0.264 | **0.931** | 0.265 | | | | | | |
| *TaNAS1-2* | **0.873** | 0.050 | **0.884** | 0.026 | **0.782** | 0.001 | | | | | | |
| *TaNAS1-3* | **0.970** | 0.632 | **0.977** | 0.685 | **0.696** | 0.173 | **1.020** | 0.760 | **1.137** | 0.158 | **1.212** | 0.076 |
| *TaNAS3-1* | **0.995** | 0.941 | **0.967** | 0.592 | **1.021** | 0.863 | | | | | | |
| *TaNAS3-2* | **1.176** | 0.143 | **1.182** | 0.020 | **1.368** | 0.012 | **1.041** | 0.610 | **1.024** | 0.642 | **1.117** | 0.509 |
| *TaNAS3-3* | **0.959** | 0.407 | **0.964** | 0.412 | **1.003** | 0.969 | | | | | | |
| *TaNAS4-1* | **1.171** | 0.122 | **1.065** | 0.193 | **1.068** | 0.675 | **1.029** | 0.590 | **0.978** | 0.841 | **0.965** | 0.747 |
| *TaNAS4-2* | **1.025** | 0.705 | **1.045** | 0.331 | **0.781** | 0.480 | | | | | | |
| *TaNAS4-3* | **0.900** | 0.218 | **1.018** | 0.717 | **0.932** | 0.557 | | | | | | |
| *TaNAS6-1* | **0.958** | 0.632 | **1.032** | 0.431 | **0.634** | 0.020 | | | | | | |
| *TaNAS6-2* | **1.047** | 0.529 | **1.085** | 0.408 | **1.146** | 0.302 | | | | | | |
| *TaNAS6-3* | **0.884** | 0.007 | **0.927** | 0.204 | **0.828** | 0.681 | **1.039** | 0.562 | **0.965** | 0.817 | **1.068** | 0.710 |
| *TaNAS6S-1* | **0.859** | 0.173 | **0.971** | 0.612 | **0.919** | 0.501 | **0.980** | 0.649 | **0.986** | 0.924 | **1.230** | 0.176 |
| *TaNAS6S-2* | **0.888** | 0.236 | **1.022** | 0.814 | **0.981** | 0.835 | | | | | | |
| *TaNAS6S-3* | **1.227** | 0.058 | **1.156** | 0.088 | **1.387** | 0.021 | | | | | | |
| *TaNAS7-1* | **0.881** | 0.200 | **0.951** | 0.519 | **0.913** | 0.314 | **1.000** | 1.000 | **1.006** | 0.967 | **0.913** | 0.373 |
| *TaNAS7-2* | **0.947** | 0.590 | **0.990** | 0.904 | **0.931** | 0.568 | | | | | | |
| *TaNAS7-3* | **1.109** | 0.238 | **1.153** | 0.059 | **1.088** | 0.252 | **1.042** | 0.465 | **0.991** | 0.940 | **1.269** | 0.091 |

**Table S6. Gene IDs and coding sequences used as queries to blast for novel bread wheat *NAS* genes.**

| NAS gene ID | Coding sequence |
|---|---|
| AT5G04<br>950 | ATGGCTTGCCAAAACAATCTCGTTGTGAAGCAAATCATCGACTTGTACGACCAAATC<br>TCAAAGCTCAAGAGCTTAAAACCTTCCAAAAATGTCGACACTTTGTTCGGACAACTC<br>GTGTCCACGTGCTTACCCACGGATACAAACATCGATGTCACAAATATGTGTGAAGAA<br>GTCAAAGACATGAGAGCTAATCTCATCAAGCTTTGTGGTGAAGCCGAAGGTTATTTA<br>GAGCAACACTTCTCCACAATTTTGGGATCTTTACAAGAAGACCAAAACCCACTTGAC<br>CATTTACACATCTTTCCTTACTACTCCAACTACCTCAAGCTAGGCAAGCTCGAGTTCG<br>ATCTCCTGAGCCAACACTCAAGCCATGTCCCCACCAAGATTGCCTTCGTGGGTTCG<br>GGTCCGATGCCTCTCACATCCATCGTATTGGCCAAGTTTCACCTCCCCAACACGAC<br>GTTCCACAACTTTGACATCGACTCACACGCAAACACACTCGCTTCAAACCTCGTCTC<br>TCGCGACCCGGACCTCTCAAAACGCATGATCTTCCACACAACGGACGTACTAAACG<br>CAACCGAAGGCCTTGACCAATATGACGTCGTTTTCTTAGCGGCGCTTGTAGGGATG<br>GACAAAGAGTCAAAGGTCAAAGCCATCGAGCACTTGGAGAAACACATGGCTCCTGG<br>AGCTGTTCTTATGCTAAGGAGTGCTCATGCTCTCAGAGCTTTCTTATATCCAATCGTT<br>GACTCGTCTGATCTCAAAGGCTTTCAACTCTTGACCATCTATCATCCAACCGATGAC<br>GTGGTTAACTCGGTTGTGATCGCACGTAAGCTCGGTGGTCCGACCACGCCCGGGG<br>TTAATGGTACTCGTGGATGCATGTTTATGCCTTGTAACTGCTCCAAGATTCACGCGAT<br>CATGAACAACCGTGGTAAGAAGAATATGATCGAGGAGTTTAGTGCCATCGAGTAA |
| AT5G56<br>080 | ATGGCTTGCGAAAACAACCTCGTTGTGAAGCAAATCATGGACTTATACAACCAAATC<br>TCAAACCTCGAGAGCTTAAAACCATCCAAGAATGTCGACACTTTGTTCAGACAACTT<br>GTGTCCACGTGCTTACCAACGGACACGAACATCGATGTCACAGAGATACACGATGA<br>AAAAGTCAAAGACATGAGATCTCATCTCATCAAGCTTTGTGGTGAAGCCGAAGGTTA<br>TTTAGAGCAACACTTTTCAGCAATCTTAGGCTCTTTTGAAGACAACCCTCTAAACCAT<br>TTACACATCTTCCCCTATTACAACAACTATCTCAAACTAGGCAAACTCGAATTCGATCT<br>CCTATCTCAACACACAACCCATGTCCCGACCAAAGTCGCCTTTATTGGTTCCGGTCC<br>GATGCCACTTACTTCCATCGTCTTGGCCAAGTTCCACCTCCCCAACACAACGTTCCA<br>CAACTTCGACATCGACTCACACGCCAACACACTCGCTTCAAACCTCGTTTCTCGTG<br>ATTCTGACCTTTCCAAACGCATGATTTTCCACACAACTGATGTATTAAACGCTAAGGA<br>GGGGTTAGACCAATACGATGTTGTTTTCTTGGCAGCTCTTGTTGGGATGGATAAAGA<br>GTCAAAGGTCAAAGCTATTGAGCATTTAGAGAAGCATATGGCTCCTGGAGCTGTGGT<br>GATGCTAAGAAGTGCTCATGGTCTTAGAGCTTTCTTGTATCCAATCGTTGACTCTTGT<br>GATCTTAAAGGGTTTGAGGTGTTAACCATTTATCATCCGTCTGACGACGTGGTTAATT<br>CGGTGGTCATCGCACGTAAGCTTGGTGGTTCAAATGGAGCTCGAGGCAGCCAGAT<br>CGGACGGTGTGTGGTTATGCCTTGTAATTGCTCTAAGGTCCACGCGATCTTGAACAA<br>TCGTGGTATGGAGAAGAATTTGATCGAGGAGTATAGTGCCATCGAGTAA |
| AT1G09<br>240 | ATGGGTTGCCAAGACGAACAATTGGTGCAAACAATATGCGATCTCTACGAAAAGATC<br>TCAAAGCTTGAGAGTCTAAAACCATCCGAAGATGTCAACATTCTCTTCAAGCAGCTC<br>GTTTCCACATGCATACCACCAAACCCTAACATCGATGTCACCAAGATGTGTGACAGA<br>GTCCAAGAGATTCGACTTAATCTCATCAAGATTTGTGGTCTAGCCGAAGGTCACTTA<br>GAAAACCATTTCTCTTCGATCTTGACCTCTTACCAAGACAACCCACTTCATCATTTAA<br>ACATTTTCCCTTATTACAACAACTATTTGAAACTCGGAAAGCTCGAGTTCGACCTCCT<br>CGAACAAAACCTAAATGGCTTTGTCCCAAAGAGTGTGGCTTTCATTGGATCTGGTCC<br>TCTTCCTCTCACTTCCATCGTTCTTGCTTCATTCCATCTCAAAGACACAATCTTTCAC<br>AACTTTGACATCGACCCATCAGCGAACTCACTCGCTTCTCTTCTGGTTTCCTCTGAT<br>CCAGACATCTCTCAACGCATGTTCTTCCACACCGTTGATATAATGGACGTGACAGAG<br>AGCTTAAAGAGCTTTGATGTCGTGTTTCTAGCTGCTCTTGTTGGAATGAACAAAGAG<br>GAGAAAGTTAAAGTGATCGAGCATCTGCAGAAACACATGGCTCCTGGTGCTGTGCT<br>CATGCTTAGGAGTGCTCATGGTCCGAGAGCGTTTCTTTATCCGATCGTTGAGCCGTG<br>TGATCTTCAGGGGTTCGAGGTTTTGTCTATTTATCACCCAACAGATGATGTTATCAAC |

| | |
|---|---|
| | TCCGTGGTGATCTCTAAAAAGCATCCAGTTGTTTCAATTGGGAATGTTGGTGGTCCTAATTCATGCTTGCTCAAGCCTTGCAACTGTTCCAAGACCCACGCGAAAATGAACAAGAACATGATGATCGAGGAGTTCGGAGCTAGGGAGGAACAGTTGTCTTAA |
| AT1G56430 | ATGGGTTATTGCCAAGACGACCAACTCGTAAACAAGATCTGCGATCTTTACGAAAAGATCTCGAAGCTTGAGACCCTAAAGCCTTGTGAAGATGTCGACACTCTCTTCAAGCAGCTCGTGTCCACATGCATACCACCAAACCCTAACATCGACGTCACCAAGATGTCTGAAAACATCCAAGAGATGAGATCAAACCTCATCAAAATCTGTGGTGAGGCTGAAGGTTACTTAGAGCATCACTTCTCTTCAATCTTAACCTCTTTTGAAGATAACCCCCTTCATCATTTGAATCTTTTTCCTTACTACAACAACTACCTCAAACTAAGCAAGCTCGAGTTTGATCTCCTCGAACAGAACCTAAACGGTTTTGTTCCAAGGACTGTAGCTTTCATTGGCTCTGGTCCTCTCCCTCTTACTTCCGTCGTTCTTGCCTCTTCCCATCTCAAAGACTCGATCTTTCATAACTTTGACATCGACCCATCAGCGAATATGGTAGCAGCTCGTTTGGTTTCGTCTGATCCTGATCTTTCTCAACGTATGTTTTTCCATACTGTTGATATAATGGATGTAACCGAGAGCTTGAAGGGCTTCGACGTTGTGTTCTTGGCTGCTCTTGTAGGGATGGATAAAAAGGAGAAGGTTAAGGTGGTCGAGCATCTTGAGAAACACATGTCTCCTGGTGCTTTGCTCATGCTGAGAAGCGCTCATGGACCTAGAGCTTTTCTCTATCCAATCGTTGAGCCTTGTGATCTCGAAGGTTTCGAAGTTTTATCGGTTTATCACCCTACCGATGAAGTTATCAACTCCATTGTAATCTCAAGGAAGCTAGGTGAAGATGCTAATGGTGTTGTTCATGATCATATAGATCAAGCTTCGGATCTCGCGTGTAACTGTTCCAAGATCCACGTGATTATGAACAAGAAGAAGAGCATTATCGAGGAGTTCGCAGGTGCTAATGAAGAACAACTTACCTAG |
| HORVU.MOREX.r3.6HG0628880 | ATGGACGCCCAGAGCAAGGAGGTCGACGCCCTTGTCCAGAAGATCACCGGCCTCCACGCCGCCATCGCCAAGCTGCCCTCGCTCAGCCCGTCCCCGGACGTCGACGCGCTCTTCACCGACCTGGTCACCGCGTGCGTGCCCCCGAGCCCCGTGGACGTGACCAAGCTCGCCCCGGAGGCGCAGGCGATGCGGGAGGGCCTCATCCGCCTCTGCTCCGAGGCCGAGGGCAAGCTGGAGGCGCACTACTCCGACATGCTCGCCGCCTTCGACAACCCGCTCGACCACCTCGGCGTCTTCCCCTACTACAGCAACTACATCAACCTCAGCAAGCTCGAGTACGAGCTCCTCGCGCGCTACGTGCCCGGCGGCATCGCCCCGGCCCGCGTCGCCTTCATCGGCTCCGGCCCGCTGCCGTTCAGCTCCTACGTCCTCGCCGCGCGCCACCTGCCCGACACCGTGTTCGACAACTACGACCTGTGCGGCGCGGCCAACGACCGCGCGAGCAGGCTGTTCCGCGCGGACAAGGACGTCGGCGCCCGCATGTCGTTCCACACCGCCGACGTCGCGGACCTCACCGACGAGCTCGCTACGTACGACGTCGTCTTCCTGGCCGCGCTCGTTGGCATGGCCGCCGAGGACAAGGCCAAGGTGATCGCGCACCTTGGCGCGCACATGGCGGACGGGGCGGCCCTCGTCGTGCGCAGCGCGCACGGGGCGCGTGGGTTCCTCTACCCGATCGTCGATCCCCAGGACATTGGTCGAGGCGGGTTCGAGGTGCTCGCCGTGTGTCACCCCGACGACGACGTGGTGAACTCCGTCATCATCGCACAAAAGAGCAACGACGTGCACGAGTATGGACTTGGCAGCGGGCGTGGTGGACGGTACGCGCGAGGCACGGTGGTGCCGGTGGTCAGCCCACCCTGCAGGTTCGGCGAGATGGTGGCAGACGTGACCCAGAAGAGAGAGGAGTTTGCCAAGGCGGAAGTGGCCTTCTGA |
| HORVU.MOREX.r3.4HG0417690 | ATGGCTGCCCAGAACAACAACAAGGATGTCGCTGCCCTGGTGGAGAAGATCACCGGGCTCCACGCCGCCATCGCCAAGCTGCCGTCGCTCAGCCCATCCCCGGACGTCGACGCGCTCTTCACCGAGCTGGTCACGGCGTGCGTTCCCCCGAGCCCCGTGGACGTGACCAAGCTCGGCCCCGAGGCGCAGGAGATGCGGGAGGGCCTCATCCGCCTCTGCTCCGAGGCCGAGGGGAAGCTGGAGGCGCACTACTCCGACATGCTCGCCGCCTTCGACAACCCGCTGGATCACCTCGGCATCTTCCCCTACTACAGCAACTACATCAACCTCAGCAAGCTGGAGTACGAGCTCCTGGCGCGCTACGTGCCCGGCGGCATCGCCCCGGTCCGCGTCGCGTTCATCGGCTCCGGCCCGCTGCCGTTCAGCTCCTTTGTCCTGGCCGCGCGCCACCTGCCCGACACCATGTTTGACAACTACGACCTTTGCGGCGCGGCCAACGATCGCGCCAGCAAGCTCTTCCGCGCGGACACGGACGTGGGTGCCCGCATGTCGTTCCACACGGCCGACGTCGCGGACCTCGCCAGCGAGCTCGCCAAGTACGACGTCGTCTTCCTGGCCGCGCTCGTCGGCATGGCTGCCGAGGACAAGGCCAAGGTGATCGTGCACCTCGGCGCACACATGGCAGACGGGGCGGCCCTCGTCGTGCGCAG |

| | |
|---|---|
| | CGCACACGGAGCGCGCGGGTTCCTGTACCCGATTGTCGACCCCCAGGACATCGGCCGCGGCGGGTTCGAGGTGCTGGCCGTGTGCCACCCCGACGACGACGTGGTGAACTCCGTCATCATCGCACAGAAGTCCAAGGAGGTGCATGCCGATGGACTTGGCAGCGCGCGTGGTGCCGGTGGACAGTACGCGCGCGGCACGGTGCCGGTTGTCAGCCCCCCGTGCAGGTTCGGTGAGATGGTGGCGGACGTGACCCAGAACCACAAGAGAGACGAGTTTGCCAACGCCGAAGTGGCCTTTTGA |
| HORVU.MOREX.r3.2HG0099840 | ATGGAGGCCGAAAACGGCGAGGTGGCTGCTCTGGTCGAGAAGATCACCGGTCTCCACGCCGCCATCTCCAAGCTCCCGTCACTAAGCCCGTCTCCTCAAGTCGACGCGCTCTTCACCGAGCTGGTCGCGGCGTGCGTCCCATCAAGCCCGGTGGACGTGACCAAGCTCGGCCCGGAGGCGCAGGAGATGCGGCAGGACCTCATCCGCCTCTGCTCGGCCGCCGAGGGGCTGCTCGAGGCGCACTACTCCGACATGCTCACCGCGTTGGACAGCCCGCTCGACCACCTCGGCCGCTTCCCTTACTTTGACAACTACGTCAACCTCAGCAAGCTCGAGCACGATCTTCTGGCAGGTCACGTGGCGGCCCCGGCCCGCGTGGCGTTCATCGGGTCGGGGCCACTGCCGTTCAGCTCGCTCTTCCTCGCGACGTACCACCTGCGGGACACCCGGTTCGACAACTACGACCGGTGCAGCGTGGCGAACGGCCGGGCGATGAAGTTGGTCGGCGCGGCGGACGAGGGCGTGCGATCACGCATGGCGTTCCACACGGCCGAAGTCACGAACCTCACGGCTGAGCTCGGCACTTACGACGTGGTTTTCCTGGCCGCGCTCGTGGGAATGACGTCCGAGGAGAAGGCCGACGCCATAGCGCACTTGGGGAAGCACATGGCAGATGGGGCGGTGCTCGTGGCGCGAAGCGCGCACGGGGCGCGAGCGTTCCTGTATCCTGTAGTGGAGCTGGACGATGTCGGGCGTGGCGGGTTCCAAGTGCTGGCGGTGCACCACCCTGCAGGCGATGAGGTGTTCAACTCATTCATAGTTGCCCGGAAGGTGAAAATGAGTGCTTAA |
| LOC_Os03g19427 | ATGGAGGCTCAGAACCAAGAGGTCGCTGCCCTGGTCGAGAAGATCGCCGGCCTCCACGCCGCCATCTCCAAGCTGCCGTCGCTGAGCCCATCCGCCGAGGTGGACGCGCTCTTCACCGACCTCGTCACGGCGTGCGTCCCGGCGAGCCCCGTCGACGTGGCCAAGCTCGGCCCGGAGGCGCAGGCGATGCGGGAGGAGCTCATCCGCCTCTGCTCCGCCGCCGAGGGCCACCTCGAGGCGCACTACGCCGACATGCTCGCCGCCTTCGACAACCCGCTCGACCACCTCGCCCGCTTCCCGTACTACGGCAACTACGTCAACCTGAGCAAGCTGGAGTACGACCTCCTCGTCCGCTACGTCCCCGGCATTGCCCCCACCCGCGTCGCCTTCGTCGGGTCGGGCCCGCTGCCGTTCAGCTCCCTCGTGCTCGCCGCGCACCACCTGCCGGACGCGGTGTTCGACAACTACGACCGGTGCGGCGCGGCCAACGAGCGGGCGAGGAGGCTGTTCCGCGGCGCCGACGAGGGCCTCGGCGCGCGCATGGCGTTCCACACCGCCGACGTGGCGACCCTGACGGGGGAGCTCGGCGCGTACGACGTCGTGTTCCTGGCGGCGCTCGTGGGCATGGCGGCCGAGGAGAAGGCCGGGGTGATCGCGCACCTGGGCGCGCACATGGCGGACGGCGCGGCGCTCGTCGTGCGGAGCGCGCACGGGGCGCGCGGGTTCCTGTACCCGATCGTCGATCCCGAGGACGTCAGGCGTGGCGGGTTCGACGTTCTGGCGGTGTGCCACCCGGAGGACGAGGTGATCAACTCCGTCATCGTCGCCCGCAAGGTCGGTGCCGCCGCCGCCGCCGCCGCGGCGCGCAGAGACGAGCTCGCGGACTCGCGCGGCGTGGTTCTGCCGGTGGTCGGGCCGCCGTCCACGTGCTGCAAGGTGGAGGCGAGCGCGGTTGAGAAGGCAGAAGAGTTTGCCGCCAACAAGGAGCTGTCCGTCTAA |
| LOC_Os03g19420 | ATGGAGGCTCAGAACCAAGAGGTCGCTGCCCTGGTCGAGAAGATCGCCGGCCTCCACGCCGCCATCTCCAAGCTGCCGTCGCTGAGCCCATCCGCCGAGGTGGACGCGCTCTTCACCGACCTCGTCACGGCGTGCGTCCCGGCGAGCCCCGTCGACGTGGCCAAGCTCGGCCCGGAGGCGCAGGCGATGCGGGAGGAGCTCATCCGCCTCTGCTCCGCCGCCGAGGGCCACCTCGAGGCGCACTACGCCGACATGCTCGCCGCCTTCGACAACCCGCTCGACCACCTCGCCCGCTTCCCGTACTACGGCAACTACGTCAACCTGAGCAAGCTGGAGTACGACCTCCTCGTCCGCTACGTCCCCGGCATTGCCCCCACCCGCGTCGCCTTCGTCGGGTCGGGCCCGCTGCCGTTCAGCTCCCTCGTGCTCGCCGCGCACCACCTGCCGGACGCGGTGTTCGACAACTACGACCGGTGCGGCGCGGCCAACGAGCGGGCGAGGAGGCTGTTCCGCGGCGCCGACGAGGGCCTCGGCGCGCGCATGGCGTTCCACACCGCCGACGTGGCGACCCTGACGGGGGAGCTCGGCGCGTACGACGT |

| | |
|---|---|
| | CGTGTTCCTGGCGGCGCTCGTGGGCATGGCGGCCGAGGAGAAGGCCGGGGTGAT<br>CGCGCACCTGGGCGCGCACATGGCGGACGGCGCGGCGCTCGTCGTGCGGAGCG<br>CGCACGGGGCGCGCGGGTTCCTGTACCCGATCGTCGATCTCGAGGACATCCGGCG<br>GGGCGGGTTCGACGTGCTGGCCGTGTACCACCCCGACGACGAGGTGATCAACTCC<br>GTCATCGTCGCTCGCAAGGCCGACCCGCGTCGCGGCGGCGGGCTCGCCGGCGCA<br>CGCGGCGCGGTTCCAGTGGTGAGCCCGCCGTGCAAGTGCTGCAAGATGGAGGCG<br>GCCGCCGGCGCGTTCCAGAAGGCGGAAGAGTTCGCCGCCAAGAGGCTATCCGTCT<br>GA |
| LOC_Os07g48980 | ATGACGGTGGAAGTGGAGGCGGTGACCATGGCGAAGGAGGAGCAGCCGGAGGAG<br>GAGGAGGTGATCGAGAAGTTGGTTGAGAAGATCACCGGGCTGGCGGCGGCCATCG<br>GCAAGCTGCCGTCGCTGAGCCCGTCGCCGGAGGTGAACGCGCTGTTCACGGAGC<br>TGGTGATGACCTGCATCCCGCCCAGCAGCGTGGACGTGGAGCAGCTGGGGGCGG<br>AGGCGCAGGACATGCGCGGCCGCCTCATCCGCCTCTGCGCCGACGCCGAGGGCC<br>ACCTCGAGGCGCACTACTCCGACGTCCTCGCCGCCCACGACAACCCGCTCGACCA<br>CCTCGCCCTCTTCCCCTACTTCAACAACTACATCCAGCTCGCCCAGCTCGAGTACG<br>CCCTCCTCGCCCGCCACCTCCCCGCCGCCCCGCCGCCCTCCCGCCTCGCCTTCC<br>TCGGCTCCGGCCCGCTCCCGCTCAGCTCCCTCGTCCTCGCCGCCCGCCACCTCC<br>CCGCCGCCTCCTTCCACAACTACGACATCTGCGCCGACGCCAACCGCCGCGCCAG<br>CCGCCTCGTCCGCGCCGACCGCGACCTGTCCGCGCGCATGGCCTTCCACACCTC<br>CGACGTCGCCCACGTCACCACCGACCTCGCCGCCTACGACGTCGTGTTCCTGGCG<br>GCGCTGGTGGGTATGGCCGCCGAGGAGAAGGCGCGCATGGTGGAGCACCTCGGG<br>AAGCACATGGCGCCCGGCGCCGCCCTGGTGGTGCGGAGCGCGCACGGCGCCCG<br>GGGATTCCTGTACCCCGTGGTGGACCCGGAGGAGATCCGCCGCGGCGGCTTCGA<br>CGTGCTCGCCGTGCACCACCCGGAGGGCGAGGTGATCAACTCCGTCATCATCGCG<br>CGCAAGCCGCCCGTGGCGGCGCCGGCGTTGGAGGGAGGAGACGCGCACGCGCA<br>CGGCCATGGCGCCGTGGTGAGCCGTCCATGCCAGCGCTGCGAGATGGAGGCGAG<br>GGCGCACCAGAAGATGGAGGACATGTCCGCCATGGAGAAGCTGCCCTCCTCGTAG |
| TraesCS2A02G033500 | ATGGAGGCCGAAAACAGCGAGGTGGCTGCTCTGGTCGAGAAGATCACCGGCTTCC<br>ACGCCGCCATCTCCAAGCTCCCGTCGCTAAGCCCGTCCCCTCAAGTCGACGCGCT<br>CTTCACCGAGCTGGTCGCGGCGTGCGTCCCGTCGAGCCCGGTGGACGTGACCAA<br>GCTCGGCCCGGAGGCGCAGGAGATGCGGCAGGACCTCATCCGCCTGTGCTCAAC<br>CGCCGAGGGGCTGCTCGAGGCGCACTACTCCGACATGCTCACCGCCTTGGACAGC<br>CCGCTCGACCACCTCGGCCGCTTCCCTTACTTTGACAACTACATCAACCTGAGCAA<br>GCTCGAGAACGACCTTCTGGCCGGTCACATGGCGGCTCCGGCCCGCGTGGCGTT<br>CATCGGGTCAGGGCCGCTGCCGTTCAGCTCGCTCTTCCTCGCGACATACCACCTG<br>CCGGACACCCGCTTCGACAACTACGACCGGTGCAGCGTGGCCAATGGCCGGGCG<br>ATGAAGCTGGTCGGCGCGGCGGATGTGGACGTGCGCTCGCGCATGCCGTTCCACA<br>CGGCCGAAGTCGCGGACCTCACGTCTGAGCTCGGCGCGTACGACGTGGTTTTCCT<br>GGCGGCGCTCGTGGGGATGACGTCCGAGGAGAAGGCCAACACCATCGCGCACTT<br>GGGGAAGCACATGGCAGATGGGGCGGTGCTCGTCGCGCGAAGCGCGCACGGGG<br>CGCGAGCGTTCCTCTATCCTGTAGTGGAGCTGGACGATATCGGGCGTGGCGGGTT<br>CCAAGTGCTGGCCGTGCACCACCCTGCAGGTGATGAGGTGTTCAACTCATTCATTG<br>TTGCACAGAAGGTGAAGATATGA |
| TraesCS2B02G047100 | ATGGAGGCCGAAAACAGCGAGGTGGCTGCTCTGGTCGAGAAGATCACCGGCCTCC<br>ACGCCGCCATCTCCAAGCTCGCGTCTCTAAGCCCGTCCCCCGAAGTCGACGCGCT<br>CTTCACGGAGCTGGTCGCGGCGTGCGTCCCGTCGAGCCCGGTGGACGTGACCAA<br>GCTCGGCCCGGAGGCGCAGGAGATGCGGCAGGACCTCATCCGCCTCTGCTCGGC<br>CGCCGAGGGGCTGCTCGAGGCGCACTACTCCAACATGCTCACCGCCTTGGACAAC<br>CCGCTCGACCACCTCGGCCGCTTCCCTTACTTCGACAACTACATCAACCTGAGCAA<br>GCTCGAGCACGATCTTCTTGCCGGTCACGTGGCGGCCCCGGCCCGCGTGGCGTT<br>CATCGGGTCGGGGCCGCTGCCGTTCAGCTCCCTCTTCCTCGCGATGTACCACCTG<br>CCGGACACCCGGTTCGACAACTACGACCGGTGCAGCGTGGCCAATGGCCGTGCGA |

| | |
|---|---|
| | TGAAGCTGGTCGGCGCGGCGGACGAGGGCGTGCGTGCGCGCATGGCGTTCCACA<br>CGGCCGAAGTCGCGGACCTCACGGCTGAGCTCGGCGCGTATGATGTGGTCTTCCT<br>GGCGGCGCTCGTGGGAATGACGTCCGAGGAGAAGGCCAACACCATCGCGCACTT<br>GGGGAAGCACATGGCAGATGGCGCGGTGCTCGTCGCGCGAAGCGCCCACGGTGC<br>GCGAGCGTTCCTGTATCCTGTAGTGGAGCTGGACGATATCGGGCGTGGCGGGTTC<br>CAAGTGCTGGCTGTGCACCATCCTGCGGGCGATGAGGTGTTCAACTCATTTATTGTT<br>GCGCGGAAGGAGCATATTTCTACCGTGGGCACTATAGGGTGCTCATGGCAGCACGA<br>ATGTATATGGACTACAATTGAGGGTGTGGCAGACGGAGAGATGTCATTCAAGTAA |
| TraesCS6A02G163100 | ATGGATGCCCAGAAGATGGAGGTCGCTGCTCTGATCGAGAAGATCGCCGGTCTCCA<br>GGCCGCCATCGCCGGGCTGCCGTCGCTGAGCCCGTCCCCTGAGGTCGACAGGCT<br>CTTCACCGACCTCGTCACCGCGTGCGTCCCGCCGAGCCCCGTCGACGTGACGAA<br>GCTCAGCCCGGAGCACCAGAGGATGCGGGAGGCGCTCATCCGCCTCTGCTCCGC<br>CGCCGAGGGGAAGCTCGAGGCGCACTACGCCGACCTGCTCGCCACCTTCGACAA<br>CCCGCTCGACCACCTCGGCCGCTTCCCCTACTACAGCAACTACGTCAACCTCAGCA<br>GGCTGGAGTACGAGCTGCTGGCGCGCCACGTGCCGGGCATCGCGCCGGCGCGC<br>GTCGCCTTCGTCGGCTCCGGCCCGCTGCCGTTCAGCTCGTTCGTCCTCGCCGCGC<br>ACCACCTGCCCGACGCGCAGTTCGACAACTACGACCTGTGCGGCGCGGCCAACGA<br>GCGCGCCAGGAAGCTGTTCGGCGCGAGGGAGGACGGCGTGGGCGCGCGCATGA<br>AGTTCCACACGGCGGACGTCGCCGACCTCACGCAGGAGCTCGGTGCGTACGACG<br>TGGTCTTCCTCGCCGCGCTCGTCGGCATGGCGGCCGAGGAGAAGGCCAAGGTGAT<br>AGCCCACCTGGGCGCGCACATGGTGGAGGGGGCGTCCCTGGTCGTGCGGAGCGC<br>GCACGGCGCTCGCGGCTTCCTGTACCCCATCGTCGACCCGGAGGACATCAGGCGG<br>GGCGGGTTCGAGGTGCTGGCCGTGCACCACCCCGAAGGTGAGGTGATCAACTCT<br>GTCATCGTCGCCCGTAAGGCCGTCGACGCGCAGCTCAGTGGGCCGCAGAACGGA<br>GCACGGGGCGCGGTGCCGCTGGTCAGCCCGCCATGCAGCTTCTCCACCAAGATG<br>GAGGCGAGCGCGCTTGAGAAGAGCGAAGAGTTGGCCACCAAAGAGCTGGCCTTTT<br>GA |
| TraesCS6D02G148200 | ATGGATGCCCAGAACAACGAGGTCGCTGCTCTGATCGAGAAGATCGCCGGTCTCCA<br>GGCCGCCATCGCTGAGCTGCCGTCGCTGAGCCCGTCCCCCGAGGTCGACAGGCT<br>CTTCACCGCCCTCGTCACGGCCTGCGTCCCGCCGAGCCCCGTCGACGTGACGAA<br>GCTCAGCCCGGAGCACCAGAGGATGCGGGAGGCGCTCATCCGCCTCTGCTCCGC<br>CGCCGAGGGGAAGCTCGAGGCGCACTACGCCGACCTGCTCGCCACCTTCGACAA<br>CCCGCTCGACCACCTCGGCCGCTTCCCCTACTACAGCAACTATGTCAACCTCAGCA<br>GGCTGGAGTACGAGCTCCTGGCGCGCCACGTGCCGGGCATCGCGCCGGCGCGCG<br>TCGCCTTCGTCGGCTCCGGCCCGCTGCCGTTCAGCTCGTTCGTCCTCGCCGCGCA<br>CCACCTGCCCGACGCCCAGTTCGACAACTACGACCTGTGCGGCGCGGCCAACGA<br>GCGCGCCAGGAAGCTGTTCGGCGCGAGCGAGGACGGCGTGGGCGCGCGCATGA<br>AGTTCCACACGGCGGACGTCGCCGACCTCACGCAGGAGCTCGGCGCGTACGACG<br>TGGTCTTCCTCGCCGCGCTCGTCGGCATGGCGGCAGAGGAGAAGGCCAAGGTGAT<br>AGCCCACCTGGGCGCGCACATGGTGGAGGGGGCGTCCCTGGTCGTGCGGAGCGC<br>GCACGGCGCCCGCGGCTTCCTGTACCCCATCGTCGACCCGGAGGACATCAGGCG<br>GGGCAGGTTCGAGGTGCTGGCCGTGCACCACCCCGAAGGTGAGGTGATCAACTCT<br>GTCATCGTCGCCCGTAAGGTCGTCGACGCGAAGCTCAGTGGGCCGCAGAACGGAG<br>ACGCGCACGCACGGGGCGCGGTGCCGCTGGTCAGCCCGCCATGCAGCTTCTCCA<br>CCAAGATGGAGGCGGGCGCGCTTGAGAAGAGCGAAGAGCTGGCCGCCAAAGAGC<br>TGGCCTTTTGA |
| TraesCS6D02G148600 | ATGGATGCCCAGAACAAGGAGGTCGCTGCTCTGATCGAGAAGATCGCCGGTCTCC<br>AGTCCGCCATCGCCGAGCTGCCGTCGCTGAGCCCGTCCCCCGAGGTCGACAGGC<br>TCTTCACCGACCTCGTCACGGCCTGCGTCCCGCCGAGCCCCGTCGACGTGACGAA<br>GCTCAGCCCGGAGCACCAGAGGATGCGGGAGGCGCTCATCCGCCTCTGCTCCGC<br>CGCCGAGGGGAAGCTCGAGGCGCACTACGCCGACCTGCTCGCCACCTTCGACAA<br>CCCGCTCGACCACCTCGCCCGCTTCCCCTACTACAGCAACTACGTCAACCTCAGCA |

| | |
|---|---|
| | GGCTGGAGTACGAGCTCCTGGCGCGCCACGTGCCGGGCATCGCGCCGGCGCGCG<br>TCGCCTTCATCGGCTCCGGCCCGCTGCCGTTCAGCTCGTTCGTCCTCGCCGCGCA<br>CCACCTGCCCGACGCGCACTTCGACAACTACGACCTGTGCGGCGCGGCCAACGA<br>GCGCGCCAGGAAGCTGTTCGGCGCGAGCGAGGACGGCGTGGGCGCGCGCATGA<br>AGTTCCACACGGCGGACGTCGCCGACCTCTCGCAGGAGCTCGGCGCGTACGACG<br>TGGTCTTCCTCGCCGCGCTCGTCGGCATGGTGGCCGAGGAGAAGGCCAAGGTGAT<br>AGCCCACCTGGGCGCGCACATGGTGGAGGGGGCGTCCCTGGTCGTGCGGAGCGC<br>GCACGGCGCCCGCGGCTTCCTGTACCCCATCGTCGACCCGGAGGACATCAGGCGA<br>GGCGGGTTCGAGGTGCTGGCCGTGCACCACCCCGAAGGTGAGGTCATCAACTCTG<br>TCATTGTCGCCCGTAAGGTCGTCGACGCGCAGCTCAGTGGGCCGCTGAACGGAGA<br>CGCGCACGCGCGGGGCGCGGTGCCGCTGGTCAGCCCGCCGTGCAGCTTCTCCAC<br>CAAGATGGAGGCGGGCGCGCTTGAGAAGAGCGACGAGTTGGCCACCAAAGAGCT<br>GGCCTTTTGA |
| TraesC<br>S2A02<br>G04990<br>0 | ATGGCTCTCCAGAACAAGGAGGTGGATGCCCTGGTCCAGAAGATCACCGGACTCC<br>ACGCCGCCATCGCCAAGCTGCCGTCGCTCAGCCCGTCCCCGGACGTAGACGCGCT<br>CTTCACCGAGCTGGTCACCGCGTGCGTTCCCCCGAGCCCCGTGGACGTGACCAAG<br>CTCGGCCCGGAGGCGCAGGAGATGCGGGAGGGCCTCATCCGCCTCTGCTCTGAG<br>GCCGAGGGGAAGCTGGAGGCGCACTACTCCGACATGCTCGCGGCCTTCGACAACC<br>CGCTCGACCACCTCGGTATGTTCCCCTACTACAACAACTACATCAACCTCAGCAAGC<br>TTGAATACGAGCTCCTGGCGCGCTACGTGCCCGGCGGCATCGCCCCGGCCCGCGT<br>CGCCTTCATCGGCTCCGGCCCACTGCCGTTCAGCTCTTTCGTCCTCGCCGCACGC<br>CACTTGCCCGACACCATGTTCGACAACTACGACCTATGTGGCGCGGCCAATGACCG<br>CGCCAGCAAGCTGTTCCGCGCGGACAAGGACGTGGGCGCCCGCATGTCGTTCCA<br>CACGGCCGACGTCGCGGACCTTGCCGGCGAGCTCGCTAAGTACGACGTCGTCTTC<br>CTGGCCGCGCTCGTGGGCATGGCCGCAGAGGACAAGGCCAAGGTGATCGCGCAC<br>CTCGGCGCACACATGGCAGACGGGGCGGCCCTCGTTGTACGCAGTGCGCACGGG<br>GCACGCGGGTTCCTGTACCCGATCGTAGACCCCCAGGACATCGCCGGAGGCGGGT<br>TCAAGGTGCTCGCCGTGTGCCACCCCGACGACGACGTGGTGAACTCCGTTATCATC<br>GCACAGAAGTCCAAGGACTTGCATGCCAATGGACATCACCGTGGGCATGGTGGAC<br>AGTGCGCGCATGGCACGGTGCCGGTGGTCAACCCACCGTGCAGGTTTGGTGAGAT<br>GGTGACAGACATGGCCCAGAAGAGAGAGGAGTTTGCCAACGCCGAAGTGGTCTTT<br>TGA |
| TraesC<br>S2B02<br>G06080<br>0 | ATGGCTGCCCAGAACAAGGAGGTGGATGCCCTGGTCCAGAAGATCACCGGACTCC<br>ACGCCGCCATCGCCAAGCTGCCGTCGCTCAGCCCGTCCCCGGACGTCGACGCGC<br>TCTTCACCGAGCTGGTCACCGCGTGCGTTCCTCCGAGCCCTGTGGACGTGACCAA<br>GCTCGGCCCGGAGGCGCAGGAGATGCGGGAGGGCCTCATCCGCCTCTGCTCCGA<br>GGCCGAGGGAAAGCTGGAGGCGCACTACTCCGACATGCTCGCTGCCTTTGACAAC<br>CCACTCGACCACCTAGGTATGTTCCCCTACTACAACAACTACATCAACCTCAGCAAG<br>CTTGAGTACGAGCTCCTGGCGCGCTATGTGCCCGGTGGCATCGCCCCGGCCCGGG<br>TCGCCTTCATCGGCTCCGGCCCACTGCCGTTCAGCTCTTTCGTCCTCGCCGCGCG<br>CCACTTGCCCGACACCATGTTCGACAACTATGACCTATGTGGGGCGGCCAACGACC<br>GCGCCAGCAAGCTGTTCCGCGCGGACAAGGACGTGGGCGCCCGCATGTCGTTCC<br>ACACGGCCGATGTAGCGGACCTCGCTGCTGAGCTCGCTACATACGACGTCGTCTTC<br>CTGGCTGCGCTCGTGGGCATGGCCGCGGAGGACAAGGCCAAGGTGATCGCGCAC<br>CTCGGTGCACACATGGCAGACGGTGCGGCCCTTGTTGTACGCAGTGCACACGGGG<br>CACGCGGGTTCCTGTACCCGATCGTCGACCCCCAGGACATCGCAGGAGGCGGGTT<br>CGAGGTGCTGGCCGTGTGCCACCCCGATGACGACGTGGTGAACTCCGTCATCATC<br>GCACAGAAGTCCAAGGACGTGCTTGCCAATGGACTTCGCCGCGGGCATGGTGGAC<br>AGTACGCGTGCGGCACGGTGCCGGTGGTCAGCCCACCGTGCAGGTTCGGTGAGA<br>TGGTGACGGACGTGACCCACAAGAGAGAGGAGTTTGCCAACGCCGAAGTGGTCTT<br>TTGA |

| | |
|---|---|
| TraesCS5A02G552400 | ATGGCTGCCCAGAACAACAAGGAGGTGGATGCCCTGGTGGAGAAGATCACCGGGCTCCACGCCGCCATCGCCAAGCTGCCGTCGCTCAGCCCTTCCCCGGCCGTCGACCTGCTCTTCACCGAGCTGGTCACGGCGTGCGTTCCCCCGAGCCCCGTGGACGTGACCAAGCTCGGCCCGGAGGCGCAGGAGATGCGGGAGGGCCTCATCCGCCTCTGCTCCGAGGCCGAGGGGAAGCTGGAGGCGCACTACTCCGACATGCTCGCCGCCTTCGACAACCCTCTGGACCACCTCGGCATGTTCCCCTACTACAGCAACTACATCAACCTCAGCAAGCTGGAGTACGAGCTCCTGGCCCGCTACGTGCCTGGTGGCATCGCCCCTGCCCGCGTCGCCTTCATCGGCTCCGGCCCGCTGCCGTTCAGCTCCTTCGTCCTCGCCGCGCGCCACCTGCCCGACACCATGTTCGACAACTACGACCTGTGCGGCGCGGCCAATGACCGCGCCAGCAAGCTGTTCCGCGCCGACAAGGACATGGGCGCCCGCATGTCGTTCCACACGGCCGACGTAGCGGACCTCGCTGGCGAGCTCGCCAAGTACGACGTGGTCTTCCTGGCCGCACTTGTTGGCATGGCGGCTGAGGACAAGGCCAAGGTGATAGCACACCTCGGCACACACATGGCAGACGGGGCGGCCCTCGTCGTGCGCAGCGCACACAGGGCACGCGGGTTCCTGTACCCGATCGTCGACCCCCAGGACATCACCCTAGGTGGGTTCAAGGTGTTGGCCGTGTGCCATCCAGACGACGACGTGGTGAACTCCGTCATCATCGCACAGAAGTCCAAGGACGTGCATGTTAGTGGACTTCGCAGCGGGCCTGCTGTGGGTGGACAGTCTGCTCGCTCCACGGTGCCGGTGGTCAGCCCGCCGTGCAGGTTCGGTGAGATGGTGGCGGATATGACACAGAAGAGAGAGGAGTTTGCCAACGCCGAAGTGGCCTTTTGA |
| TraesCSU02G125200 | ATGGCTGCCCAGAACAACAAGGAGGTGGATGCCCTGGTGGAGAAGATCACCGGCCTCCACGCCGCCATCGCCAAGCTGCCGTCGCTCAACCCATCCCCGGACGTCGACGCGCTCTTCACTGAGCTGGTCACGGCGTGCGTTCCCCCGAGCCCGGTGGACGTGACCAAGCTCGGCCCGGGGGCGCAGGAGATGCGGGAGGGCCTCATCCGCCTTTGCTCCGAGGCCGAGGGGAAGCTGGAGGCGCACTACTCCGACATGCTTGCCGCCTTCGACAACCCTCTGGATCACCTCGGCATGTTCCCCTACTACAGCAACTACATCAACCTCAGCAAGCTTGAGTACGAGCTCCTGGCGCGCTACGTGCCCGGTGGCATCGCCCCTGCCCGCGTCGCCTTCATCGGCTCCGGCCCACTCCCGTTCAGCTCCTTTGTCCTGGCCGCGCGCCATCTGCCCGACACCATGTTCGACAACTACGACCTGTGCGGTGCGGCCAACGACCGTGCCAGCAAGCTGTTCCGTGCGGACACGGACGTGGGCGCCCGCATGTCGTTCCACACGGCCGACGTAGCGGACCTCGCCGGCGAGCTCGCCAAGTACGACGTGGTCTTCCTGGCCGCACTTGTTGGCATGGCCGCCGAGGACAAGGCGAAGGTGATCGCACACCTCGGCGCACATATGGCGGACGGTGCGGCTCTCGTCGTGCGCAGCGCTCACGGGGCACGCGGGTTCCTGTACCCGATCGTCGACCCCCAGGACATCACCCTAGGCGGGTTCGAGGTGCTGGCCGTGTGCCACCCAGACGACGACGTGGTGAACTCCGTCATCATCGCACAGAAGTCCAAGGACGTGCATGTGAGTGGACTTCACAGTGGGCGTGCTGTGGGTGGACAGTCTGCTCGCGGCACGGTGCCGGTGGTCAGCCCGCCGTGCAGGTTCGGTGAGATGGTGGCGGAGGTGACGCAGAAGAGAGAGGAGTTCGCCAACGCCGAAGTGGCCTTCTGA |
| TraesCS3B02G068500 | ATGTTTGACAACTACGACCTGTGCGGCGCGGCCAACGAGCGCGCCAGCAAGCTGTTCCGCGCGGACACGGACGTGGGCGCCCGCATGTCGTTCCACACGGCCGACGTCGCGGACCTCGCCGGCGAGCTTGCCAAGTACGACGTCGTCTTCCTGGCGGCGCTCGTGGGCATGGCCGCCGAGGAAAAGGCCAGGGTGATCGCGCACCTTGGCACGCACATGGCAGACGGGGCGGCCCTCGTCGTGCGCAGCGCGCACGGCGCACGCGGGTTCCTATACCCGATCGTGGACCCCCAGGACATCGGCCGAGGCGGGTTTGAGGTGCTGGCCGTGTGCCACCCCGACGACGACGTGGTGAACTCCGTCATTATCGCACACAAGTCGAAGGACATGCATGCCAGTGGACTTCGCAGCGAGCGTGCCGGTGGTGGGCAGTATGCGCGTGGCACGGTGCCGGTGGTCAGCCCGCCGTGCAGGTTCGGCAAGATGGTGGCCGACGTGAACCAGAAGAGAGAGGAGTTTGCCAAGGCCGAAGTGGCTTTTTGA |
| TraesCS4B02G183900 | ATGGATGCCCAGAACAAGGAGGTCGACGCCCTGGTCCACAAGATCACCGGCCTCCACGCCGCCATCGCCAAGCTGCCGTCCCTCAGCCCATCCCCCGACGTCGACGCGCTCTTCACCGACCTGGTCACCGCGTGCGTCCCCCCGAGCCCCGTGGACGTGACCAAGCTCGGCCCGGAGGCGCAGGAGATGCGGGAGGGCCTCATCCGCCTCTGCTCCGA |

| | |
|---|---|
| | GGCCGAGGGGAAGCTGGAGGCGCACTACTCCGACATGCTCGCCGCCTTCGACAACCCGCTCCACCACCTCGCCATCTTCCCCTACTACAGCAACTACATCAACCTCAGCAAGCTGGAGTACGAGCTCCTGGCGCGCTACGTGCCCGGCGGCATCGCCCCGGCCCGCGTCGCGTTCATCGGCTCCGGCCCGCTGCCGTTCAGCTCCTACGTCCTCGCCGCCCGCCACCTGCCCGACACCATGTTTGACAACTACGACCTGTGTGGCGCGGCCAACGACCGTGCGAGCAAGCTGTTCCGCGCGGACAAGGAAGTGGGCGCCCGCATGTCGTTCCACACCGCTGACGTCGCGGACCTTGCCGGCGAGCTCGCCGCGTACGACGTCGTCTTCCTGGCCGCGCTCGTGGGCATGGCCGCCGAGGACAAGGCCAAGGTGATCGCACACCTCGGCGCGCACATGGCGGACGGGTCGGCCCTCGTCGTGCGCAGCGCGCACGGGGCGCGTGGGTTCCTGTACCCGATCGTCGATCCCCAGGACATCGGCCGAGGCGGGTTTGAGGTGCTGGCCGTGTGTCACCCCGACGACGACGTGGTGAACTCCGTCATCATCGCACAGAAGTCCAAGGACATGCATGCCAATGAACACCGCAACGGGCGTGGTGGACAGTACGCGCGGGGCACGGTGCCGGTGGTGAGCCCGCCGTGCAGGTTCGGCGAGATGGTGGCGGACGTGACCCAAAAGAGAGAGGAGTTCGCCAATGCCGAAGTGGCCTTCTGA |
| TraesCS4D02G184900 | ATGCACAGCTTTGCCAGCTCCATCGATGAGTGCTCTAGACGGCAGGCAGCTACATATACCCCGTGCTCCTCGTTCCATAGCTCACCAAGCAGCCCGATCCACCAACTCCACTCCACTCCTGTGCCTCAGAGTTCATCAAGTACTCGTCAGGTACCAGGTAAAATGGATGCCCAGAACAAGGAGGTCGCCGCCCTGGTCCACAAGATCACCGGCCTCCACGCCGCCATCGCCAAGCTGCCGTCCCTCAGCCCATCCCCCGACGTCGACGCGCTCTTCACCGACCTGGTCACCGCGTGCGTCCCCCCGAGCCCCGTGGACGTGACCAAGCTCGGCCCGGAGGCGCAGGAGATGCGGGAGGGCCTCATCCGCCTCTGCTCCGAGGCCGAGGGGAAGCTGGAGGCGCACTACTCCGACATGCTCGCCGCCTTCGACAACCCGCTCGACCACCTCGGCATCTTCCCCTACTACAGCAACTACATCAACCTCAGCAAGCTGGAGTACGAGCTCCTGGCGCGCTACGTGCCCGGCGGCATCGCCCCGGCCCGCGTCGCCTTCATCGGCTCCGGCCCGCTGCCGTTCAGCTCCTACGTCCTCGCCGCCCGCCACCTGCCCGACACCATGTTCGACAACTACGACCTGTGCGGCGCGGCCAACGACCGCGCGAGCAAGCTGTTCCGCGCCGACAAGGACGTGGGCGCCCGCATGTCGTTCCACACCGCCGACGTCGCGGACCTCGCCGGCGAGCTCGCCGCGTACGACGTCGTCTTCCTGGCCGCGCTCGTGGGCATGGCTGCCGAGGACAAGGCCAAGGTGATCGCGCACCTCGGCGCGCACATGGCGGACGGGGCGGCCCTCGTCGTGCGCAGCGCGCACGGCGCGCGTGGGTTCCTGTACCCGATCGTCGATCCCCAGGACATCGGCCGAGGCGGGTTCGAGGTGCTGGCCGTGTGTCACCCCGACGACGACGTGGTGAACTCCGTCATCATCGCACAGAAGTCCAAGGACATGCATGCCAATGAACATCGCAACGGGCGTGGTGGACAGTACGCGCGGGGCACGGTGCCGGTGGTGAGCCCACCGTGCAGGTTCGGCGAGATGGTGGCGGACGCGGCCCAGAAGAGAGAGGAGTTCGCCAACGCTGAAGTGGCCTTCTGA |
| TraesCS6A02G093000 | ATGGATGCCCAGAACAAGGAGGTCGATGCCCTGGTCCAGAAGATCACCGGCCTCCACGCCGCCATCGCCAAGCTGCCCTCGCTCAGCCCGTCCCCCGACGTCGACGCGCTCTTCACCGACCTGGTCACCGCGTGCGTCCCCCCGAGCCCCGTGGACGTGACCAAGCTCGGCCCGGAGGCACAGGCGATGCGGGAGGGCCTCATCCGCCTCTGCTCCGAGGCCGAGGGGAAGCTGGAGGCGCACTACTCCGACATGCTCGCCGCCTTCGACAACCCGCTCGACCACCTCGGCGTCTTCCCCTACTACAGCAACTACATCAACCTCAGCAAGCTGGAGTACGAGCTCCTGGCGCGCTACGTCCCCGGCGGCATCGCCCCGGCCCGTGTCGCCTTCATCGGCTCCGGCCCGCTGCCGTTCAGCTCGTACGTCCTCGCCGCACGCCACCTGCCCGACGCGGTGTTCGACAACTACGACCTGTGTGGCGCGGCCAACGACCGTGCGAGCAAGCTGTTCCGCGCCGACAAGGACGTGGGCGCACGCATGTCATTCCACACCGCCGACGTAGCGGACCTCACCGACGAGCTCGCCACGTACGACGTCGTCTTCCTGGCCGCGCTCGTGGGCATGGCTGCCGAGGACAAGGCCAAGGTGATCGCACACCTTGGCGCGCACATGGCGGACGGGGCGGCCCTCGTCGTGCGGAGCGCGCACGGGGCGCGTGGGTTCCTGTACCCGATCGTCGATCCCCAGGACATCGGCCGAGGCGGGTTCGAGGTGCTCGCCGTGTGTCACCCCGACGACGACGTGGTGAACTCTGTCA |

| | |
|---|---|
| | TCATCGCGCAGAAGTCCAGCGACATGCATGCGAATGGATTTCGCAACGGGCGTGGT<br>GGACAGTGCGCGCGGGGCACGGTGCCGGTGGTCAGCCCACCCTGCAGGTTCGGC<br>GAGATGGTGGCGGACGTGAGCCAGAAGAGAGAGGAGTTTGCCAACGCGGAAGTG<br>GCCTTCTAA |
| TraesC<br>S6A02<br>G38620<br>0 | ATGGACGCCCAGAACAAGGAGGTGGACGCCCTGGTCCAGAAGATCACCGGCCTCC<br>ACGCCGCCATCGCCAAGCTGCCCTCGCTCAGCCCGTGCCCCGCCGTCGACGCGC<br>TCTTCACCGACCTCGTCACCGCGTGCGTCCCCCCGAGCCCCGTGGACGTGACCGA<br>GCTCGGCCCGGAGGCGCAGGCGATGCGCGAGGGCCTCATCCGCCTCTGCTCCGA<br>GGCCGAGGGGAAGCTGGAGGCGCACTACTCCGACATGCTCGCCGCCTTCGACAAC<br>CCGCTCGACCACCTCGGCGTCTTCCCTTACTACAGCAACTACATCAACCTCAGCAA<br>GCTGGAGTACGAGCTCCTGTCGCGCTACGTGCCCGGCGGCATCGCCCCGGCCCG<br>TGTCGCCTTCATCGGCTCCGGCCCGCTGCCGTTCAGCTCCTACGTCCTCGCCGCG<br>CGCCACCTGCCCGACACGGTGTTCGACAACTACGACCTGTGCGGCGCGGCCAAC<br>GCCCGCGCGAGCAGGCTGTTCCGCGCGGACAGGGACGTGGGCGCCCGCATGTC<br>GTTCCACACCGCCGACGTCGCGGACCTCACCGACGTGCTCGCTACGTATGATGTC<br>GTCTTCCTGGCCGCGCTCGTGGGCATGGCGGCCGAGGACAAGGCCAAGGTGATC<br>GCGCACCTTGGCGCGCACATGGCGGACGGGGCGGCCCTCGTCGTGCGCAGCGC<br>GCACGGGGCGCGTGGGTTCCTGTACCCGATCGTCGATCCCCAGGACATCGGCCGA<br>GGCGGGTTCGAGGTGCTCGCCGTGTGTCACCCAGACGACGACGTGGTGAACTCC<br>GTCATCATCGCGCAAAAGACCAACGACGTGCATGAGTATGGACTTGGCAACGGGCG<br>TGGTGGACGGTACGCGCGGGGCACGGTGCCGGTGGTCAGCCCGCCCTGCAGGTT<br>CGGCGAGATGGTGGCGGACGTGACCCAGAAGAGAGAGGAGTTTGCCAACGCGGA<br>ACTGGCCTTCTGA |
| TraesC<br>S6D02<br>G37080<br>0 | ATGGACGCCCAGAACAAGGAGGTCGACGCCCTGGTCCAGAAGATCACCAGGCTCC<br>ACGCCGCCATCTCCAAGCTGCCCTCGCTCAGCCCGTCCCCCGACGTCGACGCGCT<br>CTTCACCGACCTCGTCACCGCGTGCGTCCCCGCGAGCGCCGTGGACGTGACCAA<br>GCTCGGCCCGGAGGCGCAGGCGATGCGGGAGGGCCTCATCCGCCTCTGCTCCGA<br>GGCCGAGGGAAAGCTGGAGGCGCACTACTCCGACATGCTCGCCGCCTTCGACAAC<br>CCGCTCGACCACCTCGGCGTCTTCCCCTACTACAGCAACTACATCAACCTCAGCAA<br>GCTGGAGTACGAGCTCCTCGCGCGCTACGTGCCCGGCGGCATCGCCCCGGCACG<br>TGTCGCGTTCATCGGCTCCGGCCCGCTGCCGTTCAGCTCCTACGTCCTCGCCGCG<br>CGCCACCTGCCCGACACGGTGTTCGACAACTACGACCTGTGCGGCGCGGCCAAC<br>GCCCGTGCGAGCAAGCTGTTCCGTGCGGACAGGGACGTGGGCGCCCGCATGTCG<br>TTCCACACCGCCGACGTCGCGGACCTCACCGACGTGCTCGCCACGTACGACGTCG<br>TCTTCCTGGCCGCGCTCGTGGGCATGGCTGCCGAGGACAAGGCCAAGGTGATCGC<br>ACACCTTGGCGCGCGCATGGCGGACGGGGCGGCCCTCGTCGTGCGCAGCGCGCA<br>CGGGGCGCGTGGGTTCCTGTACCCGATCGTCGATCCCCAGGACATCGGCCGAGGC<br>GGGTTCGAGGTGCTCGCCGTGTGTCACCCCGACGACGACGTGGTGAACTCCGTCA<br>TCATCGCACAAAAGTCCAACGACATGCATGCGAATGGACTTCGCAACGGGCGTGGT<br>GGACAGTACGCGCGGGGCACGGTGCCGGTGGTCAGCCCACCCTGCAGGTTCGGC<br>GAGATGGTGGCAGACGTGGCCCAGAAGAGAGAGGAGTTTGCCAACGCGGAAGTG<br>GCCTTCTGA |
| TraesC<br>S2A02<br>G09570<br>0 | ATGGGCGTGGAGGGCTGCTGCAACAAGAAGGTGATGGAGGAGGAGGCCCTGGTG<br>AAGAAGATCACCGGCCTCGCCGCGGCCATCGGCGAGCTGCCCTCGCTGAGCCCCT<br>CGCCGGCGGTGAACGCGCTCTTCACCGAGCTGGTGACGTCCTGCATCCCGGCGA<br>GCACCGTCGACGTCGACGCGCTGGGCCCGGAGGCGCAGGAGATGCGCGCCCGC<br>CTCATCCGCCTCTGCGCCGACGCCGAGGGCCACCTGGAGGCGCACTACTCCGACC<br>TCCTCGCCGCGCACGACAACCCGCTCGACCACCTCACCCTCTTCCCCTACTTCAAC<br>AACTACATCAAGCTCAGCCAGCTGGAGCACGGCCTCCTCGCCCGCCACGTCCCGG<br>GCCCGGCCCCGGCCCGCGTCGCCTTCCTCGGCTCCGGCCCGCTGCCGCTCAGCT<br>CCCTCGTGCTCGCCGCGCGCCACCTGCCCGACGCCTCCTTCGACAACTACGACAT<br>CTCCGGCGAGGCCAACGAGCGCGCCAGCCGGCTGGTCCGCAACGACGCCGACG |

| | |
|---|---|
| | CCGGCGCGCGCATGGCGTTCCGCACGGCAGACGTGGCCGACGTGACGACCGAGC<br>TGGCGGGCTACGACGTGGTGTTCCTGGCGGCGCTGGTGGGCATGGCGGCCGAGG<br>AGAAGGCGCGCCTGGTGGAGCACCTCGGGAGGCACATGGCCCCCGGCGCGGCG<br>CTCGTGGTGCGGAGCGCGCACGGCGCGCGCGGGTTCCTGTACCCGGTGGTGGAC<br>CCCGAGGAGATCCGGCGAGGCGGGTTCGAGGTGCTCACCGTGCACCACCCGGAG<br>GACGAGGTGATCAACTCCGTCATCATCGCGCGCAAGGCGGCGGCGCCAGCAGCC<br>GTCGCCGCCGACGGCGACGTGCCGGTGAACAACGGCGGCATGCCGGCCCAGTGC<br>GCGGTGGCGGTGAGCAGACCGTGCCTGGGCTGCGCCTGCGAGCTGGGGACGAG<br>GGCGCACCAGAAGATGAAGGAGATGGCCATGGAGGAGATGGAGGCGTAG |
| TraesC<br>S2B02<br>G11110<br>0 | ATGGGCATGGAAGGCTGCTGCAACAAGAAGGTGATGGAGGAGGAGGCCCTGGTGA<br>AGAAGATCACCGGCCTCGCCGCGGCCATCGGCGAGCTGCCCTCGCTGACCCCGT<br>CGCCGGCGGTGAACGCGCTCTTCACCGAGCTGGTGACGTCCTGCATCCCGGCGA<br>GCACCGTCGACGTCGACGCGCTGGGCCCGGAGGCGCAGGAGATGCGCGCCCGC<br>CTGATCCGCCTCTGCGCCGACGCCGAGGGCCACCTGGAGGCGCACTACTCCGAC<br>CTCCTCGCCGCGCACGACAACCCGCTCGACCACCTCACCCTCTTCCCCTACTTCAA<br>CAACTACATCAAGCTCAGCCAGCTGGAGCACGGCCTCCTCGCCCGCCACGTCCCG<br>GGCCCTGCCCCGGCCCGCGTCGCCTTCCTCGGCTCCGGCCCGCTGCCGCTCAGC<br>TCCCTCGTGCTCGCCGCGCGCCACCTCCCCGACGCCTCCTTCGACAACTACGACA<br>TCTCCGGCGAGGCCAACGAGCGCGCCAGCCGGCTGGTCCGCAACGACGCCGACG<br>CCGGCGCGCGCATGGCGTTCCGCACGGCGGACGTGGCCGACGTGACCACCGAGC<br>TGGCGGGGTACGACGTGGTGTTCCTGGCGGCGCTGGTGGGCATGGCGGCCGAGG<br>AGAAGGCGCGCCTGGTGGAGCACCTCGGGAGGCACATGGCCCCCGGCGCGGCG<br>CTCGTGGTGCGGAGCGCGCACGGCGCGCGCGGGTTCCTGTACCCGGTGGTGAAC<br>CCCGAGGAGATCCGGCGAGGCGGGTTCGAGGTGCTCACCGTGCACCACCCGGAG<br>GACGAGGTGATCAACTCCGTCATCATCGCGCGCAAGGCGGCCGCGCCAGCAGCC<br>GTTGCCGCCGACGGCGACGTGCCGGTGAACAGCGGCAGCATGCCGGCCCAGTGC<br>GCGGTGGCGGTGAGCAGACCGTGCCTGGGCTGCGCCTGCGAGCTGGGGGCGAG<br>GGCGCACCAGAAGATGAAGGAGATGGCCATGGAGGAGATGGAGGCGTAG |
| TraesC<br>S2D02<br>G09420<br>0 | ATGGGCATGGAGGGGTGCTGCGACAAGAAGGTGATGGAGGAGGAGGCCCTGGTG<br>AAGAAGATCACCGGCCTCGCCGCGGCCATCGGCGAGCTGCCCTCGCTGAGCCCG<br>TCGCCGGCGGTGAACGCGCTCTTCACCGAGCTGGTGACGTCCTGCATCCCGGCGA<br>GCACCGTGGACGTGGACGCGCTGGGCCCGGAGGCGCAGGAGATGCGCGCCCGC<br>CTCATCCGCCTCTGCGCCGACGCCGAGGGCCACCTGGAGGCGCACTACTCCGACC<br>TCCTCGCCGCGCACGACAACCCGCTCGACCACCTCACCCTCTTCCCCTACTTCAAC<br>AACTACATCAAGCTCAGCCAGCTGGAGCACGGCCTCCTCGCCCGCCACGTCCCGG<br>GCCCCGCCCCGGCCCGCGTCGCCTTCCTCGGCTCCGGCCCGCTGCCGCTCAGCT<br>CCCTCGTGCTCGCGGCGCGCCACCTCCCCGACGCCTCCTTCGACAACTACGACAT<br>CTCCGGCGAGGCCAACGAGCGCGCCAGCCGGCTCGTGCGCAACGACGCCGACG<br>CCGGCGCGCGCATGGCGTTCCGCACCGCCGACGTGGCCGACGTGACGACCGAGC<br>TGGCGGGGTACGACGTGGTGTTCCTGGCGGCGCTGGTGGGCATGGCCGCCGAGG<br>AGAAGGCGCGCCTGGTGGAGCACCTCGGGAGGCACATGGCCCCCGGCGCGGCG<br>CTCGTGGTGCGGAGCGCGCACGGCGCGCGCGGCTTCCTGTACCCGGTGGTGGAC<br>CCCGAGGAGATCCGGCGAGGCGGGTTCGAGGTGCTCACCGTGCACCACCCGGAG<br>GACGAGGTGATCAACTCCGTCATCATCGCGCGCAAGGCGGCGGCGCCGGCGGCC<br>GTTGCCGCCGACGGCGACGTGCCGGTGAACAACGGCGGCATGCCGGCGCAGTGC<br>GCGGTGGCGGTGAGCAGACCGTGCCTCGGCTGCGCCTGCGAGCTGGGGACGAG<br>GGCGCACCAGAAGATGAAGGAGATGGCCATGGAGGAGATGGAGGCGTAG |
| Zm0000<br>1eb014<br>680 | ATGGAGGCCCAGAACGTGGAGGTTGCTGCCCTGGTGAAGAAGATCGCGGACCTCC<br>ACGCCGACATCACCAAGCTGCCGTCGCTGAGCCCGTCCCCCGACGTCAACGCGCT<br>ATTCACCAGCCTCGTCATGGCCTGCGTCCCGCCGAGCACTGTCGACGTGACCAAG<br>CTCAGCCCGGACTCCCAGAGGATGCGGGAGGAGCTCATTCGCCTCTGCTCCGACG<br>CCGAGGGCCACCTCGAGGCGCACTACGCCGACATGCTGGCCGCCTTCGACAACCC |

| | |
|---|---|
| | GCTCGACCATCTCGGCCGCTTCCCATACTTTAGCAACTACATCAACCTGAGCAAGCT<br>GGAGTACGACCTCCTGGTCCGCTACATCCCGGGTCTCGCGCCTTCCCGCGTCGCC<br>TTCGTCGGGTCAGGCCCCTTGCCGTTCAGCTCGCTCGTGCTCGCCGCCCGCCACC<br>TGCCCAACACGACATTCGACAACTATGACCGGTGCGCTGCCGCCAACGACCGTGC<br>GCGGAAGCTGGTCCGCGCGGACAAGGACCTGAACGCGCGCATGTCCTTCCACAC<br>GGTGGACGTAGCCAACCTGACGGATGACCTCGGCAAGTACGACGTCGTCTTCCTT<br>GCCGCGCTCGTCGGCATGGCGGCCGAGGACAAGGCCAAGGTGGTCGTGCACCTT<br>GGCAGGCACATGGCGGACGGAGCGGCTCTCGTCGTGCGGAGCGCGCACGGGGC<br>TCGCGGGTTCCTATACCCGATCGTGGATCCCGAGGACATCCGTCGTGGCGGGTTTG<br>ACGTTCTGACGGTGTACCACCCGGACGATGAGGTGATCAACTCTGTCATCATCGCG<br>CGCAAGATCGACGCCCATGCAAACACGGAGGTTTCTGCCCTGGTGCAGAAGATAAC<br>CGGCCTCCACGCCGCCATCAACAAGCTGCCATCGCTGAGCCCATCCCCCGACGTC<br>GACGCGCTGTTCACCGAGCTCGTCATGGCCTGCGTCCCACCGAGCCCCGTCGACG<br>TGACCAAGCTCGGCACGGACGCGCAGAGGATGCGGGAGGAGCTCATCCGCCTCT<br>GCTCCGACGCCGAGGGCCACCTCGAGGCGCACTATGCCGACATGCTCGCCGCCTT<br>CGACAACCCGCTCGACCATCTCGGCCGCTTCCCTTACTTCAACAACTACGTCAACC<br>TGAGCAAGTTGGAGTACGACCTCCTGGTTCGCTATGTCCCGGGCATCGCACCTTCC<br>AGAATCGCCTTCGTCGGGTCCGGACCCCTGCCGTTCAGCTCGCTCGTGCTCGCCT<br>CCCGCCACCTGCCGAACGTGATGTTCGACAACTACGACCGGTGCGCCGCCGCCAA<br>CGACCGTGCACGGAAGCTGGTCCGCGCGGACGAGGGTCTGCGCAAGCGAATGTT<br>CTTCCACACCGCCGACGTCGCCAACCTCACGGACGAGCTCCGCAAGTACGACGTG<br>GTCTTCCTGGCCGCGCTCGTCGGCATGGCGGCCGAGGACAAGGCCAAGGTGGCT<br>ACGCACCTTGGCAGACACATGGCGGATGGAGCGGCTCTCATCGTGCGGAGCGCCC<br>ACGAGGCTCGAGGGTTCCTGTACCCGATCGTCGATCCCGAGGACATCCGTCGCAG<br>CGGCTTCGACGTGCTTGCCGTGTACCACCCGGACGACGAAGTCATTAACTCCGTCA<br>TCGTCGCGCGCAAGATCAACGCCCACGTGAAGGGGCTTCAGGATGGACACGCGCA<br>TGCGCGCGGTGTGGTGCCGATAGTGAGCCCGCCATGCAAGTGCTGCAAGATGGAG<br>GCGAACACGCTCCACCAAAAGAGGGAGGAGATGGCGACGGCGTAG |
| Zm0000<br>1eb396<br>250 | ATGGAGGCCCAGAACGTGGAGGTTGCTGCCCTGGTGCAGAAGATCGCGGCCCTCC<br>ACGCCGACATCGCCAAGCTGCCGTCGCTGAGCCCGTCCCCCGACGCCAACGCGC<br>TGTTCACCAGCCTCGTCATGGCCTGCGTCCCGCCAAACCCTGTCGACGTGACCAA<br>GCTCAGCCCGGACGTCCAGGGGATGCGAGAGGAGCTCATCCGCCTCTGCTCCGAC<br>GTCGAGGGCCACCTCGAGGCGCACTACGCCGACATGCTCGCCGCCTTCGACAACC<br>CGCTCGACCACCTCGGCCGCTTTCCCTACTTCAGCAACTACATCGACCTGAGCAAG<br>CTGGAGTTTGACCTCCTTGTCCGCTACATCCCAGGGCTCGCGCCTTCCCGTGTCGC<br>CTTCGTCGGCTCCGGCCCCTTGCCGTTCAGCTCGCTCGTGCTCGCCGCGCGCCAC<br>CTGCCCAACACACTGTTCGACAACTACGATCGGTGTGCCGCCGCCAACGACCGTG<br>CTCGGAAGCTGGTCCGTGCGGACAAGGACCTGAACGCGCGCATGTCCTTCCACAC<br>GGTCGACGTCGCCAACCTCACGGACGAGCTCGCCAAGTACGAC |
| Zm0000<br>1eb218<br>440 | ATGGTCGTCATGGGCAAGCACGATGACGATGAGAAGCAGCAGCAGCAGATGGAGA<br>TGGAAGACGTACATGAAGTAGGAGCCGTCGAGGTGCTGCCGCTGGAGAATGAGGC<br>GGAGGCCGCGCTGGTGCGCAAGATCTCCGGCCTGGCCGCCGCCATCGCGAAGCT<br>GCCGTCGCTGAGCCCGTCGCCGGAGGTGAACGCGCTGTTCACGGACCTCGTCAC<br>GGCCTGCATCCCGCGCAGCACCGTGGACGTGGAACGCCTGGGCCCGGAGCTGCA<br>GCGCATGCGCGCGGGCCTCATCCGCCGCTGCGCCGACGCCGAGGCCCAGCTGGA<br>GGCGCACTACTCGGACCTGCTGGCGGCCTTGGACAGCCCGCTGGACCACCTGCC<br>GCTCTTCCCCTACTTCGGCAACTACCTGCTGCTGGGCGAGCTGGAGCACGGCCTG<br>CTGGCGCGCCACGTCCCGGGCCCGCCGCCCGCCCGCGTCGCCTTCGTGGGGTC<br>CGGCCCGCTCCCGCTCAGCTCCCTCGTGCTCGCCGCGCGCCACCTCCCCGCCGC<br>CGCCTTCGACAACTACGACATCTGCGCGGAGGCCAACGGCCGGGCGCGCCGCCT<br>CGTGCGCGCCGACGCCGCGCTCGCCGCGCGCATGGCGTTCCGCACCTCCGACGT<br>GGCCCACGTCACGCGGGGCCTGGCCACCTACGACGTCGTCTTCCTGGCGGCGCT |

| | |
|---|---|
| | CGTGGGCGTGGCGGCCGAGGAGAAGGCGCGCGTGGTGGAGCACCTCGGGAGGC<br>ACATGGCGCCCGGCGCCGCGCTGGTGGTGCGCAGCGCGCACGGCGCGCGCGCC<br>TTCCTGTACCCCGTCGTCGACCCAGACGAGATCCGCCGCGGCGGCTTCGACGTGC<br>TCGCCGTGCACCACCCGGAGGGGGAGGTGATCAACTCCGTCATCGTCGCGCGTAA<br>GCCCGTCCCCGTCGCCGGAGTGGGGCACGCGCACGCGCACGCCCACGGGCACG<br>GCGCCGTGCTCAGCCGGCCCTGCTGCCTCTGCTGCGAGATGGAGGCCCGGACGC<br>AC |
